# Genetically encoded, noise-tolerant, auxin biosensors in yeast facilitate metabolic engineering and directed evolution

**DOI:** 10.1101/2023.03.21.533585

**Authors:** Patarasuda Chaisupa, Md Mahbubur Rahman, Sherry B. Hildreth, Saede Moseley, Chauncey Gatling, Richard F. Helm, R. Clay Wright

**Author notes:** These authors contributed equally to this work (Patarasuda Chaisupa, Md Mahbubur Rahman).

## Abstract

Auxins are important plant growth regulating compounds that are applied in vast quantities to crops across the globe to control weeds and improve crop quality and yield. Auxins are also produced by nearly every kingdom of life and control both organismal behavior as well as inter-kingdom interactions. Improving our understanding of auxin biosynthesis and signaling is critical to both improving crop plants and controlling symbiotic, commensal, and parasitic inter-kingdom relationships, many of which are critical to ecosystems from forests and oceans to the human microbiome. We present a suite of auxin biosensors that will advance our understanding of and ability to engineer auxin perception by plants and auxin production by fungi. We have developed genetically encoded, ratiometric auxin biosensors in the model yeast *Saccharomyces cerevisiae*, based on the mechanism plants use to perceive auxin. The ratiometric design of these biosensors improves measurements of auxin concentration by reducing clonal and growth phase variation. These biosensors are capable of measuring exogenous auxin in yeast cultures across five orders of magnitude, likely spanning the physiologically relevant range. We implement these biosensors to measure the production of auxin during different growth conditions and phases for *S. cerevisiae*. Finally, we demonstrate how these biosensors could be used to improve quantitative functional studies and directed evolution of plant auxin perception machinery. These genetically encoded auxin biosensors will enable future studies of auxin biosynthesis, transport, and signaling in a wide range of yeast species, as well as other fungi, and plants.

**Graphical Abstract:** 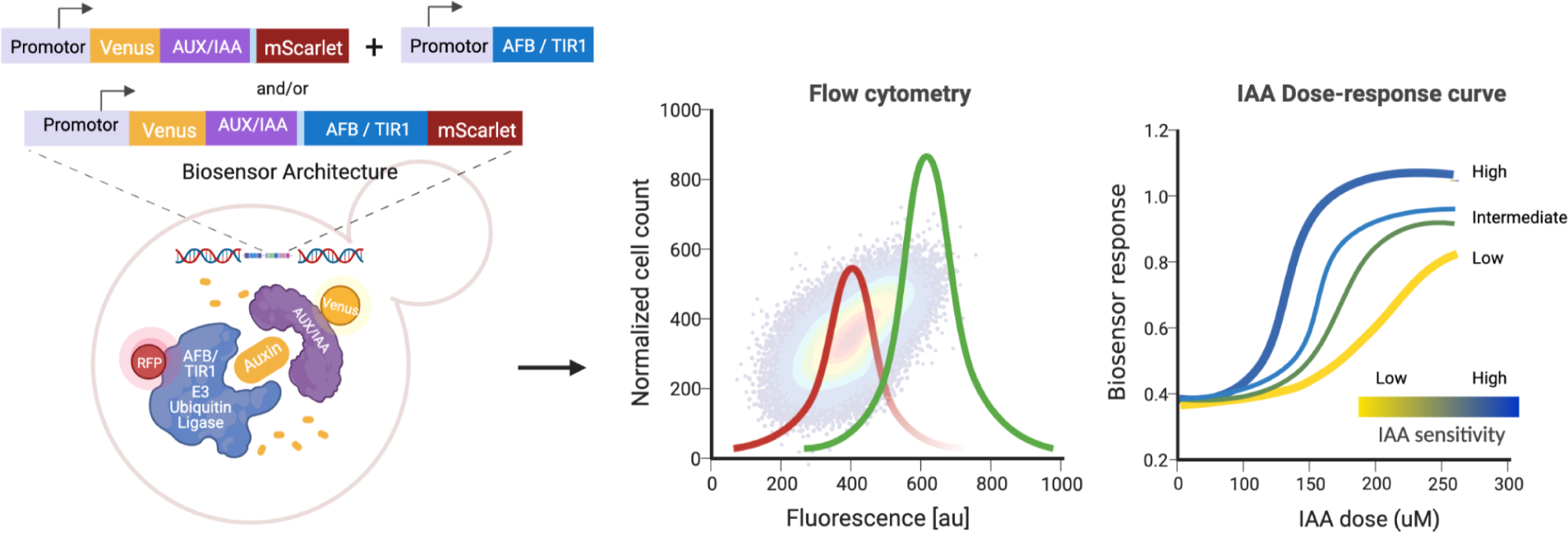

## Introduction

Auxins are important indole-derived signaling molecules that regulate the growth, metabolism, and behavior of plants, bacteria, fungi, and animals^1–6^. Auxins regulate nearly every aspect of plant growth and development^7,8^. Synthetic auxins are commonly used herbicides and plant growth regulators in agriculture^9–11^, and are regularly applied to over 200 million hectares of land each year (∼10% of the world’s total cropland)^12,13^.

Endogenous auxins, particularly the most abundant natural auxin, indole-3-acetic acid (IAA), regulate gene expression in plants, according to the canonical nuclear auxin signaling mechanism, by inducing proteasomal degradation of Aux/IAA repressive transcription factors^14,15^, which recruit TOPLESS co-repressors^16,17^ (Fig. 1). The TIR1/AFB auxin receptors^18,19^, which are part of Skp1-Cullin-F-box (SCF) ubiquitin ligase complexes, have a weak basal affinity for Aux/IAA proteins that is greatly enhanced in the presence of auxin ^20,21^. This auxin-mediated interaction with the SCF^TIR1/AFB^ ubiquitin ligase complex promotes Aux/IAA ubiquitination and proteasomal degradation. Aux/IAA degradation relieves repression of class A Auxin Response Factors (ARF), which then activate auxin-responsive gene expression^22–25^. Aux/IAA half-life varies from minutes to hours in plants^26^. The rate of Aux/IAA degradation varies across the Aux/IAA and TIR1/AFB auxin co-receptor families ^27^, and variation in TIR1/AFB or Aux/IAA sequence can alter Aux/IAA degradation rate and subsequent plant development^28–30^. As auxin signaling plays an important role in nearly every aspect of plant growth, development, and behavior, if we can parse how auxin signaling affects a particular trait, we can possibly alter the signaling machinery to engineer that trait^31^.

**Figure 1.**
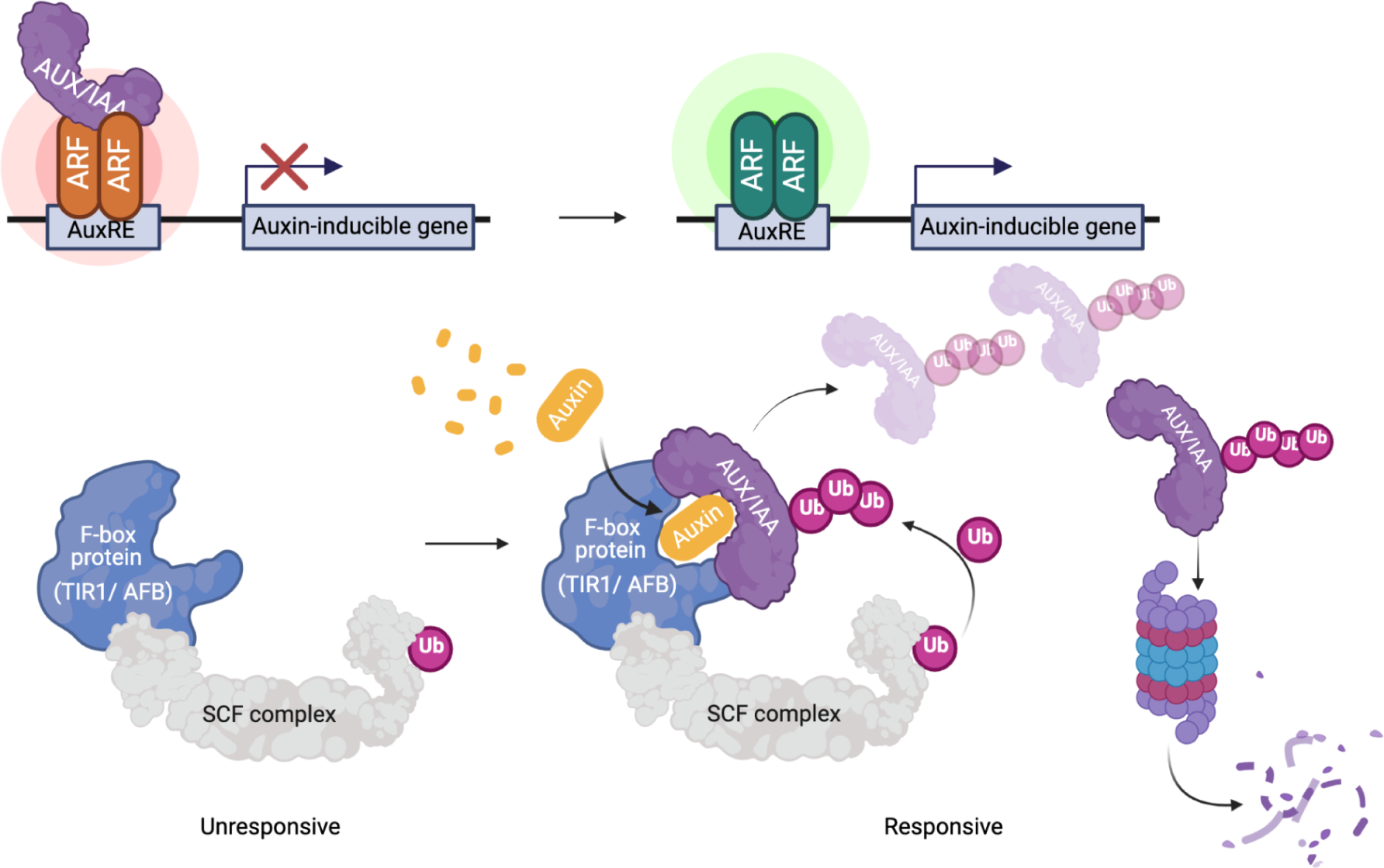
A simple model of the nuclear auxin signaling pathway of plants. Auxin related gene expression is naturally prevented by heterodimerization of transcriptional repressors Aux/IAA and the ARFs at the promoters of auxin-inducible genes containing auxin response elements (AuxREs). The presence of auxin bound to the leucine-rich repeat domain of TIR1/AFB SCF-type ubiquitin-protein ligase complexes promotes interaction of TIR1/AFB and Aux/IAA proteins. The ubiquitin-protein ligase complex consequently transfers activated ubiquitin to the Aux/IAA leading to polyubiquitination and proteasomal degradation. The ARF dimers are then released from the repression of the Aux/IAAs, to promote transcription from AuxRE-containing promoters.

Many microbes synthesize and perceive auxin, particularly indole-3-acetic acid (IAA), the most prevalent natural auxin, which may play some role in the ability of these microbes to colonize plants or animals^32–34^. Microbially produced auxin can mediate both beneficial and pathogenic microbe–plant interactions by modulating plant growth and microbiome composition, leading auxin to be called “a widespread physiological code” in interkingdom interactions^4^. Exogenous auxin increases pseudohyphae, required for adhesion and invasive growth in *Saccharomyces cerevisiae*^35,36^ and increased virulence of blast fungus^37^. Conversely, exogenous auxin treatment of barley can reduce Fusarium head blight severity and yield losses^38^. Both separate and simultaneous application of auxins and beneficial/biocontrol yeasts (which may or may not produce significant auxin) have been used successfully to combat pre-and post-harvest pathogens in a variety of fruits, often with simultaneous application providing synergistic effects^39–45^. Microbial production and perception of auxin is also known to be critical to plant interactions with rhizobacteria and mycorrhizal fungi^46–49^. Significant variation in auxin biosynthesis levels exists between strains of *S. cerevisiae*^50^, but no causal mechanism for this variation is known.

While there is growing evidence that auxin modulates microbiome composition and plant–microbe interactions (as reviewed by Kunkel and Johnson^51^) interkingdom signaling has rarely been studied systematically with dynamic resolution from both the plant and microbe perspective. While microbial auxin biosynthesis pathways have been established (as reviewed by Morffy and Strader^52^) most microbes have several redundant pathways, many enzymes remain unidentified, and little is known about how auxin production is regulated. As both pathogenic and beneficial microbes produce auxin, this leaves many open questions. What are the mechanisms by which microbes produce and secrete auxin, and how do they regulate auxin production in response to environmental cues? How do microbes use auxin to promote plant growth and colonization, and what are the molecular and cellular mechanisms underlying these effects? How does auxin production and perception differ between pathogenic and beneficial microbes? Improving our understanding how auxin affects beneficial versus pathogenic plant-microbe interactions, using auxin biosensors in microbes and plants, may provide new insights into sustainable agricultural practices.

The amount of auxin in cells or tissues is typically analyzed by conventional analytical methods such as HPLC, GC-MS, LC-MS, or enzyme-linked immunosorbent assays (ELISA)^53–57^. While these methods have high selectivity and sensitivity, they are destructive, invasive, time-consuming, and require laborious sample preparation. Also, measurements are generally limited to bulk averages of a sample of cells and require an individual sample for each time point. A sensitive bio-analytical device such as a genetically encoded biosensor (herein referred to as *biosensor*) that rapidly responds to user-defined molecules would allow sensing and coupling the molecules of interest to a more rapidly measurable output such as a fluorescence or luminescence^58,59^. Biosensors offer non-invasive, high-throughput, dynamic measurements with the potential for live cellular or subcellular resolution through imaging.

In addition to measuring the concentration of an analyte, biosensors can also be useful for studying and engineering gene function. For example, a biosensor constructed from a receptor for an analyte, can also be used to study the function of the receptor, *i.e.* perception of the analyte, via mutational scanning. Fluorescent biosensors in particular, can be paired with fluorescence-accelerated cell sorting (FACS) to enrich or isolate cells containing certain fluorescent properties, which due to the fluorescent biosensor are correlated with analyte concentrations and biosensor function. Used in this way, genetically encoded fluorescent biosensors facilitate deep mutational scanning^60^ and directed evolution^61^ of the genes comprising them, as well as associated native genes of the expression host. Deep mutational scanning pairs *en masse* functional analysis of gene variant libraries, for example by FACS, with deep sequencing of the functionally-sorted pools of variants to quantify gene variant function, based on enrichment in high-versus low-function pools^62^. Deep mutational scanning aims to map gene sequence–function relationships *en masse* to predict variant effects and inform genetic engineering efforts^63–65^. Directed evolution similarly explores gene sequence–function maps, but with the goal of pushing function in a certain direction through sequential rounds of diversification, expression, and functional selection^66,67^. As auxin signaling plays a role in nearly every aspect of plant growth, development, and behavior and has been exploited as the target of synthetic herbicides and plant growth regulators, improvements to our ability to study and engineer auxin perception can make dramatic impacts on plant science and agriculture^9,11,68–70^. Additionally, auxin-induced protein degradation is used in numerous eukaryotic organisms to control protein levels and advances in this technology have improved our understanding of essential protein dosage across many species^71–76^.

Several auxin signaling reporters and biosensors have been developed previously for use in plants and applied to study the capacity and dynamics of auxin signaling. Auxin-responsive transcriptional reporters such as DR5, DR5rev, GH3pro, SAURpro, and pIAAmotif have been successfully engineered, extensively used, and enabled great steps forward in our understanding of auxin signaling^22,77–79^. This type of reporter relies on the auxin-responsive promoter element (AuxRE) and the native plant auxin signal transduction cascade, which may vary with time and throughout development. Auxin biosensors commonly known as DII, fuse Aux/IAA degron sequences (originally identified as domain II) to a fluorescent protein^80^. In the presence of auxin, the fluorescent protein-Aux/IAA fusion forms a co-receptor complex with SCF^TIR1/AFB^ ubiquitin ligase, resulting in ubiquitination and degradation of the fluorescent protein-Aux/IAA fusion. Thus, for these biosensors, auxin signaling is inversely proportional to fluorescence. As these biosensors are only dependent on the native TIR1/AFB function, and not on transcription and translation of a reporter gene, their output provides a more immediate and reliable estimate of relative auxin concentrations within a tissue. However, there are many potential confounding factors in measurements of a fluorescent protein-Aux/IAA fusion. The total fluorescence intensity within a cell is proportional to the accumulation of the matured fluorescent protein which is the sum of the rate of translation and maturation, the rates of dilution by cell division and expansion, as well as the basal turnover rates and auxin-induced degradation. To control for such confounding factors and relativize auxin-induced degradation, quantitative ratiometric versions R2D2 and qDII have been developed using an free fluorescent protein with a separate emission spectrum expressed from the same promoter as the fluorescent protein-Aux/IAA fusion^78,81,82^. These biosensors and the commonly used DR5 reporter are sensitive to nanomolar levels of exogenous auxin and report changes in endogenous auxin and auxin sensitivity. Recently, a Förester resonance energy transfer (FRET) biosensor for auxin, called AuxSen, was engineered from a bacterial tryptophan-binding protein^83^. Through rounds of saturation mutagenesis in the tryptophan binding pocket, AuxSen specifically detects exogenous auxin in plant protoplasts from micromolar to millimolar range. These auxin sensors mentioned above have been developed for studying auxin in plants, and not fungi or other microbes, and have only very recently been preliminarily implemented in the plant pathogenic fungus *Magnaporthe oryzae*^84^.

Here, we have built upon previous auxin ratiometric biosensors in plants^80,82,78,81^ and our previous work in a synthetic recapitulation of auxin perception in yeast^27,30,85^ to build and characterize a series of auxin biosensors in yeast with the goals of measuring auxin and assessing sequence–function relationships in the TIR1/AFB–Aux/IAA auxin co-receptors. We aimed to build biosensors that allow easy, flexible manipulation of the *TIR1/AFB* and *Aux/IAA* coding sequences and are capable of measuring expression level and functional changes in both of these genes, as well as endogenous auxin production by yeast. We test and characterize four biosensor designs, constructed from the *Aux/IAA17* coding sequence fused to Venus, co-expressed with *mScarlet-I* as a polycistronic fusion via viral 2A ribosomal-skipping/self-cleaving sequences^86^, with the coding sequence of *Arabidopsis thaliana TIR1* or *AFB2* fused to *mScarlet-I* or expressed *in trans* from a separate promoter. By fusing *TIR1* or *AFB2* to *mScarlet-I* we aim to control for receptor accumulation in our measurements of auxin and auxin perception. We show that these ratiometric designs decrease cell-to-cell and clonal variation likely resulting from expression and metabolic-state variation. We demonstrate that these biosensors can measure exogenous auxin from low nanomolar to high micromolar levels, covering the likely range of endogenous auxin production levels by yeast. Finally, we demonstrate the potential of these biosensors to report functional changes in a library of *AFB2* variants.

## Results and Discussion

### Rational design and engineering of bicistronic ratiometric indole-3-acetic acid sensor circuits in *S. cerevisiae*

We aimed to design a suite of genetically encoded biosensors in yeast capable of both detecting a wide range of auxin concentrations as well reporting on the molecular function of sequence variants in the genes comprising the biosensor circuit (Fig 2). To do this, we built upon previous recapitulation of auxin perception by the plant TIR1/AFB receptor and Aux/IAA co-receptor via heterologous expression in yeast^27^. The formation of this complex promotes poly-ubiquitination of the Aux/IAA co-receptor, leading to its degradation by the proteasome. The rate of degradation of Aux/IAA-fluorescent-protein fusion can be measured by time-course flow cytometry^85^. This degradation rate depends on the *Aux/IAA* and *TIR1/AFB* isoforms expressed as well as auxin concentration^20,26,27,68,87,88^. The *Arabidopsis thaliana TIR1* or *AFB2*, and *Aux/IAA17* were chosen for auxin receptors and co-receptor, respectively, in our auxin biosensors, as Aux/IAA17 is rapidly degraded in the presence of AFB2 but much more slowly in the presence of TIR1^27^. We hypothesized that this difference in degradation rates would allow the resulting biosensors to collectively measure a wide range of auxin concentrations. Here, we chose to fuse the coding sequence of *Venus* to the 5’ of the *Aux/IAA* coding sequence, based on the rapid maturation time, high quantum yield, and alignment with our flow-cytometer optics.

**Figure 2.**
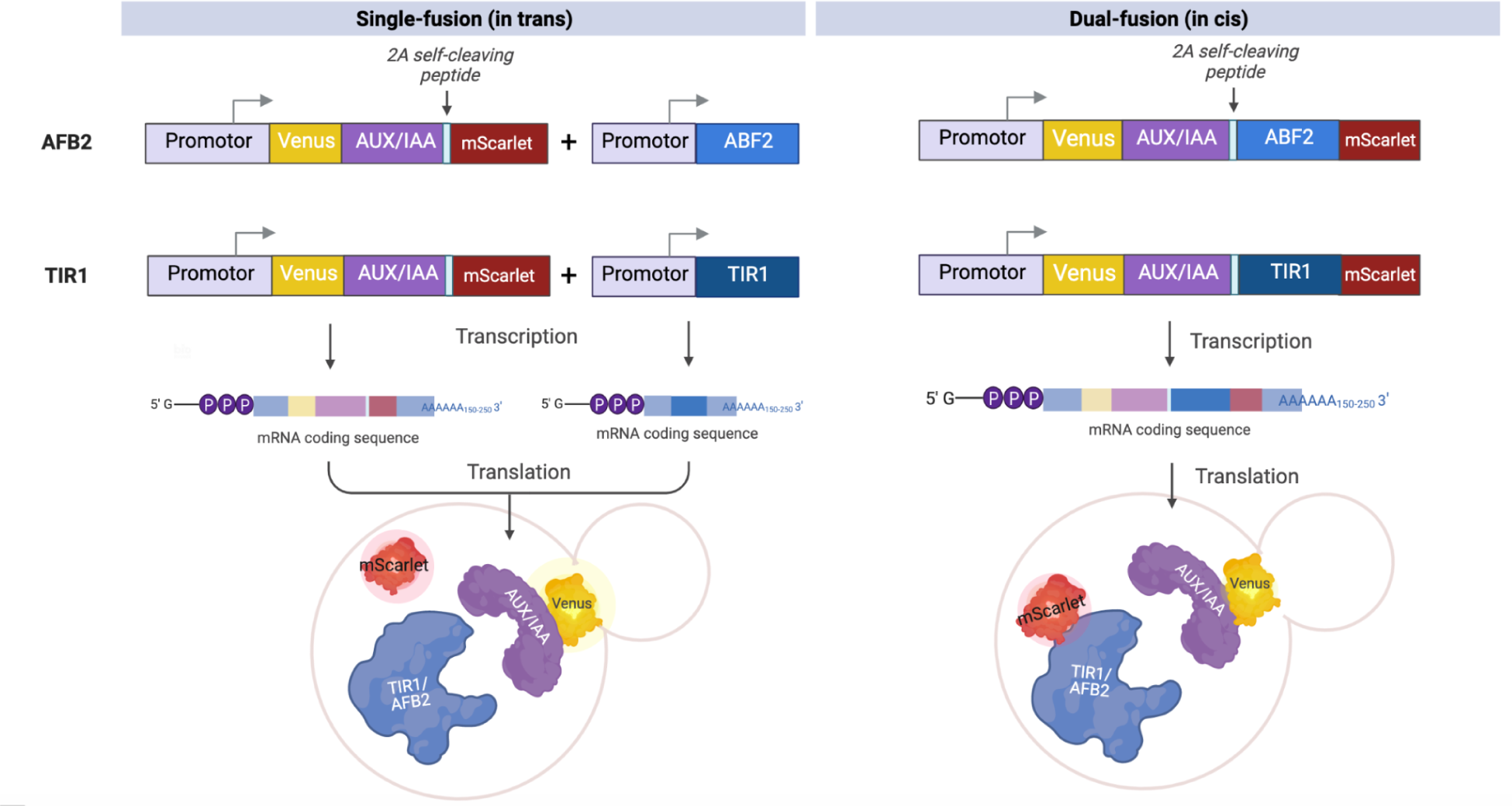
Construct design and protein schematics of the engineered auxin biosensors. The single-fusion designs consist of a Venus:IAA17 fusion and free mScarlet-I expressed from the same cistron. The bicistronic Equine Rhinitis B Virus (ERBV) 2A self-cleaving peptide is inserted into the cassettes. The auxin receptor, TIR or AFB2, is expressed separately from another construct. The dual-fusion designs consist of a Venus-Aux/IAA17 fusion and TIR1-or AFB2-mScarlet-I fusion expressed from a single mRNA via ERBV-2A, offering fewer steps of genetic manipulation and cell transformation.

In this and previous synthetic recapitulations of auxin perception in yeast, we observed high cell-to-cell variation in fluorescent-protein-Aux/IAA accumulation (Fig 3A), as well as clonal variation and fluctuations due to growth phase/metabolic status (Fig 3B). In particular, as culture growth rates decrease approaching stationary phase, expression decreases from the strong pGPD (also known as pTDH3) promoter^89^, which drives the Aux/IAA fusion protein. In the case of our biosensor, this results in decreasing Venus-Aux/IAA17 measurements over time (Fig 3B). To mitigate these sources of noise, we engineered a bicistronic expression cassette using the Equine Rhinitis B Virus (ERBV) 2A self-cleaving peptide^86^ to co-express a Venus-Aux/IAA17 fusion and mScarlet-I from the same promoter. The Equine Rhinitis B Virus (ERBV) 2A peptide has the highest reported cleavage efficiency of 91% of 2A sequences tested in yeast^86^. We term this design “single-fusion” as we express untagged TIR1 or AFB2 with a Venus-Aux/IAA17-2A-mScarlet ratiometric Aux/IAA17 degradation reporter. Changes in the intracellular accumulation of Venus-Aux/IAA17 due to transcription/translation rate, cell division, and cell expansion can be accounted for using the mScarlet-I internal control. Therefore, the ratio of Venus-Aux/IAA17 to mScarlet should more accurately reflect the auxin-induced degradation of Venus-Aux/IAA17.

**Figure 3.**
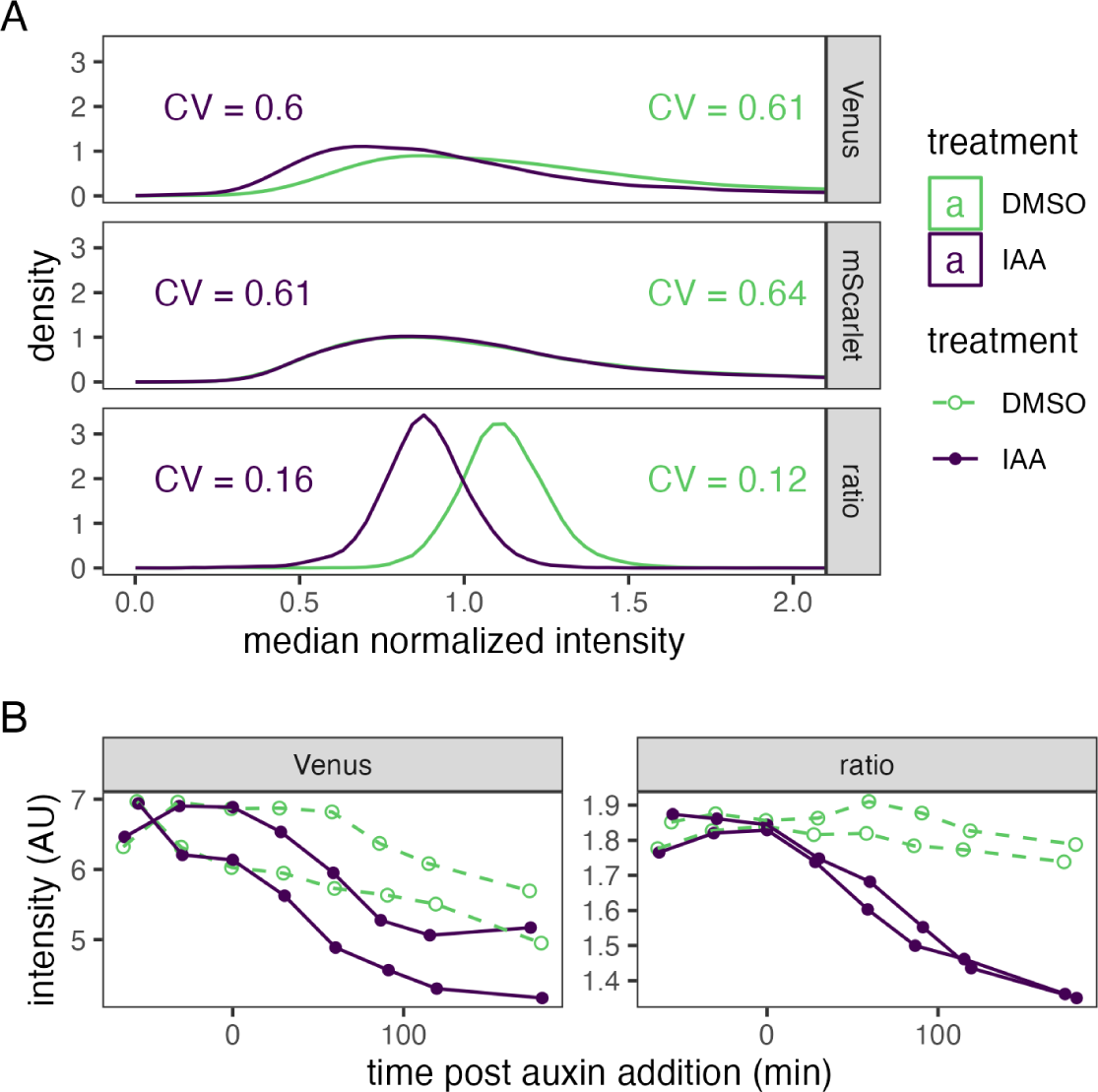
Ratiometric measurement of auxin-induced protein degradation reduces clonal and growth phase variation relative to single color measurement. (A) Two cultures of a yeast strain expressing an Venus:IAA17 fusion and free mScarlet from the same cistron, as well as TIR1 from a separate cistron were treated with 50 µM indole-3-acetic acid (IAA) or DMSO (vehicle control) for two hours prior to measurement of fluorescent intensity of single cells by flow cytometry. Values were normalized to the median of each culture and fluorophore or the ratio of raw Venus to mScarlet measurements. Lines represent kernel density estimates. Coefficients of variation (CV) for each population are shown. (B) Two cultures each of two independent clones of the yeast strain in A were treated as in A at time zero. Fluorescence intensity was measured by time-course flow cytometry. Population means are presented for each timepoint with each clonal culture shown as a separate line.

Additionally, it is possible that variation in accumulation of TIR1 or AFB2 contributes to noise in the measurement of auxin-induced degradation of Venus-Aux/IAA17. We aimed to test this hypothesis, and perhaps further increase the accuracy of our measurements of the SCF^TIR1/AFB2^–auxin-induced degradation of Venus-Aux/IAA17, by simultaneously measuring Venus-Aux/IAA17 and TIR1/AFB2-mScarlet-I fusions. To do this we inserted the *TIR1* or *AFB2* coding sequence between the 2A peptide and *mScarlet-I* coding sequences generating a bicistronic expression cassette of Venus-Aux/IAA17 and TIR1/AFB2-mScarlet-I fusion proteins. We term this design “dual-fusion” as we express TIR1/AFB2-mScarlet-I fusions with Venus-Aux/IAA17. Although we expect there to be similar variation in expression with the two fusions integrated into the genome at one or two loci^90^, we decided to engineer the fusions as a single cistron, again using the 2A peptide, to decrease variation if this biosensor is moved to a plasmid. The dual-fusion biosensor design also has the advantage of being a single ratiometric expression cassette offering fewer steps of genetic manipulation and cell transformation. Additionally, this dual-fusion biosensor allows direct measurement of both protein components of the TIR1/AFB2-auxin-Aux/IAA co-receptor complex, which should provide more accurate measurements of functional and expression variation between co-receptor complex sequence variants.

### Ratiometric biosensors reduce cell-to-cell, clonal and growth phase variation

To assess the ability of the single-fusion and dual-fusion ratiometric auxin biosensor designs to reduce variation in signal output, two cultures of independent transformants of the TIR1 versions of these biosensors in *S. cerevisiae* W303-1a were treated with auxin (50 µM IAA) or DMSO vehicle control and fluorescence of these cultures was measured over time by flow cytometry (Fig 3). Cell-to-cell variation can be represented by the coefficient of variation (CV) of the distribution of fluorescence for sample populations of each culture. For the single-fusion biosensor design, the CV of individual Venus-Aux/IAA17 and mScarlet-I measurements was approximately 4-fold greater than that of the ratio of Venus-Aux/IAA17 to mScarlet-I (Fig 3A). This reduction in cell-to-cell variation improves differentiation of the auxin treated population from vehicle control, in turn improving potential measurements of auxin or protein variant function using this ratiometric biosensor. The clonal and growth phase variation were also qualitatively reduced by using the ratiometric measurement compared to Venus-Aux/IAA17 alone (Fig 3B). This improvement will allow more accurate measurements across growth phases and of mutant yeast strains with altered metabolisms.

For the dual-fusion biosensor design, we obtained similar decreases in cell-to-cell variation (Fig 4). Ratiometric measurements of Venus-IAA17 to TIR1/AFB2-mScarlet-I have ∼3-fold lower CV compared to individual measurements. Interestingly, with the dual-fusion biosensor design we do not see as pronounced auxin induced degradation of Venus-IAA17 compared to the single-fusion biosensors with unlabeled TIR1/AFB2. We suspect this is due to the destabilizing effect of the mScarlet-I fusions, which due to the increased bulk may have increased rates of auto-ubiquitination of these SCF complexes^91–93^. While we recognize this is an imperfect comparison, the fluorescence of free mScarlet-I before treatment in the single-fusion biosensors is ∼3-fold higher than the fluorescence of TIR1/AFB2-mScarlet-I in the dual-fusion biosensors (Fig 5). In accordance with this hypothesis, auxin treatment may increase TIR1/AFB2-mScarlet-I accumulation (Fig 4), as perhaps Venus-IAA17 is recruited and ubiquitinated instead. This results in the ratiometric measurement for the dual-fusion biosensor having very similar behavior to the single-fusion biosensor (Fig 3B and Fig 6).

**Figure 4.**
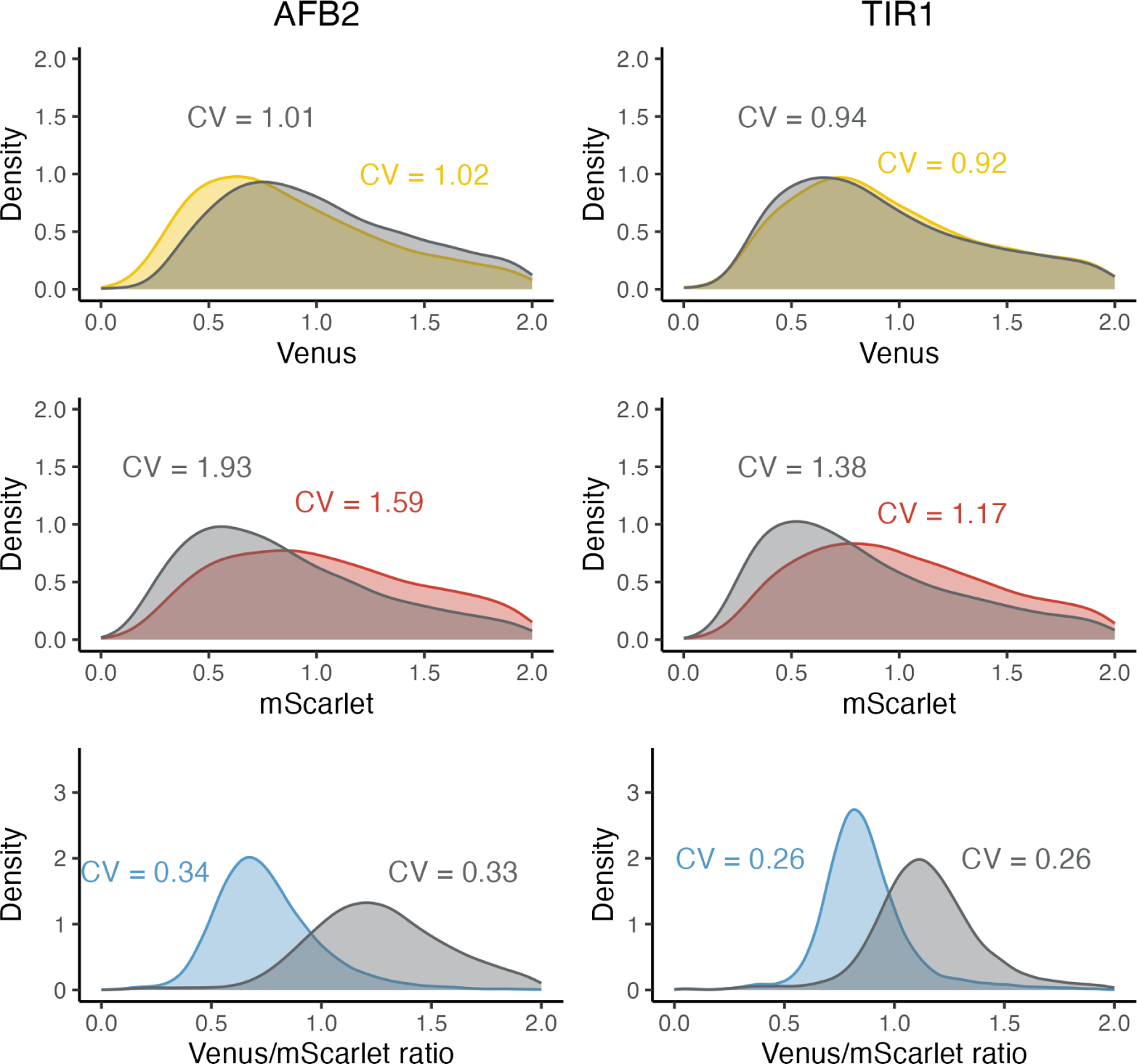
Ratiometric dual-fusion auxin biosensors also reduce cell-to-cell variation in yeast. Distributions of Venus (top), mScarlet (middle), and Venus/mScarlet (bottom) fluorescence in samples of 10^4^ cells from YPH499 yeast expressing AFB2 (left) or TIR1 (right) dual-fusion biosensors ∼4 hours after treatment with 50 µM auxin (colored) or vehicle control (gray). All data were normalized to the median of the combined treated and control distributions on each plot to center the distributions. Distributions are shown as kernel density estimates. The coefficients of variation for each population set are highlighted. The CVs for the Venus/mScarlet ratio is reduced about three-fold from the single Venus-Aux/IAA17 fluorescence measurement.

**Figure 5.**
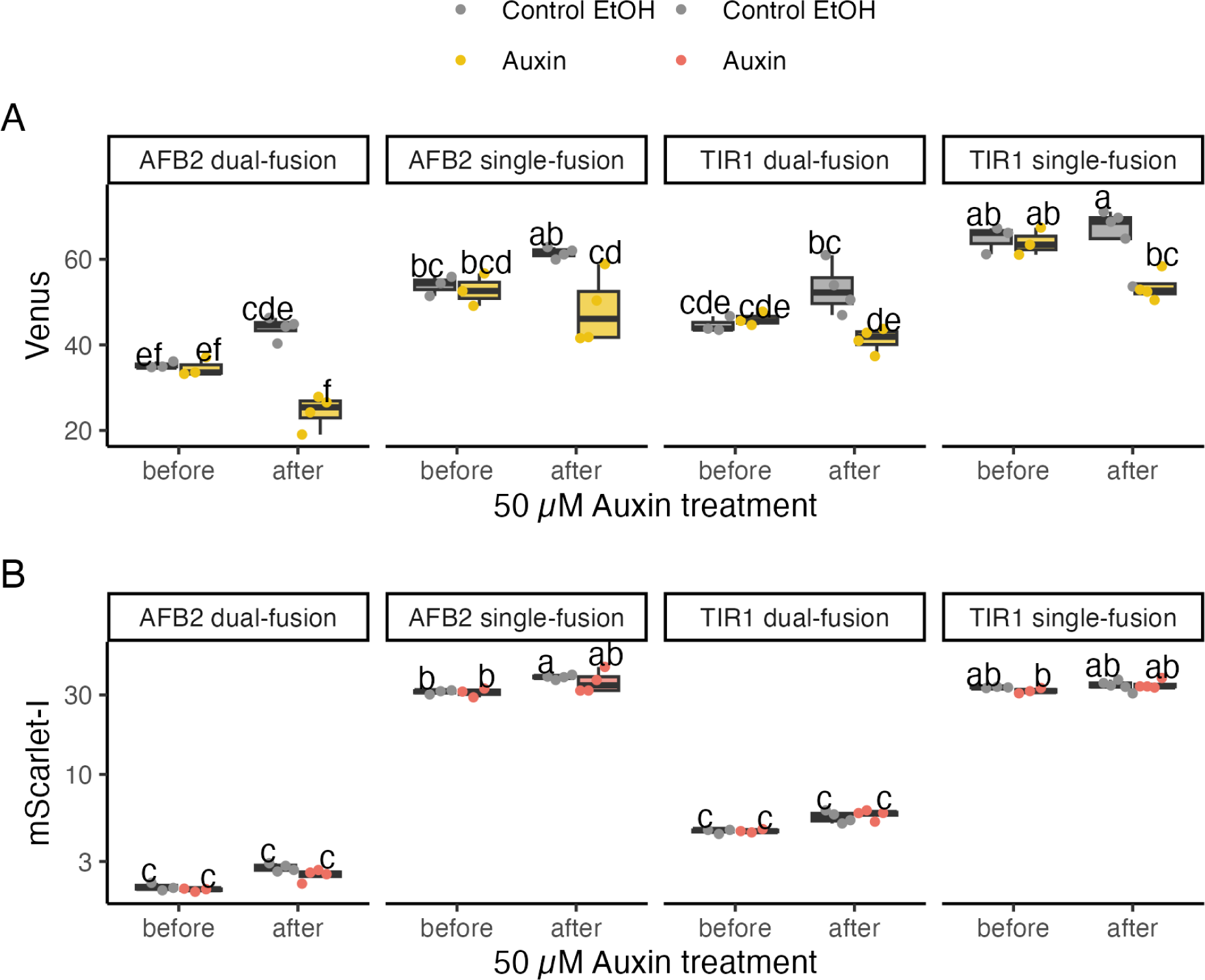
Dual-fusion designs have less Venus and mScarlet-I and fluorescence, but show similar changes in fluorescence in response to auxin treatment. Comparison of Venus and mScarlet-I fusion protein expression in auxin biosensor constructs before and after the treatment. **A)** Venus-Aux/IAA17 fluorescence and **B)** Free mScarlet-I (single-fusion) or TIR1/AFB2-mScarlet-I fluorescence (dual-fusion) of each biosensor design at steady-state before treatment and after treatment greater than 3 hours. Points represent mean fluorescence of samples of ∼10^4^ cells from cultures treated with 50 µM auxin (yellow and red) and the equivalent concentration of ethanol as a vehicle control (grey). Boxplots made from the 3-5 time point measurements are presented underneath the points. Letters above each group represent the group resulting from Tukey’s post-hoc of ANOVA.

**Figure 6.**
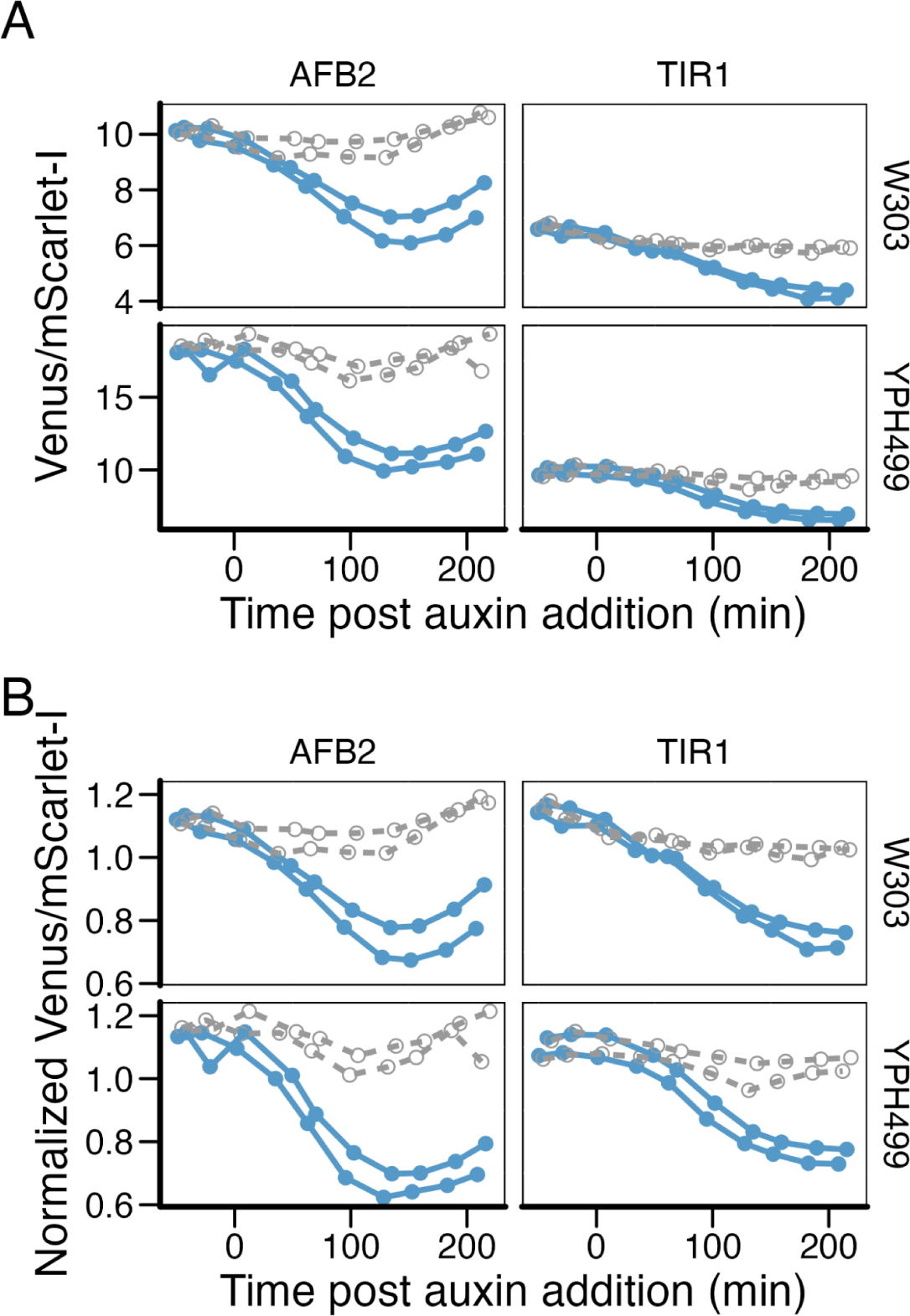
Dual-fusion biosensors show similar behaviors in two genetically distinct yeast strains in response to exogenous auxin, but have different levels of basal fluorescence. Duplicate cultures of independent transformants of W303 or YPH499 yeast expressing TIR1 or AFB2 dual-fusion biosensors early in the exponential growth phase were treated with 50 µM final concentration of auxin (IAA, solid blue), or an equivalent dilution of ethanol as a vehicle control (dashed gray). Fluorescence was measured over time by flow cytometry of samples of ∼10^4^ cells and mean fluorescence values are plotted. (B) Mean fluorescence values in A were normalized to the overall mean of values on each subplot to facilitate relative comparison of dynamics.

Comparing TIR1 to AFB2 in the both biosensor designs, AFB2 induces a more rapid decrease in the Venus/mScarlet-I ratio than TIR1 (Fig 4 and 6), as expected based on the more rapid degradation of IAA17 in the presence of AFB2^27^. AFB2 biosensors also result in lower absolute values of Venus-IAA17 fluorescence after auxin treatment than TIR1 for each biosensor design. Comparing the dual-fusion and single-fusion biosensor designs, both Venus-IAA17 and mScarlet-I (free for single-fusion, vs TIR1/AFB2-mScarlet-I fusion for dual-fusion) are lower for the dual-fusion biosensors (Fig 5. This is perhaps due to decreased stability of the longer dual-fusion biosensor transcript, as it contains the additional ∼2 kb TIR1 or AFB2 coding sequence or other transcriptional and translational burdens that are compounded in this single cistron construct. Lower Venus-IAA17 fluorescence in the dual-fusion constructs compared to single-fusion supports this hypothesis. However, the dual-fusion design does allow quantification of the relative levels of TIR1 and AFB2. TIR1-mScarlet-I accumulated to higher levels than AFB2-mScarlet-I, with a less than 2-fold difference in fluorescence that is consistent before and after treatment, while mScarlet fluorescence is very similar across all measurements of the single-fusion biosensor. In combination, this data suggests that AFB2 expression results in a higher rate of IAA17 ubiquitination and degradation than TIR1 in both the presence and absence of exogenous auxin.

To examine the portability of these biosensors between different yeast strains, and the potential for using these biosensors as to screen diverse yeast strains for auxin biosynthesis, we also tested the dual-fusion biosensors for consistent function in two distinct strains of *S. cerevisiae*, YPH499 (which is congenic to S288C)^94–96^ and W303 (which is an hybrid of S288C, ∑1278B, and perhaps other strains)^97,98^(Fig 6). These strains accumulate different levels of auxin in stationary phase cultures as measured by the colorimetric Salkowski reagent^50^. While the basal expression of the biosensors varied, with the Venus/mScarlet-I ratio being nearly 2-fold higher in YPH499 compared to W303, the qualitative behavior is similar in these strains, particularly if normalized to the average ratio of the vehicle control (Fig 6B). We expect the difference in absolute expression level is due to differences in cell size and shape, as well as expression, fluorophore maturation and basal degradation rates between these strains. Regardless of the mechanism, calibration of these biosensors is likely required when working with different strains. However, based on the similar dynamic behavior, we expect these biosensors to function similarly in comparative measurements such as dose-response or mutant analysis.

### Ratiometric auxin biosensors can accurately detect auxin from nanomolar to micromolar levels

We next examined the ability of these biosensors to detect auxin by measuring the biosensor response to a range of exogenous auxin additions in exponential phase cultures (Fig 7). We envision these biosensors may be useful for screening mutant yeast and perhaps other fungi which differentially accumulate auxin. We focus on the single plasmid/construct dual-fusion biosensors in this application to minimize genetic manipulation related to the auxin biosensor. Yeast cultures expressing the dual-fusion TIR1 or AFB2 biosensor in the exponential growth phase were treated with varying concentrations of auxin and the biosensor responses were measured over time by flow cytometry until steady-state fluorescence and/or stationary phase was reached. We define the biosensor response here as the ratio of TIR1/AFB2-mScarlet-I to Venus-IAA17, yielding a response that is directly proportional to auxin.

**Figure 7.**
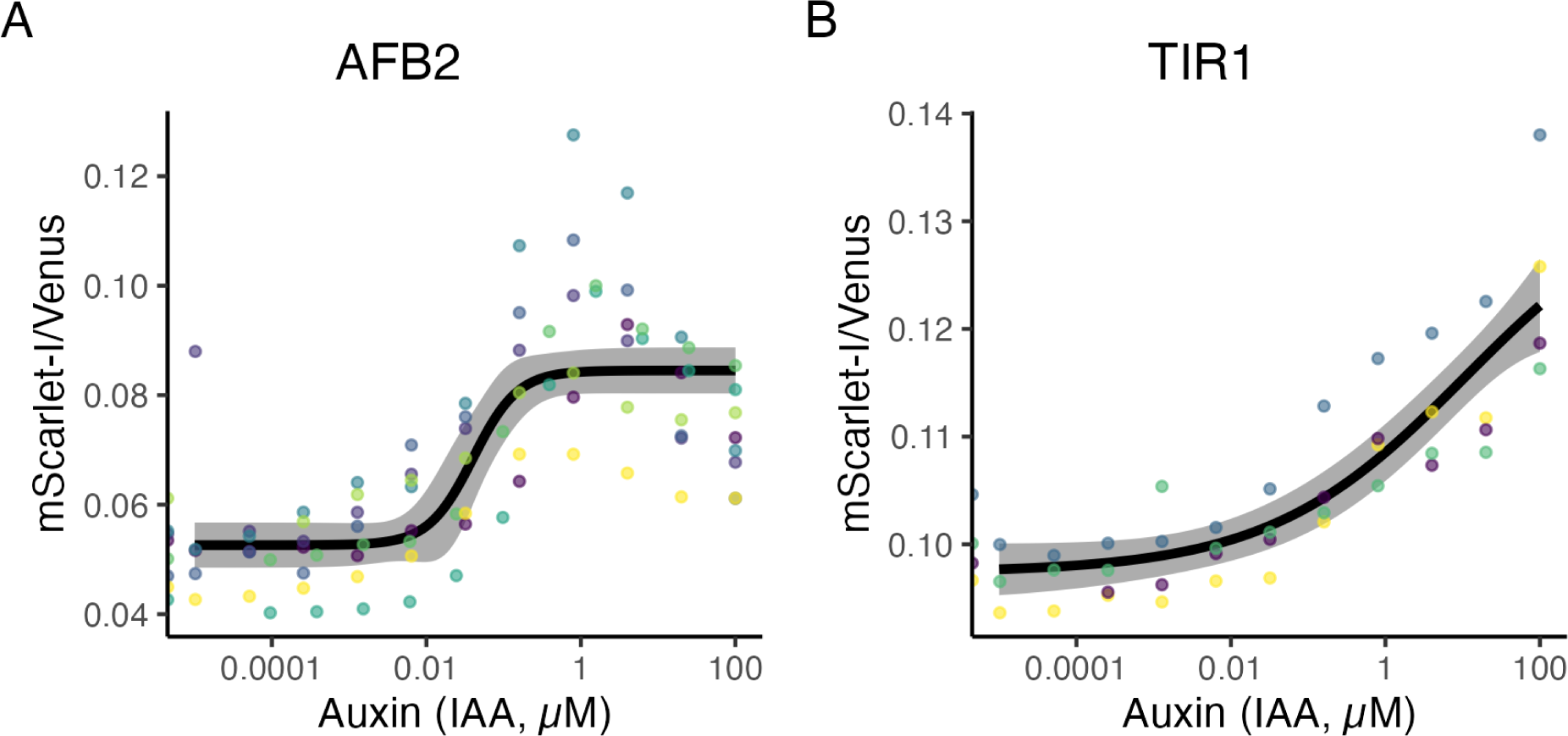
In combination the TIR1 and AFB2 dual-fusion biosensors respond to auxin concentrations spanning more than 5 orders of magnitude. Auxin dose-response curves were generated at exponential phase for the TIR1 and AFB2 dual-fusion biosensors. Biosensor-expressing yeast cultures were treated with different doses of auxin (indole-3-acetic acid, IAA) and the fluorescence ratio of TIR1/AFB2-mScarlet-I to Venus-Aux/IAA17 was measured at steady-state, ∼4 hours after treatment. **A)** Auxin dose-response curve for the AFB2 dual-fusion biosensor, based on 7 experimental replicates with different colonies on different days (points in different colors). A log-logistic model is shown as a black line, with 95% confidence interval in gray. The overall EC50 for AFB2 dual-fusion was 0.040 µM (SE = 0.016). **B)** The TIR1 dual-fusion dose-response curve, based on 4 experimental replicates, had an EC50 of 10.91 µM (SE = 20.40), however this value is poorly defined.

As expected the AFB2 biosensor is more sensitive, having a lower effective concentration for 50% response (EC50) than TIR1 (Fig 7) in correspondence with the higher Aux/IAA17 degradation rate^27^. The EC50 for the AFB2 biosensor is ∼40 nM, whereas for TIR1 it is ∼11 µM, a span of nearly 3 orders of magnitude—likely covering the physiological range of auxin concentrations in plant cells^99,100^. The EC50 for the TIR1 biosensor is less well defined as saturating levels of auxin defining the top of the dose response curve inhibit yeast growth^101^. The TIR1 biosensor has a much less steep response curve than AFB2, and is therefore sensitive to a wider range of concentrations, spanning from ∼100 nM to >100 µM, and perhaps up to the highest physiologically relevant auxin treatment levels, in the low mM range^81,102^. The steep slope of the AFB2 curve spans concentrations from ∼10 nM to 1 µM. We have found via LCMS measurements of yeast lysates that during exponential growth yeast cells typically contain around 47 nM auxin (SD = 15 nM), nearly equivalent to the EC50 of the AFB2 biosensor (Fig. 8). The lower levels of Venus-IAA17 observed in the presence of AFB2 than TIR1, as in Fig 5, may be due to the sensitivity of the AFB2-based biosensors to the level of endogenous auxin. This implies that the AFB2 biosensor should allow accurate measurement of perturbations in auxin biosynthesis during exponential growth of yeast. It may be possible to build additional biosensors spanning and expanding upon this range of auxin sensitivity for a variety of applications by testing other Aux/IAAs or through mutagenesis of the biosensors described here.

**Figure 8.**
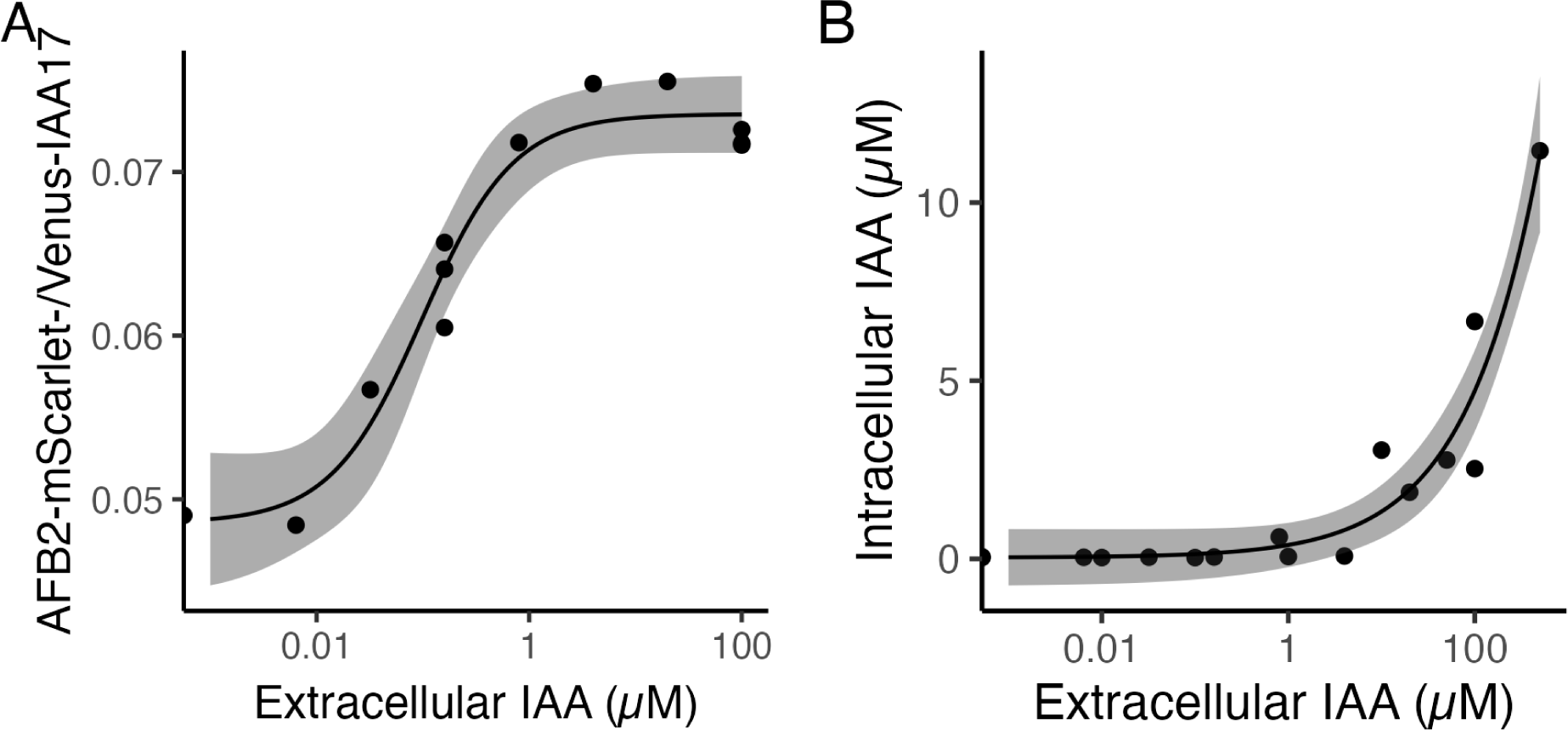
The AFB2 dual-fusion biosensor responds to extracellular auxin concentrations that did not result in significant changes in intracellular auxin by LCMS measurements. (A) AFB2 dual-fusion biosensor response to various concentrations of extracellular auxin after 3 hours (EC50 = 0.099 µM, SE = 0.031). (B) Following biosensor dose response experiments in A and another experiment, cells were harvested from cultures at various extracellular concentrations, and lysed. LCMS measurements of intracellular auxin were performed using deuterated IAA as an internal standard.

The AFB2 biosensor is able to measure very fine-grain changes at low levels, ∼10 nM exogenous auxin. Interestingly, using LCMS to measure intracellular auxin we do not observe significant deviation above the background level of ∼40 nM when less than 100 nM exogenous auxin is added to the culture medium (Fig. 8B). This could be due to the difference in spatiotemporal resolution of these two methods. Our biosensors are likely to primarily function in the nucleus of the yeast cells, whereas LCMS is limited to measuring whole cells as cell lysis is required. Additionally, the required sample preparation for LCMS could affect these measurements.

### Auxin accumulation in stationary phase is demonstrated by the AFB2 dual-fusion auxin biosensor

We and others^103–105^ have previously observed a steady decrease in fluorescence of cells expressing Aux/IAA-fluorescent protein fusions and TIR1/AFB auxin receptors as cultures approached stationary phase (Fig 3B). This could be due to auxin accumulation^36,50^ or changes in the GPD promoter activity used to drive expression of these proteins as cultures approach stationary phase. The bicistronic internal control, *mScarlet-I* or *TIR1/AFB2-mScarlet-I*, in our ratiometric auxin biosensors allows us to rule out the alternative hypothesis by controlling for changes in expression level (Fig. 2). To test whether our biosensor is able to detect this auxin accumulation as cultures enter stationary phase, we prepared cultures of our biosensor strains starting at typical cell density, as well as 10-and 100-fold higher cell densities so that these cultures would simultaneously be in exponential growth and stationary phase the following day. Indeed, our AFB2 dual-fusion ratiometric auxin biosensor predicts that auxin accumulates in stationary phase cultures (Fig. 9A), in agreement with previous literature^36,50^. The mean ratio of mScarlet-I to Venus-IAA17, which is proportional to the auxin concentration in our dose response studies, increases from exponential phase to early and then late stationary phase, and from aerobic to anaerobic conditions for cultures with the same initial conditions. To confirm that the biosensor is still responsive to auxin in early stationary phase cultures, we performed an exogenous auxin dose response assay with these cultures (Fig. 9B). The biosensor responded similarly under these conditions when accounting for the auxin accumulated from biosynthesis increasing the low end of the curve.

**Figure 9.**
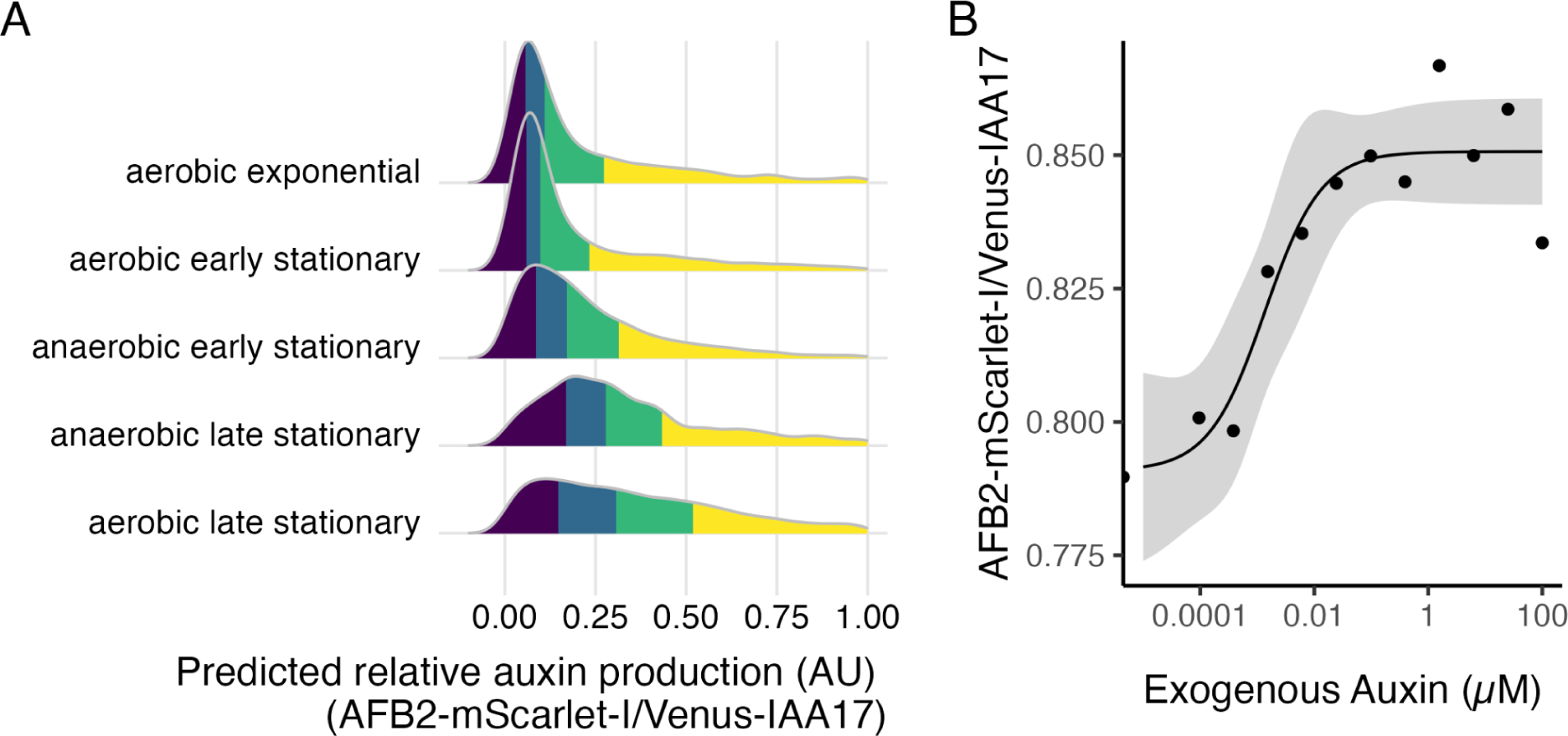
Dual-fusion AFB2 biosensor predicts that auxin accumulates in stationary phase cultures. **A)** Yeast expressing an auxin biosensor were inoculated at varying rates to reach different phases of growth after 16 hours of incubation in static fermentative (anaerobic) conditions or shaken (aerobic) conditions. Distributions represent flow cytometric measurements of the ratio of fluorescent intensity areas of AFB2-mScarlet to Venus-Aux/IAA17, which is proportional to the predicted/perceived auxin. Each quartile of the distributions is represented by a different color, with the line between blue and green representing the median of the distribution. **B)** Dose-response curve for auxin at early stationary phase. The ratiometric auxin biosensor responds to exogenous auxin in the stationary phase. The EC50 is 0.00137 µM exogenous IAA (SE =0.00094).

### Ratiometric biosensors facilitate FACS enrichment of cells based on auxin accumulation or perception

Fluorescent biosensors are often paired with fluorescence-accelerated cell sorting (FACS) to facilitate screening of variant libraries for directed evolution, strain engineering, and genetics. To demonstrate the potential use of these biosensors in these applications, we have established a system for the directed evolution of TIR1/AFB–Aux/IAA auxin co-receptor complexes. Here, we demonstrate a ratiometric FACS gating strategy and the ability of the single-fusion biosensor to report on functional variation in an *AFB2* random mutagenesis library. The reduced cell-to-cell variation of these ratiometric biosensors should enhance our ability to sort variant libraries by auxin accumulation or auxin perception using FACS.

For example, we could enrich for variants of *TIR1* or *AFB2* genes that have improved ability to induce specific degradation of Venus-Aux/IAA substrate proteins by sorting yeast cells based on distinct population shifts in a projection of Venus-Aux/IAA17 vs mScarlet-I measurements relative to the parental variant, for the single-fusion biosensor (Fig. 10). For example, after auxin treatment, sorting all cells which fall below and to the right of the wild-type population, which falls right along the diagonal in Fig. 10 would enrich cells expressing *TIR1/AFB* variants with greater ability to induce Venus-IAA17 degradation. Similarly, loss-of-function variants could be enriched by sorting those cells above and to the left of the wild-type in Fig. 10. A gate shifted further below and to the right of the auxin-treated population could be used to enrich for increased sensitivity in auxin perception or increased auxin production from yeast variant libraries.

**Figure 10.**
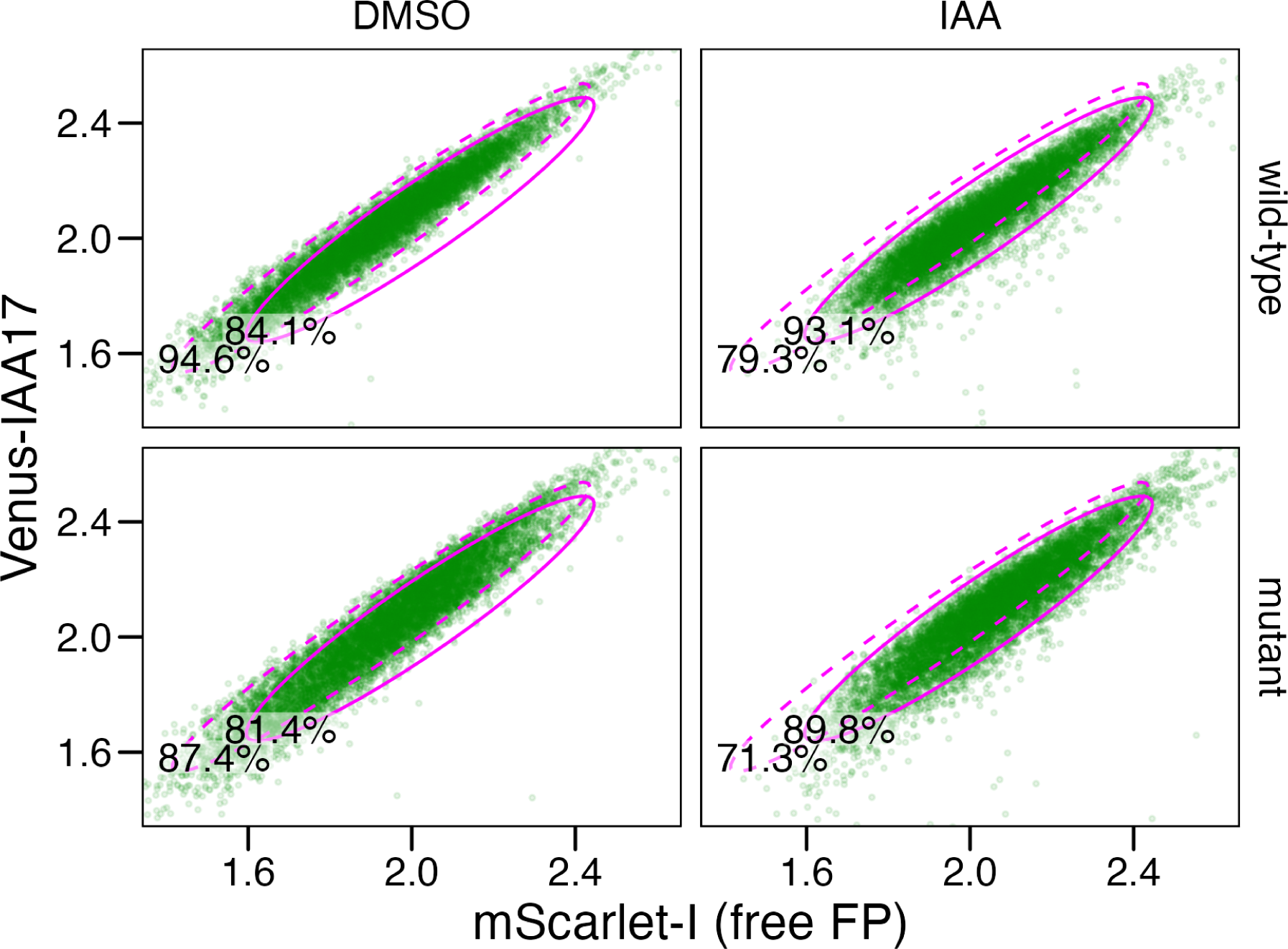
Ratiometric auxin biosensors facilitate fluorescence accelerated cell sorting of yeast for studying and engineering auxin co-receptor proteins or auxin biosynthesis pathways. Yeast cultures expressing the single-fusion auxin biosensor, with AFB2 expressed from a cytosolic plasmid that is replicated by wild-type high-fidelity polymerase (resulting in wild-type AFB2), or mutant error-prone polymerase (resulting in AFB2 mutants), were treated with DMSO (vehicle control) or auxin (50 µM indole-3-acetic acid) for 6 hours. Scatter plots are shown with points representing individual events. Two elliptical gates were drawn (pink), one to contain ∼95% of the DMSO-treated wild-type population (dotted line), and another to contain ∼95% of the IAA-treated wild-type population (solid line). The percent of events in each gate is shown in the lower left corner of the gate.

To demonstrate this use-case of these biosensors we prepared a library of *AFB2* variants using OrthoRep *in vivo* mutagenesis^106,107^. In brief, we integrated the single-fusion auxin biosensor (pGPD::Venus-Aux/IAA17:2A:mScarlet-I) into the *trp1* locus of F102-2 yeast. We then co-transformed this strain with a AFB2 expression cassette with arms of homology to the linear cytosolic p1 plasmid along with a CEN/ARS plasmid expressing expressing either an error prone or high fidelity polymerase that specifically replicates the P1 plasmid. We then selected co-transformants on solid media and inoculated cultures with 3 colonies from these plates and incubated the culture for 24 hours. Throughout the growth of these cultures mutations will accumulate in the p1 plasmid, which contains *AFB2*, at a much higher rate in the presence of the mutant error prone polymerase (10^-5^ substitutions per base pair or s.p.b.) than the wild-type high fidelity (10^-9^ s.p.b.)^107^. Based on the substitution rate, culture time, and length of *AFB2*, we predict that between 1.6% and 22% of the error prone polymerase culture contained a nonsynonymous mutation in *AFB2*. In the absence of IAA ∼7% more of a sample of the mutant population fell outside of an elliptical gate that contained ∼95% of a sample of the wild-type population. In the presence of IAA an additional ∼3% of the mutant sample fell outside of the gate which contained ∼93% of the wild-type sample (Fig 10).

We confirmed that the SCF^AFB2^ auxin-activated ubiquitin ligase complex is functional when AFB2 is expressed from a cytosolic plasmid capable of error-prone replication, with our single-fusion biosensor, which showed an auxin-specific decrease in the Venus-IAA17/mScarlet-I ratio over time (Fig 11). Additionally, the error prone polymerase expressing culture showed a broadening of the ratiometric response compared to the high fidelity polymerase (Fig 11). Interestingly, the increased breadth of this functional distribution for the *AFB2* library was primarily towards increased Venus-IAA17 degradation, resulting in a lower Venus-IAA17/mScarlet ratio. We speculate this is due to adaptation of the coding sequence for yeast, reducing stalled ribosomes and increasing translational efficiency, as there is constant growth selection for efficiently translating sequence variants^108^. Perhaps in support of this hypothesis, after 1 hour of auxin treatment the high-fidelity distribution has shifted further left than the error-prone distribution, but after 6 hours the error-prone distribution’s heavier left tail is more apparent. This suggests that the error-prone-polymerase-expressing culture is not necessarily enriched for improved dynamic ability to induce ubiquitination and degradation of Venus-IAA17, but that increased accumulation of these AFB2 variants affects the steady-state Venus-IAA17 levels. However, expressing *TIR1* from the pGPD promoter at one or two loci in the yeast genome does not significantly affect Aux/IAA degradation dynamics in yeast ^27^. Still, this may not be the case with *AFB2* or when it is expressed from the cytosolic p1 plasmid and p10B2 promoter. It is perhaps the case that an early mutation that occurred in the error-prone polymerase expressing culture had this slightly increased Venus-Aux/IAA17 degradation function and is simply overrepresented. To control for AFB2 expression level variants could be subcloned into the dual-fusion biosensor and screened in this system where AFB2-mScarlet-I can be measured directly. We will further characterize this library and others in future work.

**Figure 11.**
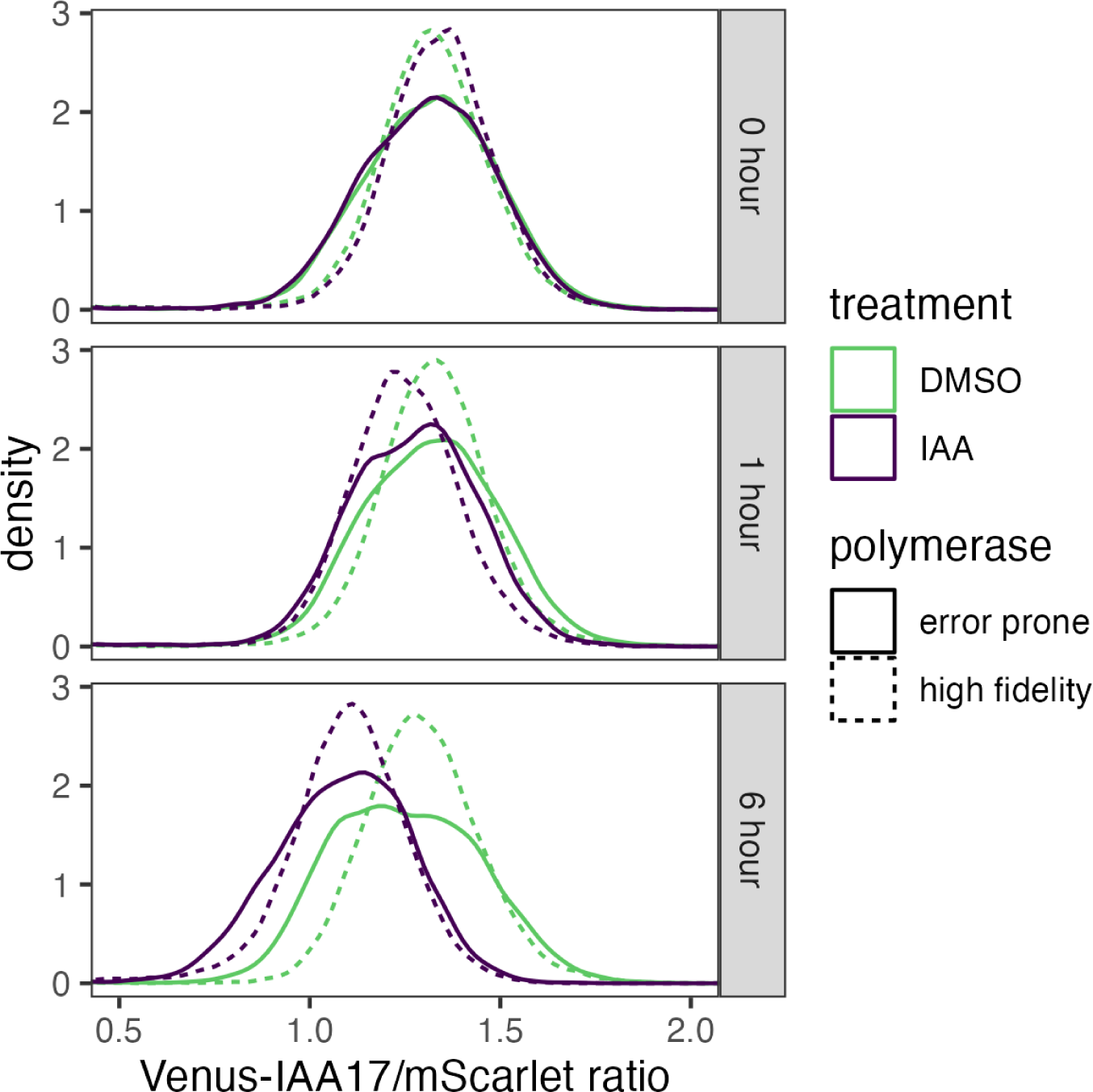
The single-fusion biosensor measures divergent function in an AFB2 variant library. Yeast expressing the single-fusion ratiometric sensor with the AFB2 auxin receptor replicated either by a high-fidelity polymerase (dotted line) or an error-prone polymerase (solid line) were cultured for 24 hours. Samples of these cultures were treated with 50 µM auxin (indole-3-acetic acid, IAA, green) or vehicle control (DMSO, purple). The ratio of Venus-IAA17 to mScarlet-I fluorescence intensity for a sample of 10^4^ cells was measured immediately prior to treatment and 1 and 6 hours post treatment (as indicated by vertical labels on right). The sample distributions are presented as kernel density estimates.

## Conclusion and Future Perspectives

We have developed a series of ratiometric auxin biosensors in *S. cerevisiae*. These biosensors are based on the plant nuclear auxin signaling machinery which varies widely within and between species. These ratiometric biosensors improve upon previous technologies by reducing cell-to-cell and clonal variability, making these biosensors more robust to different growth conditions and genetic backgrounds. Additionally, our dual-fusion biosensor design reports the accumulation of TIR1/AFB auxin receptors, providing additional mechanistic information. Through calibration of the response of these biosensors to exogenous auxin and LCMS measurements, as well as biosensor measurements at stationary phase when yeast cultures accumulate auxin, we have demonstrated that the dual-fusion AFB2 biosensor is capable of reporting auxin biosynthesis and accumulation in yeast. While candidate biosynthetic pathways for auxin in yeast and other fungi are known, these pathways are incomplete and their regulation has only been preliminarily studied^4,35–37,109–115^. The biosensors we have developed here provide new tools to understand how and why fungi synthesize and respond to auxin. These biosensors will allow exploration of the signaling and physiology of auxin in plant–fungi interactions as well. Single fluorescent protein reporters based on the same signaling mechanism as those presented here have been used to study auxin’s involvement in *Magnaporthe oryzae* infection of rice^84^. Auxin-induced protein degradation has been used in many eukaryotic species, and will likely function across fungi, with minimal alterations^72,75,116^. It may be possible to make single plasmid constructs that could be used to simultaneously examine a wide variety of yeast species^117^. Further optimization of auxin co-receptor proteins, 2A self-cleaving peptide, fluorescent proteins, and codon usage may be necessary to optimize measurement in other species, and may also improve the function of these sensors in *S. cerevisiae* and other fungi.

Here, we envision and provide preliminary proof-of-concept for two engineering use-cases for this biosensor: studying auxin biosynthesis in yeast and mutational scanning and directed evolution of auxin co-receptor pairs. Through biosensor-guided engineering, auxin biosynthesis in yeasts could be increased or decreased to examine the role of auxin in different microbiomes. However, our understanding of the metabolic pathways fungi use to produce auxin is still incomplete. Previous attempts to knock down one biosynthetic route to auxin in yeast actually resulted in an increase in auxin accumulation^35^ revealing complex regulation within the several biosynthesis pathways. Because the mechanism of this biosensor requires a functional ubiquitin-proteasome system, genetic screens for auxin biosynthesis will likely need to be targeted using CRISPR/Cas mutagenesis. However, it may be possible to devise a chemical screening strategy using auxinole, a TIR1/AFB antagonist^118^, and exogenous auxin to verify biosensor function for more wide-scale genetic screens. Performing mutational scanning of known auxin metabolic enzymes and transporters native to yeast, or heterologously expressed is certainly feasible with these biosensors. In these cases, a similar dual-fusion construct, with a fluorescent protein fused to the gene of interest may be useful. However it may also lead to unanticipated results as with the decrease in overall fluorescence of our dual-fusion constructs. While these constructs are capable of reporting auxin concentration, the large increase in TIR1/AFB2-mScarlet fluorescence compared to the slight decrease in VENUS-IAA17, demonstrates greatly reduced expression or function of TIR1/AFB2-mScarlet compared to free TIR1/AFB2. Similar mCitrine fusions however complement mutants in *Arabidopsis thaliana* when expressed from native promoters^119^. Perhaps this observation for the dual-fusion biosensors is a yeast-specific phenomenon. The single-fusion biosensor, which is more closely related to previous tools for assessing the function of nuclear auxin signaling pathway components in yeast^27,30,85^, may be more useful in assessing auxin co-receptor function, both due to its more canonical behavior and the ability to separately modify *TIR1/AFB* and *Aux/IAA.* Follow up experiments with similar modifications in dual-fusion biosensors can then be used to parse the effects of TIR1/AFB accumulation versus Aux/IAA degradation.

We have demonstrated how the single-fusion biosensor can be used to determine the distribution of function for a library of *TIR1/AFB* variants, and plan to perform comprehensive mutational scanning and directed evolution of these proteins in the future, to further refine the sequence function map of these multifaceted signaling proteins^120–125^ and to engineer novel functions^126^. For mutational studies in Aux/IAAs the single-fusion ratiometric sensor, also provides a simple quantitative reporter of Aux/IAA degradation, which would prove useful in studying the numerous functional elements of these transcriptional regulators^21,127–132^. While the ratiometric Venus-IAA17-2A-mScarlet-I construct would not be amenable to OrthoRep continuous mutagenesis, due to the included fluorescent protein reporters, we expect alternative means of creating specific *IAA17* variant libraries cloned between *Venus* and the 2A sequence would yield similar results. As we have observed here pairing of these single- and dual-fusion biosensors to measure auxin co-receptor ubiquitin ligase function from multiple perspectives can provide mechanistic insight into their function.

From a protein engineering perspective, the three-body problem presented by the formation of the TIR1/AFB-auxin-Aux/IAA complex^19,133^ is a fascinating system to study. Similar chemically activated ubiquitin ligase complexes also allow plants to perceive many other internal and environmental chemical signals^134^ including jasmonates^135,136^, gibberellins^137–140^, strigolactones^141,142^, and karrikins^143,144^, which spatially and temporally control gene expression and coordinate plant growth, development, and behavior, as well as shape microbial interactions^145^. It is possible that similar ratiometric biosensors could also be established for these signaling pathways, and some have already been established in plants^78,82,146^. We hope the biosensors we present here will shed additional light on how auxin is produced and perceived by fungi, as well as allow us to re-engineer this interkingdom signaling pathway for applications in agriculture, medicine, and biotechnology.

## Methods

### Plasmid construction

Primers for cloning DNA fragments for the ratiometric biosensor constructs were designed using the online DIVA J5 DNA assembly design tools (public-diva.jbei.org) ^147^. The complete list of primers used in this study can be found in Supplementary Tables S1. The DNA fragments and constructs containing *TIR1* or *AFB2*, Aux/IAA fused to the Venus and mScarlet fluorescent proteins were amplified and inserted into pGp4G2 and/or pGp8G2 downstream of a GPD promoter. The PCRs were performed with Q5® High-Fidelity 2X Master Mix (New England Biolabs) with the designed synthetic primers (SigmaAldrich). The PCR products of each synthetic part were purified, assembled, and inserted into the vectors via Gibson assembly. The mixture of DNA fragments was purified by the DNA cleaning kit before transforming into the competent *E. coli* cells via chemical transformation and plated onto the selective LB agar plates containing 100 ug/ml Ampicillin (Fisher Scientific). The list of *E.coli* cells can be found in Supplementary Tables S1. Negative control and positive control of pUC19 were included in the transformation process. The transformants are expected to be grown on the ampicillin selective LB agar plates overnight at 37°C. The grown colonies were selected and subjected to colony PCR prior to Sanger sequencing.

### Yeast transformation

The successfully cloned plasmids were digested with *PmeI* restriction enzyme (New England Biolabs) to linearize prior cell transformations into either W303 or YPH499 (MATa) yeast strains. All yeast strains are routinely stuck out on a sterile YPAD plate consisting of 20 g/L dextrose (Fisher Scientific), 20 g/L peptone (Fisher Scientific), 20 g/L yeast extract (Fisher Scientific), 20 g/L agar (Fisher Scientific), and 40 mg/L Adenine Hemisulfate (Fisher Scientific). The procedure for making competent yeast cells and yeast transformation were performed according to Gietz and Schiestl^148^. Briefly, a single colony of each yeast strain was grown in 2X YPAD media in a 250 mL Erlenmeyer flask at 30 °C, 250 rpm for 16 hours. The cultures were then diluted into fresh and pre-warm 2X YPAD medium, and incubated at 30 °C, 250 rpm for 4 hours prior to the cell harvesting process at the cell specific density established in the protocol. The harvested yeast pellets were also resuspended with the sterile frozen competent cells (FCC) solution (5% v/v glycerol (Fisher Scientific) and 10% v/v DMSO (Fisher Scientific) for stocks storing at −70 °C freezer. The complete list of *S. cerevisiae* strains in this study can be found in Supplementary Tables S1. The yeast transformants with an auxotrophic marker were expected to grow at 30 °C on the selective synthetic media plates. Yeast transformants were confirmed for correct integration by isolation on appropriate selective plates at 30 °C additional to yeast colony PCR. Yeast colony PCR was performed using Zymolyase Yeast lytic enzyme (Zymo Research) to lyse the cells. Briefly 3 units of zymolyase was mixed with a barely turbid solution of cells in a 15 µL volume. In a thermocycler the reaction was incubated for 30 min at 30°C followed by 10 min at 95°C. 85 µL of water was added to each lysate and 2 µL of this was used as template for a 20 µL Taq PCR containing primers to amplify from the terminator of the transgene expression cassette across the site of homologous recombination and ∼200 bases into genomic DNA. The presence of appropriate fluorescent protein expression in the colonies was confirmed using iBright Imaging Systems (Thermo Fisher Scientific) and flow cytometer. Successful yeast transformants were stored in 15% glycerol at −70°C.

### Flow cytometry measurements and data analysis

#### Auxin-induce degradation time-course assays

Each yeast strain carrying the ratiometric biosensor was struck to isolation on fresh YPAD plates. After 2–3 days incubation at 30°C, ∼¼ of a healthy uniform colony was suspended in complete synthetic medium (SCM) (Takara Bio USA). The cell concentration of each inoculum was measured by flow cytometry and then diluted to 1 cell/ul in an erlenmeyer flask. The cultures were grown overnight at 30 °C, 300 rpm. On the following day, at the exponential growth phase, the cell cultures were aliquoted into a 96-deep well plate. An Indole-3-acetic acid (IAA) working solution from 50 mM IAA in 48% ethanol stock solution (Fisher Scientific) was freshly prepared and added to the cell cultures to obtain a final concentration of 50 µM IAA. Another set of cultures were treated with the equivalent solvent control. All cultures were cultivated at 30°C, 300 rpm and a sample was measured for the change in fluorescence via cytometry every 30 minutes. All recorded events were annotated, and analyzed using the flowTime R package ^85,149^. Data analysis is documented in Supplementary File Data Analysis S2. Intra- and inter-day replications were performed using different yeast colonies.

#### Dose-response assays

Cultures were prepared following the protocol above. A stock solution 50 mM IAA in 48% ethanol was prepared and freshly diluted serially to obtain the final concentrations in the cultures ranging from 100 µM to 0 µM (typically 100, 20, 4, 0.8, 0.16, 0.032, 0.0064, 0.00128, 0.000256, 0.0000512, and 0.00001020 µM). At the exponential growth phase, different concentrations of IAA working solutions, including the solvent control, were added to each culture. For the stationary phase dose-response, the yeast was cultured in SCM in an erlenmeyer flask for 48 hours at 30 °C, 300 rpm. 1200 ul of the culture was aliquoted into each well of a 96-deep well plate, and incubated at 30 °C, 300 rpm. IAA was added to the aliquoted cultures as above. The cultures were diluted 50-fold in SCM and gently mixed with a multichannel pipette immediately prior to cytometer measurements to ensure isolated events. The growth of yeast cells and fluorescence signals in response to different doses of IAA were measured over time and plotted to determine when a new steady-state fluorescence was reached. All recorded events were annotated, and the data were analyzed using the flowTime R package ^85,149^ and the drc R package was used to fit 4-parameter log-logistic dose-response curves including the median effective concentration (EC_50_).

#### Auxin biosynthesis assays

Cultures were prepared following the protocol above, except that early and late stationary phase cultures were inoculated at 10 cells/µL and 100 cells/µL respectively. Fermentative/anaerobic cultures were not shaken. After 24 hours of incubation the cultures were diluted 50-fold in SCM and gently mixed with a multichannel pipette immediately prior to cytometer measurements to ensure isolated events.

### Quantification of intracellular IAA from yeast by LCMS

Cultures were prepared as above and during exponential growth phase 7 mL was collected, centrifuged, washed with 1000 ul sterile water, and resuspended in 200 µL of 50% acetonitrile. The suspension was mixed vigorously and loaded into a 2 mL lysing matrix Y tube (MP Biomedicals). After loading the sample, the tube was placed in a high-speed homogenizer (MP Biomedicals Fastprep-24 Sample Preparation System) to disrupt the cells at 6.5 m/s, 10 seconds, for 10 cycles. The lysed samples were incubated on ice for 30 s prior to centrifugation at 16,000 g for 2 minutes. Complete lysis was confirmed via light microscopy, by lack of intact cells in several fields of view. The lysate was then collected and stored at −20 °C for further analysis. Immediately prior to analysis the samples were centrifuged at 16,000 g for 5 min, diluted 1:100 with internal standard in 50% acetonitrile and transferred to a LCMS vial.

LCMS analysis was performed on a Shimadzu Nextera X2 UPLC interfaced with a Shimadzu 8060 triple quadrupole mass spectrometer. A 5-minute binary gradient starting at 80% solvent A, water with 0.1% formic acid, and 20 % solvent B, methanol with 0.1% formic acid was used for the analysis with a flow rate of 0.4 ml/min. The gradient conditions were isocratic for 1 minute, a linear gradient to 90% B at 3.5 min and returning to initial conditions at 4.1 minutes. A 5 ul aliquot was injected onto a Shimadzu Nexcol C18 1.8 um, 50 x 2.1 mm column maintained at 35C. Deuterated indole-3-acetic acid (d7-IAA, Cambridge Isotope Laboratories) was used as an internal standard at a concentration of 75 nM in all samples and standards. The mass spectrometer was operated in positive mode and the MRM transition for IAA quantification was 176->130 and the MRM transition for d7-IAA was 183->136. The limit of detection was determined to be 5 nM and all standards and samples were analyzed in triplicate. The peak area of IAA was normalized to the internal standard and quantification was based upon a standard curve prepared with a purchased IAA standard (Sigma-Aldrich, I15148).

### OrthoRep continuous in vivo mutagenesis

To build a yeast strain capable of continuous mutagenesis and FACS by variant function we initially integrated the *single-fusion* biosensor reporter construct *TRP1:pGPD:Venus::Aux/IAA17::2A::mScarlet-I* into the *trp1* locus of F102-2 yeast, via homologous recombination and selection for *TRP1* prototrophy. We then co-transformed this strain with a *TIR1* or *AFB2* expression cassette with arms of homology to the linear cytosolic p1 plasmid based on FDP-p10B2-URA3 along with a HIS3+ CEN/ARS plasmid expressing expressing either an error prone or high fidelity polymerase that specifically replicates the P1 plasmid, using plasmids and protocols as per the OrthoRep publications^107,150^. We then selected co-transformants on solid SCM lacking tryptophan, uracil, and histidine. Liquid cultures of 50 mL of the same medium were inoculated with 3 colonies from these plates and incubated for 24 hours at 30°C and 250 RPM. Samples of these cultures were treated with 50 µM auxin or vehicle control (0.1% DMSO). The ratio of Venus to mScarlet-I fluorescence intensity for each cell was measured immediately prior to treatment and periodically for 6 hours post treatment via flow cytometry.

## Code availability

Code for analyzing all data and creating all plots is available on github (https://github.com/PlantSynBioLab/auxin-biosensor-data) and in Supplementary File Data Analysis S2. All source data is provided in the paper, supplemental code via github, and will be deposited on flowRepository prior to publication.

## Supporting information

Supplementary File Data Analysis S2

## Acknowledgements

Research in the Wright Plant Synthetic Biology Laboratory is supported by the United States Department of Agriculture National Institute of Food and Agriculture Agriculture and Food Research Initiative Plant Breeding for Agricultural Production (Grant No. 2022-67013-36293) and Hatch Project [VA-1021738]; The Virginia Space Grant Consortium Innovative Project Grant; Virginia Tech Institute for Critical Technologies and Sciences Junior Faculty Award; and Virginia Tech College of Agriculture and Life Sciences Strategic Plan Advancement 2021 Integrated Internal Competitive Grants through the Center for Advanced Innovation in Agriculture. This work was conceived and initially supported by the National Science Foundation Postdoctoral Research Fellowship in Biology (NSF-1402222) to R.C.W. Additional support for P.C. was provided by the Ministry of Science and Technology, Royal Thai Government. BioRender was used to create the graphical abstract and figures 1 and 2. We would like to thank Chang C. Liu and Jennifer Nemhauser for providing plasmids and yeast strains. We also would like to thank Arjun Khakhar and Bastiaan Bargmann for the helpful discussions and reading of the manuscript.

## Author contributions

P.C., M.R., and R.C.W conceptualized and designed the study. P.C. and R.C.W wrote, edited and designed all figures for the manuscript. All authors edited and approved of the final manuscript. Contributions to the experimental work include plasmid design, cloning, and strain constructions (P.C., M.R., S.M., C.G., and R.C.W.); dose-response and auxin induce degradation assay (P.C.); biosensor calibration and applications (P.C., M.R., and R.C.W); LC-MS analysis (R.H. and S.H); general data and statistical analysis (P.C., M.R., and R.C.W).

## Notes

All authors declare no competing financial interest

## References

1. Kunkel, B. N.; Harper, C. P. The Roles of Auxin during Interactions between Bacterial Plant Pathogens and Their Hosts. J. Exp. Bot. 2018, 69 (2), 245–254. https://doi.org/10.1093/jxb/erx447.

2. Ludwig-Müller, J. Bacteria and Fungi Controlling Plant Growth by Manipulating Auxin: Balance between Development and Defense. J. Plant Physiol. 2015, 172, 4–12. https://doi.org/10.1016/j.jplph.2014.01.002.

3. Qian, J.-M.; Bai, Y. Stuck on You: Bacterial-Auxin-Mediated Bacterial Colonization of Plant Roots. Cell Host Microbe 2021, 29 (10), 1471–1473. https://doi.org/10.1016/j.chom.2021.09.014.

4. Fu, S.-F.; Wei, J.-Y.; Chen, H.-W.; Liu, Y.-Y.; Lu, H.-Y.; Chou, J.-Y. Indole-3-Acetic Acid: A Widespread Physiological Code in Interactions of Fungi with Other Organisms. Plant Signal. Behav. 2015, 10 (8), e1048052. https://doi.org/10.1080/15592324.2015.1048052.

5. Chanclud, E.; Morel, J.-B. Plant Hormones: A Fungal Point of View. Mol. Plant Pathol. 2016, 17 (8), 1289–1297. https://doi.org/10.1111/mpp.12393.

6. Chanclud, E.; Lacombe, B. Plant Hormones: Key Players in Gut Microbiota and Human Diseases? Trends Plant Sci. 2017, 22 (9), 754–758. https://doi.org/10.1016/j.tplants.2017.07.003.

7. Omelyanchuk, N. A.; Wiebe, D. S.; Novikova, D. D.; Levitsky, V. G.; Klimova, N.; Gorelova, V.; Weinholdt, C.; Vasiliev, G. V.; Zemlyanskaya, E. V.; Kolchanov, N. A.; Kochetov, A. V.; Grosse, I.; Mironova, V. V. Auxin Regulates Functional Gene Groups in a Fold-Change-Specific Manner in Arabidopsis Thaliana Roots. Sci. Rep. 2017, 7, 2489. https://doi.org/10.1038/s41598-017-02476-8.

8. Kato, H.; Nishihama, R.; Weijers, D.; Kohchi, T. Evolution of Nuclear Auxin Signaling: Lessons from Genetic Studies with Basal Land Plants. J. Exp. Bot. 2018, 69 (2), 291–301. https://doi.org/10.1093/jxb/erx267.

9. Todd, O. E.; Figueiredo, M. R. A.; Morran, S.; Soni, N.; Preston, C.; Kubeš, M. F.; Napier, R.; Gaines, T. A. Synthetic Auxin Herbicides: Finding the Lock and Key to Weed Resistance. Plant Sci. 2020, 110631. https://doi.org/10.1016/j.plantsci.2020.110631.

10. Gianfagna, T. Natural and Synthetic Growth Regulators and Their Use in Horticultural and Agronomic Crops. In Plant Hormones: Physiology, Biochemistry and Molecular Biology; Davies, P. J., Ed.; Springer Netherlands: Dordrecht, 1995; pp 751–773. https://doi.org/10.1007/978-94-011-0473-9_34.

11. Rademacher, W. Plant Growth Regulators: Backgrounds and Uses in Plant Production. J. Plant Growth Regul. 2015, 34 (4), 845–872. https://doi.org/10.1007/s00344-015-9541-6.

12. Busi, R.; Goggin, D. E.; Heap, I. M.; Horak, M. J.; Jugulam, M.; Masters, R. A.; Napier, R. M.; Riar, D. S.; Satchivi, N. M.; Torra, J.; Westra, P.; Wright, T. R. Weed Resistance to Synthetic Auxin Herbicides. Pest Manag. Sci. 2018, 74 (10), 2265–2276. https://doi.org/10.1002/ps.4823.

13. Ritchie, H.; Roser, M. Land Use. Our World Data 2013.

14. Gray, W. M.; Kepinski, S.; Rouse, D.; Leyser, O.; Estelle, M. Auxin Regulates SCF(TIR1)-Dependent Degradation of AUX/IAA Proteins. Nature 2001, 414 (6861), 271–276. https://doi.org/10.1038/35104500.

15. Kepinski, S.; Leyser, O. Auxin-Induced SCFTIR1-Aux/IAA Interaction Involves Stable Modification of the SCFTIR1 Complex. Proc Natl Acad Sci U A 2004, 101 (33), 12381–12386. https://doi.org/10.1073/pnas.0402868101.

16. Long, J. A.; Ohno, C.; Smith, Z. R.; Meyerowitz, E. M. TOPLESS Regulates Apical Embryonic Fate in Arabidopsis. Science 2006, 312 (5779), 1520–1523. https://doi.org/10.1126/science.1123841.

17. Szemenyei, H.; Hannon, M.; Long, J. A. TOPLESS Mediates Auxin-Dependent Transcriptional Repression During Arabidopsis Embryogenesis. Science 2008, 319 (5868), 1384–1386. https://doi.org/10.1126/science.1151461.

18. Dharmasiri, N.; Dharmasiri, S.; Estelle, M. The F-Box Protein TIR1 Is an Auxin Receptor. Nature 2005, 435 (7041), 441–445. https://doi.org/10.1038/nature03543.

19. Tan, X.; Calderon-Villalobos, L. I. A.; Sharon, M.; Zheng, C.; Robinson, C. V.; Estelle, M.; Zheng, N. Mechanism of Auxin Perception by the TIR1 Ubiquitin Ligase. Nature 2007, 446 (7136), 640–645. https://doi.org/10.1038/nature05731.

20. Calderón Villalobos, L. I. A.; Lee, S.; De Oliveira, C.; Ivetac, A.; Brandt, W.; Armitage, L.; Sheard, L. B.; Tan, X.; Parry, G.; Mao, H.; Zheng, N.; Napier, R.; Kepinski, S.; Estelle, M. A Combinatorial TIR1/AFB-Aux/IAA Co-Receptor System for Differential Sensing of Auxin. Nat Chem Biol 2012, 8 (5), 477–485. https://doi.org/10.1038/nchembio.926.

21. Niemeyer, M.; Moreno Castillo, E.; Ihling, C. H.; Iacobucci, C.; Wilde, V.; Hellmuth, A.; Hoehenwarter, W.; Samodelov, S. L.; Zurbriggen, M. D.; Kastritis, P. L.; Sinz, A.; Calderón Villalobos, L. I. A. Flexibility of Intrinsically Disordered Degrons in AUX/IAA Proteins Reinforces Auxin Co-Receptor Assemblies. Nat. Commun. 2020, 11 (1), 2277. https://doi.org/10.1038/s41467-020-16147-2.

22. Ulmasov, T.; Murfett, J.; Hagen, G.; Guilfoyle, T. J. Aux/IAA Proteins Repress Expression of Reporter Genes Containing Natural and Highly Active Synthetic Auxin Response Elements. Plant Cell 1997, 9 (11), 1963–1971. https://doi.org/10.1105/tpc.9.11.1963.

23. Weijers, D.; Benkova, E.; Jäger, K. E.; Schlereth, A.; Hamann, T.; Kientz, M.; Wilmoth, J. C.; Reed, J. W.; Jürgens, G. Developmental Specificity of Auxin Response by Pairs of ARF and Aux/IAA Transcriptional Regulators. EMBO J. 2005, 24 (10), 1874–1885. https://doi.org/10.1038/sj.emboj.7600659.

24. Tiwari, S. B.; Hagen, G.; Guilfoyle, T. J. Aux/IAA Proteins Contain a Potent Transcriptional Repression Domain. Plant Cell 2004, 16 (2), 533–543. https://doi.org/10.1105/tpc.017384.

25. Ulmasov, T.; Hagen, G.; Guilfoyle, T. J. Activation and Repression of Transcription by Auxin-Response Factors. Proc. Natl. Acad. Sci. 1999, 96 (10), 5844–5849. https://doi.org/10.1073/pnas.96.10.5844.

26. Dreher, K. A.; Brown, J.; Saw, R. E.; Callis, J. The Arabidopsis Aux/IAA Protein Family Has Diversified in Degradation and Auxin Responsiveness. Plant Cell 2006, 18 (3), 699–714. https://doi.org/10.1105/tpc.105.039172.

27. Havens, K. A.; Guseman, J. M.; Jang, S. S.; Pierre-Jerome, E.; Bolten, N.; Klavins, E.; Nemhauser, J. L. A Synthetic Approach Reveals Extensive Tunability of Auxin Signaling. Plant Physiol. 2012, 160 (1), 135–142. https://doi.org/10.1104/pp.112.202184.

28. Guseman, J. M.; Hellmuth, A.; Lanctot, A.; Feldman, T. P.; Moss, B. L.; Klavins, E.; Villalobos, L. I. A. C.; Nemhauser, J. L. Auxin-Induced Degradation Dynamics Set the Pace for Lateral Root Development. Development 2015, 142 (5), 905–909. https://doi.org/10.1242/dev.117234.

29. Moss, B. L.; Mao, H.; Guseman, J. M.; Hinds, T. R.; Hellmuth, A.; Kovenock, M.; Noorassa, A.; Lanctot, A.; Villalobos, L. I. A. C.; Zheng, N.; Nemhauser, J. L. Rate Motifs Tune Auxin/Indole-3-Acetic Acid Degradation Dynamics. Plant Physiol. 2015, 169 (1), 803–813. https://doi.org/10.1104/pp.15.00587.

30. Wright, R. C.; Zahler, M. L.; Gerben, S. R.; Nemhauser, J. L. Insights into the Evolution and Function of Auxin Signaling F-Box Proteins in *Arabidopsis Thaliana* Through Synthetic Analysis of Natural Variants. Genetics 2017, 207 (2), 583–591. https://doi.org/10.1534/genetics.117.300092.

31. Wright, R. C.; Moss, B. L.; Nemhauser, J. L. The Systems and Synthetic Biology of Auxin. Cold Spring Harb. Perspect. Biol. 2022, 14 (1), a040071. https://doi.org/10.1101/cshperspect.a040071.

32. Hussain, A.; Ullah, I.; Hasnain, S. Microbial Manipulation of Auxins and Cytokinins in Plants. Methods Mol. Biol. Clifton NJ 2017, 1569, 61–72. https://doi.org/10.1007/978-1-4939-6831-2_4.

33. Keswani, C.; Singh, S. P.; Cueto, L.; García-Estrada, C.; Mezaache-Aichour, S.; Glare, T. R.; Borriss, R.; Singh, S. P.; Blázquez, M. A.; Sansinenea, E. Auxins of Microbial Origin and Their Use in Agriculture. Appl. Microbiol. Biotechnol. 2020, 104 (20), 8549–8565. https://doi.org/10.1007/s00253-020-10890-8.

34. Spaepen, S.; Vanderleyden, J. Auxin and Plant-Microbe Interactions. Cold Spring Harb. Perspect. Biol. 2011, 3 (4), a001438. https://doi.org/10.1101/cshperspect.a001438.

35. Prusty, R.; Grisafi, P.; Fink, G. R. The Plant Hormone Indoleacetic Acid Induces Invasive Growth in *Saccharomyces Cerevisiae*. Proc. Natl. Acad. Sci. 2004, 101 (12), 4153–4157. https://doi.org/10.1073/pnas.0400659101.

36. Rao, R. P.; Hunter, A.; Kashpur, O.; Normanly, J. Aberrant Synthesis of Indole-3-Acetic Acid in Saccharomyces Cerevisiae Triggers Morphogenic Transition, a Virulence Trait of Pathogenic Fungi. Genetics 2010, 185 (1), 211–220. https://doi.org/10.1534/genetics.109.112854.

37. Dong, L.; Ma, Y.; Chen, C.-Y.; Shen, L.; Sun, W.; Cui, G.; Naqvi, N. I.; Deng, Y. Z. Identification and Characterization of Auxin/IAA Biosynthesis Pathway in the Rice Blast Fungus Magnaporthe Oryzae. J. Fungi 2022, 8 (2), 208. https://doi.org/10.3390/jof8020208.

38. Petti, C.; Reiber, K.; Ali, S. S.; Berney, M.; Doohan, F. M. Auxin as a Player in the Biocontrol of Fusarium Head Blight Disease of Barley and Its Potential as a Disease Control Agent. BMC Plant Biol. 2012, 12 (1), 224. https://doi.org/10.1186/1471-2229-12-224.

39. Ferreira-Pinto, M. M.; Moura-Guedes, M. C.; Barreiro, M. G.; Pais, I.; Santos, M. R.; Silva, M. J. Aureobasidium Pullulansas a Biocontrol Agent of Blue Mold in “Rocha” Pear. Commun. Agric. Appl. Biol. Sci. 2006, 71 (3 Pt B), 973–978.

40. Di Canito, A.; Mateo-Vargas, M. A.; Mazzieri, M.; Cantoral, J.; Foschino, R.; Cordero-Bueso, G.; Vigentini, I. The Role of Yeasts as Biocontrol Agents for Pathogenic Fungi on Postharvest Grapes: A Review. Foods 2021, 10 (7), 1650. https://doi.org/10.3390/foods10071650.

41. Di Francesco, A.; Di Foggia, M.; Corbetta, M.; Baldo, D.; Ratti, C.; Baraldi, E. Biocontrol Activity and Plant Growth Promotion Exerted by Aureobasidium Pullulans Strains. J. Plant Growth Regul. 2020. https://doi.org/10.1007/s00344-020-10184-3.

42. Yu, T.; Zheng, X. D. Indole-3-Acetic Acid Enhances the Biocontrol of Penicillium Expansum and Botrytis Cinerea on Pear Fruit by Cryptococcus Laurentii. FEMS Yeast Res. 2007, 7 (3), 459–464. https://doi.org/10.1111/j.1567-1364.2006.00171.x.

43. Rahman, M.; Rahman, M.; Islam, T. Improving Yield and Antioxidant Properties of Strawberries by Utilizing Microbes and Natural Products. Strawb. - Pre-Post-Harvest Manag. Tech. High. Fruit Qual. 2019. https://doi.org/10.5772/intechopen.84803.

44. Figueroa, C. R.; Opazo, M. C.; Vera, P.; Arriagada, O.; Díaz, M.; Moya-León, M. A. Effect of Postharvest Treatment of Calcium and Auxin on Cell Wall Composition and Expression of Cell Wall-Modifying Genes in the Chilean Strawberry (Fragaria Chiloensis) Fruit. Food Chem. 2012, 132 (4), 2014–2022. https://doi.org/10.1016/j.foodchem.2011.12.041.

45. Wang, L.; Dou, G.; Guo, H.; Zhang, Q.; Qin, X.; Yu, W.; Jiang, C.; Xiao, H. Volatile Organic Compounds of Hanseniaspora Uvarum Increase Strawberry Fruit Flavor and Defense during Cold Storage. Food Sci. Nutr. 2019, 7 (8), 2625–2635. https://doi.org/10.1002/fsn3.1116.

46. Breakspear, A.; Liu, C.; Roy, S.; Stacey, N.; Rogers, C.; Trick, M.; Morieri, G.; Mysore, K. S.; Wen, J.; Oldroyd, G. E. D.; Downie, J. A.; Murray, J. D. The Root Hair “Infectome” of Medicago Truncatula Uncovers Changes in Cell Cycle Genes and Reveals a Requirement for Auxin Signaling in Rhizobial Infection. Plant Cell 2014. https://doi.org/10.1105/tpc.114.133496.

47. Laplaze, L.; Lucas, M.; Champion, A. Rhizobial Root Hair Infection Requires Auxin Signaling. Trends Plant Sci. 2015, 20 (6), 332–334. https://doi.org/10.1016/j.tplants.2015.04.004.

48. Chen, X.; Chen, J.; Liao, D.; Ye, H.; Li, C.; Luo, Z.; Yan, A.; Zhao, Q.; Xie, K.; Li, Y.; Wang, D.; Chen, J.; Chen, A.; Xu, G. Auxin-Mediated Regulation of Arbuscular Mycorrhizal Symbiosis: A Role of SlGH3.4 in Tomato. Plant Cell Environ. n/a (n/a). https://doi.org/10.1111/pce.14210.

49. Etemadi, M.; Gutjahr, C.; Couzigou, J.-M.; Zouine, M.; Lauressergues, D.; Timmers, A.; Audran, C.; Bouzayen, M.; Bécard, G.; Combier, J.-P. Auxin Perception Is Required for Arbuscule Development in Arbuscular Mycorrhizal Symbiosis1[W]. Plant Physiol. 2014, 166 (1), 281–292. https://doi.org/10.1104/pp.114.246595.

50. Liu, Y.-Y.; Chen, H.-W.; Chou, J.-Y. Variation in Indole-3-Acetic Acid Production by Wild Saccharomyces Cerevisiae and S. Paradoxus Strains from Diverse Ecological Sources and Its Effect on Growth. PloS One 2016, 11 (8), e0160524. https://doi.org/10.1371/journal.pone.0160524.

51. Kunkel, B. N.; Johnson, J. M. B. Auxin Plays Multiple Roles during Plant-Pathogen Interactions. Cold Spring Harb. Perspect. Biol. 2021, a040022. https://doi.org/10.1101/cshperspect.a040022.

52. Morffy, N.; Strader, L. C. Old Town Roads: Routes of Auxin Biosynthesis across Kingdoms. Curr. Opin. Plant Biol. 2020, 55, 21–27. https://doi.org/10.1016/j.pbi.2020.02.002.

53. Novák, O.; Floková, K. An UHPLC-MS/MS Method for Target Profiling of Stress-Related Phytohormones. In Plant Metabolomics; António, C., Ed.; Springer New York: New York, NY, 2018; Vol. 1778, pp 183–192. https://doi.org/10.1007/978-1-4939-7819-9_13.

54. Peres, L. E. P.; Mercier, H.; Kerbauy, G. B.; Zaffari, G. R. Endogenous levels of IAA, cytokinins and ABA in a shootless orchid and a rootless bromeliad determined by means of HPLC and ELISA. Rev. Bras. Fisiol. Veg. Braz. 1997.

55. Rivier, L. GC-MS of Auxins. In Gas Chromatography/Mass Spectrometry; Linskens, H. F., Jackson, J. F., Eds.; Modern Methods of Plant Analysis; Springer: Berlin, Heidelberg, 1986; pp 146–188. https://doi.org/10.1007/978-3-642-82612-2_8.

56. Pence, V. C.; Caruso, J. L. Elisa Determination of IAA Using Antibodies against Ring-Linked IAA. Phytochemistry 1987, 26 (5), 1251–1255. https://doi.org/10.1016/S0031-9422(00)81791-4.

57. Šimura, J.; Antoniadi, I.; Široká, J.; Tarkowská, D.; Strnad, M.; Ljung, K.; Novák, O. Plant Hormonomics: Multiple Phytohormone Profiling by Targeted Metabolomics. Plant Physiol. 2018, 177 (2), 476–489. https://doi.org/10.1104/pp.18.00293.

58. Wright, R. C.; Nemhauser, J. Plant Synthetic Biology: Quantifying the “Known Unknowns” and Discovering the “Unknown Unknowns”1[OPEN]. Plant Physiol. 2019, 179 (3), 885–893. https://doi.org/10.1104/pp.18.01222.

59. Alexandrov, K.; Vickers, C. E. In Vivo Protein-Based Biosensors: Seeing Metabolism in Real Time. Trends Biotechnol. 2022. https://doi.org/10.1016/j.tibtech.2022.07.002.

60. Fowler, D. M.; Araya, C. L.; Fleishman, S. J.; Kellogg, E. H.; Stephany, J. J.; Baker, D.; Fields, S. High-Resolution Mapping of Protein Sequence-Function Relationships. Nat. Methods 2010, 7 (9), 741–746. https://doi.org/10.1038/nmeth.1492.

61. Francisco, J. A.; Campbell, R.; Iverson, B. L.; Georgiou, G. Production and Fluorescence-Activated Cell Sorting of Escherichia Coli Expressing a Functional Antibody Fragment on the External Surface. Proc. Natl. Acad. Sci. U. S. A. 1993, 90 (22), 10444–10448.

62. Fowler, D. M.; Fields, S. Deep Mutational Scanning: A New Style of Protein Science. Nat. Methods 2014, 11 (8), 801–807. https://doi.org/10.1038/nmeth.3027.

63. Shin, H.; Cho, B.-K. Rational Protein Engineering Guided by Deep Mutational Scanning. Int. J. Mol. Sci. 2015, 16 (9), 23094–23110. https://doi.org/10.3390/ijms160923094.

64. Alley, E. C.; Khimulya, G.; Biswas, S.; AlQuraishi, M.; Church, G. M. Unified Rational Protein Engineering with Sequence-Based Deep Representation Learning. Nat. Methods 2019, 16 (12), 1315–1322. https://doi.org/10.1038/s41592-019-0598-1.

65. Gelman, S.; Fahlberg, S. A.; Heinzelman, P.; Romero, P. A.; Gitter, A. Neural Networks to Learn Protein Sequence–Function Relationships from Deep Mutational Scanning Data. Proc. Natl. Acad. Sci. 2021, 118 (48). https://doi.org/10.1073/pnas.2104878118.

66. Romero, P. A.; Arnold, F. H. Exploring Protein Fitness Landscapes by Directed Evolution. Nat. Rev. Mol. Cell Biol. 2009, 10 (12), 866–876. https://doi.org/10.1038/nrm2805.

67. Packer, M. S.; Liu, D. R. Methods for the Directed Evolution of Proteins. Nat. Rev. Genet. 2015, advance online publication. https://doi.org/10.1038/nrg3927.

68. Dharmasiri, N.; Dharmasiri, S.; Weijers, D.; Lechner, E.; Yamada, M.; Hobbie, L.; Ehrismann, J. S.; Jürgens, G.; Estelle, M. Plant Development Is Regulated by a Family of Auxin Receptor F Box Proteins. Dev. Cell 2005, 9 (1), 109–119. https://doi.org/10.1016/j.devcel.2005.05.014.

69. Tripathi, D. K.; Yadav, S. R.; Mochida, K.; Tran, L.-S. P. Plant Growth Regulators: True Managers of Plant Life. Plant Cell Physiol. 2022, 63 (12), 1757–1760. https://doi.org/10.1093/pcp/pcac170.

70. Zimran, G.; Feuer, E.; Pri-Tal, O.; Shpilman, M.; Mosquna, A. Directed Evolution of Herbicide Biosensors in a Fluorescence-Activated Cell-Sorting-Compatible Yeast Two-Hybrid Platform. ACS Synth. Biol. 2022, acssynbio.2c00297. https://doi.org/10.1021/acssynbio.2c00297.

71. Nishimura Kohei; Kanemaki Masato T. Rapid Depletion of Budding Yeast Proteins via the Fusion of an Auxin-Inducible Degron (AID). Curr. Protoc. Cell Biol. 2014, 64 (1), 20.9.1–20.9.16. https://doi.org/10.1002/0471143030.cb2009s64.

72. Zhang, L.; Ward, J. D.; Cheng, Z.; Dernburg, A. F. The Auxin-Inducible Degradation (AID) System Enables Versatile Conditional Protein Depletion in *C*. Elegans. Dev. Camb. Engl. 2015, 142 (24), 4374–4384. https://doi.org/10.1242/dev.129635.

73. Brown, K. M.; Long, S.; Sibley, L. D. Conditional Knockdown of Proteins Using Auxin-Inducible Degron (AID) Fusions in Toxoplasma Gondii. Bio-Protoc. 2018, 8 (4). https://doi.org/10.21769/BioProtoc.2728.

74. Miura, K.; Matoba, S.; Ogonuki, N.; Namiki, T.; Ito, J.; Kashiwazaki, N.; Ogura, A. Application of Auxin-Inducible Degron Technology to Mouse Oocyte Activation with PLCζ. J. Reprod. Dev. 2018. https://doi.org/10.1262/jrd.2018-053.

75. Li, S.; Prasanna, X.; Salo, V. T.; Vattulainen, I.; Ikonen, E. An Efficient Auxin-Inducible Degron System with Low Basal Degradation in Human Cells. Nat. Methods 2019, 16 (9), 866–869. https://doi.org/10.1038/s41592-019-0512-x.

76. Nishimura, K.; Yamada, R.; Hagihara, S.; Iwasaki, R.; Uchida, N.; Kamura, T.; Takahashi, K.; Torii, K. U.; Fukagawa, T. A Super-Sensitive Auxin-Inducible Degron System with an Engineered Auxin-TIR1 Pair. Nucleic Acids Res. 2020, 48 (18), e108–e108. https://doi.org/10.1093/nar/gkaa748.

77. Friml, J.; Vieten, A.; Sauer, M.; Weijers, D.; Schwarz, H.; Hamann, T.; Offringa, R.; Jürgens, G. Efflux-Dependent Auxin Gradients Establish the Apical-Basal Axis of Arabidopsis. Nature 2003, 426 (6963), 147–153. https://doi.org/10.1038/nature02085.

78. Liao, C.-Y.; Smet, W.; Brunoud, G.; Yoshida, S.; Vernoux, T.; Weijers, D. Reporters for Sensitive and Quantitative Measurement of Auxin Response. Nat. Methods 2015, 12 (3), 207–210. https://doi.org/10.1038/nmeth.3279.

79. Weijers, D.; Schlereth, A.; Ehrismann, J. S.; Schwank, G.; Kientz, M.; Jürgens, G. Auxin Triggers Transient Local Signaling for Cell Specification in Arabidopsis Embryogenesis. Dev. Cell 2006, 10 (2), 265–270. https://doi.org/10.1016/j.devcel.2005.12.001.

80. Brunoud, G.; Wells, D. M.; Oliva, M.; Larrieu, A.; Mirabet, V.; Burrow, A. H.; Beeckman, T.; Kepinski, S.; Traas, J.; Bennett, M. J.; Vernoux, T. A Novel Sensor to Map Auxin Response and Distribution at High Spatio-Temporal Resolution. Nature 2012, 482 (7383), 103–106. https://doi.org/10.1038/nature10791.

81. Galvan-Ampudia, C. S.; Cerutti, G.; Legrand, J.; Brunoud, G.; Martin-Arevalillo, R.; Azais, R.; Bayle, V.; Moussu, S.; Wenzl, C.; Jaillais, Y.; Lohmann, J. U.; Godin, C.; Vernoux, T. Temporal Integration of Auxin Information for the Regulation of Patterning. eLife 2020, 9, e55832. https://doi.org/10.7554/eLife.55832.

82. Wend, S.; Bosco, C. D.; Kämpf, M. M.; Ren, F.; Palme, K.; Weber, W.; Dovzhenko, A.; Zurbriggen, M. D. A Quantitative Ratiometric Sensor for Time-Resolved Analysis of Auxin Dynamics. Sci. Rep. 2013, 3 (1), 2052. https://doi.org/10.1038/srep02052.

83. Herud-Sikimić, O.; Stiel, A. C.; Kolb, M.; Shanmugaratnam, S.; Berendzen, K. W.; Feldhaus, C.; Höcker, B.; Jürgens, G. A Biosensor for the Direct Visualization of Auxin. Nature 2021, 592 (7856), 768–772. https://doi.org/10.1038/s41586-021-03425-2.

84. Dong, L.; Shen, Q.; Chen, C.-Y.; Shen, L.; Yang, F.; Naqvi, N. I.; Deng, Y. Z. Fungal Auxin Is a Quorum-Based Modulator of Blast Disease Severity. Cold Spring Harbor Laboratory October 26, 2021, p 2021.10.26.465851. https://doi.org/10.1101/2021.10.26.465851.

85. Pierre-Jerome, E.; Wright, R. C.; Nemhauser, J. L. Characterizing Auxin Response Circuits in Saccharomyces Cerevisiae by Flow Cytometry. In Plant Hormones: Methods and Protocols; Kleine-Vehn, J., Sauer, M., Eds.; Methods in Molecular Biology; Springer: New York, NY, 2017; pp 271–281. https://doi.org/10.1007/978-1-4939-6469-7_22.

86. Souza-Moreira, T. M.; Navarrete, C.; Chen, X.; Zanelli, C. F.; Valentini, S. R.; Furlan, M.; Nielsen, J.; Krivoruchko, A. Screening of 2A Peptides for Polycistronic Gene Expression in Yeast. FEMS Yeast Res. 2018, 18 (5). https://doi.org/10.1093/femsyr/foy036.

87. Zenser, N.; Ellsmore, A.; Leasure, C.; Callis, J. Auxin Modulates the Degradation Rate of Aux/IAA Proteins. Proc. Natl. Acad. Sci. 2001, 98 (20), 11795–11800. https://doi.org/10.1073/pnas.211312798.

88. Parry, G.; Calderon-Villalobos, L. I.; Prigge, M.; Peret, B.; Dharmasiri, S.; Itoh, H.; Lechner, E.; Gray, W. M.; Bennett, M.; Estelle, M. Complex Regulation of the TIR1/AFB Family of Auxin Receptors. Proc. Natl. Acad. Sci. 2009, 106 (52), 22540–22545. https://doi.org/10.1073/pnas.0911967106.

89. Xiong, L.; Zeng, Y.; Tang, R.-Q.; Alper, H. S.; Bai, F.-W.; Zhao, X.-Q. Condition-Specific Promoter Activities in Saccharomyces Cerevisiae. Microb. Cell Factories 2018, 17 (1), 58. https://doi.org/10.1186/s12934-018-0899-6.

90. Lee, M. E.; DeLoache, W. C.; Cervantes, B.; Dueber, J. E. A Highly Characterized Yeast Toolkit for Modular, Multipart Assembly. ACS Synth. Biol. 2015, 4 (9), 975–986. https://doi.org/10.1021/sb500366v.

91. Galan, J.-M.; Peter, M. Ubiquitin-Dependent Degradation of Multiple F-Box Proteins by an Autocatalytic Mechanism. Proc. Natl. Acad. Sci. 1999, 96 (16), 9124–9129. https://doi.org/10.1073/pnas.96.16.9124.

92. Zhou, P.; Howley, P. M. Ubiquitination and Degradation of the Substrate Recognition Subunits of SCF Ubiquitin–Protein Ligases. Mol. Cell 1998, 2 (5), 571–580. https://doi.org/10.1016/S1097-2765(00)80156-2.

93. Yu, H.; Zhang, Y.; Moss, B. L.; Bargmann, B. O. R.; Wang, R.; Prigge, M.; Nemhauser, J. L.; Estelle, M. Untethering the TIR1 Auxin Receptor from the SCF Complex Increases Its Stability and Inhibits Auxin Response. Nat. Plants 2015, 1 (3), 14030. https://doi.org/10.1038/nplants.2014.30.

94. Johnston, M.; Davis, R. W. Sequences That Regulate the Divergent GAL1-GAL10 Promoter in Saccharomyces Cerevisiae. Mol. Cell. Biol. 1984, 4 (8), 1440–1448. https://doi.org/10.1128/mcb.4.8.1440-1448.1984.

95. Sikorski, R. S.; Hieter, P. A System of Shuttle Vectors and Yeast Host Strains Designed for Efficient Manipulation of DNA in Saccharomyces Cerevisiae. Genetics 1989, 122 (1), 19–27.

96. Sobel, S. G.; Wolin, S. L. Two Yeast La Motif-Containing Proteins Are RNA-Binding Proteins That Associate with Polyribosomes. Mol. Biol. Cell 1999, 10 (11), 3849–3862.

97. Ralser, M.; Kuhl, H.; Ralser, M.; Werber, M.; Lehrach, H.; Breitenbach, M.; Timmermann, B. The Saccharomyces Cerevisiae W303-K6001 Cross-Platform Genome Sequence: Insights into Ancestry and Physiology of a Laboratory Mutt. Open Biol. 2012, 2 (8), 120093. https://doi.org/10.1098/rsob.120093.

98. Matheson, K.; Parsons, L.; Gammie, A. Whole-Genome Sequence and Variant Analysis of W303, a Widely-Used Strain of Saccharomyces Cerevisiae. G3 GenesGenomesGenetics 2017, 7 (7), 2219–2226. https://doi.org/10.1534/g3.117.040022.

99. Novák, O.; Hényková, E.; Sairanen, I.; Kowalczyk, M.; Pospíšil, T.; Ljung, K. Tissue-Specific Profiling of the Arabidopsis Thaliana Auxin Metabolome. *Plant J*. Cell Mol. Biol. 2012, 72 (3), 523–536. https://doi.org/10.1111/j.1365-313X.2012.05085.x.

100. Petersson, S. V.; Johansson, A. I.; Kowalczyk, M.; Makoveychuk, A.; Wang, J. Y.; Moritz, T.; Grebe, M.; Benfey, P. N.; Sandberg, G.; Ljung, K. An Auxin Gradient and Maximum in the Arabidopsis Root Apex Shown by High-Resolution Cell-Specific Analysis of IAA Distribution and Synthesis. Plant Cell 2009, 21 (6), 1659–1668. https://doi.org/10.1105/tpc.109.066480.

101. Nicastro, R.; Raucci, S.; Michel, A. H.; Stumpe, M.; Osuna, G. M. G.; Jaquenoud, M.; Kornmann, B.; Virgilio, C. D. Indole-3-Acetic Acid Is a Physiological Inhibitor of TORC1 in Yeast. PLOS Genet. 2021, 17 (3), e1009414. https://doi.org/10.1371/journal.pgen.1009414.

102. Reinhardt, D.; Mandel, T.; Kuhlemeier, C. Auxin Regulates the Initiation and Radial Position of Plant Lateral Organs. Plant Cell 2000, 12 (4), 507–518. https://doi.org/10.1105/tpc.12.4.507.

103. Morawska, M.; Ulrich, H. D. An Expanded Tool Kit for the Auxin-Inducible Degron System in Budding Yeast. Yeast 2013, 30 (9), 341–351. https://doi.org/10.1002/yea.2967.

104. Khakhar, A.; Bolten, N. J.; Nemhauser, J.; Klavins, E. Cell–Cell Communication in Yeast Using Auxin Biosynthesis and Auxin Responsive CRISPR Transcription Factors. ACS Synth. Biol. 2016, 5 (4), 279–286. https://doi.org/10.1021/acssynbio.5b00064.

105. Snyder, N. A.; Kim, A.; Kester, L.; Gale, A. N.; Studer, C.; Hoepfner, D.; Roggo, S.; Helliwell, S. B.; Cunningham, K. W. Auxin-Inducible Depletion of the Essentialome Suggests Inhibition of TORC1 by Auxins and Inhibition of Vrg4 by SDZ 90-215, a Natural Antifungal Cyclopeptide. G3 GenesGenomesGenetics 2019, 9 (3), 829–840. https://doi.org/10.1534/g3.118.200748.

106. Ravikumar, A.; Arrieta, A.; Liu, C. C. An Orthogonal DNA Replication System in Yeast. Nat. Chem. Biol. 2014, 10 (3), 175–177. https://doi.org/10.1038/nchembio.1439.

107. Ravikumar, A.; Arzumanyan, G. A.; Obadi, M. K. A.; Javanpour, A. A.; Liu, C. C. Scalable, Continuous Evolution of Genes at Mutation Rates above Genomic Error Thresholds. Cell 2018, 175 (7), 1946–1957.e13. https://doi.org/10.1016/j.cell.2018.10.021.

108. Drummond, D. A.; Raval, A.; Wilke, C. O. A Single Determinant Dominates the Rate of Yeast Protein Evolution. Mol. Biol. Evol. 2006, 23 (2), 327–337. https://doi.org/10.1093/molbev/msj038.

109. Bunsangiam, S.; Sakpuntoon, V.; Srisuk, N.; Ohashi, T.; Fujiyama, K.; Limtong, S. Biosynthetic Pathway of Indole-3-Acetic Acid in Basidiomycetous Yeast Rhodosporidiobolus Fluvialis. Mycobiology 2019, 47 (3), 292–300. https://doi.org/10.1080/12298093.2019.1638672.

110. Nakamura, T.; Murakami, T.; Saotome, M.; Tomita, K.; Kitsuwa, T.; Meyers, S. P. Identification of Indole-3-Acetic Acid in Pichia Spartinae, an Ascosporogenous Yeast from Spartina Alterniflora Marshland Environments. Mycologia 1991, 83 (5), 662–664. https://doi.org/10.2307/3760223.

111. Nutaratat, P.; Srisuk, N.; Arunrattiyakorn, P.; Limtong, S. Indole-3-Acetic Acid Biosynthetic Pathways in the Basidiomycetous Yeast Rhodosporidium Paludigenum. Arch. Microbiol. 2016, 198 (5), 429–437. https://doi.org/10.1007/s00203-016-1202-z.

112. Xin, G.; Glawe, D.; Doty, S. L. Characterization of Three Endophytic, Indole-3-Acetic Acid-Producing Yeasts Occurring in Populus Trees. Mycol. Res. 2009, 113 (9), 973–980. https://doi.org/10.1016/j.mycres.2009.06.001.

113. Mihaljevic ulj, M.; Tomaz, I.; Maslov Bandić, L.; Puhelek, I.; Jagatić Korenika, A. M.; Jeromel, A. Influence of Different Yeast Strains on Metabolism of Tryptophan and Indole-3-Acetic Acid during Fermentation. South Afr. J. Enol. Vitic. 2015, 36 (1), 44–49.

114. Kumla, J.; Suwannarach, N.; Matsui, K.; Lumyong, S. Biosynthetic Pathway of Indole-3-Acetic Acid in Ectomycorrhizal Fungi Collected from Northern Thailand. PLOS ONE 2020, 15 (1), e0227478. https://doi.org/10.1371/journal.pone.0227478.

115. Perelta, A. Identifying the Molecular Mechanism of Indole-3-Acetic Acid Detection in the Fungi Saccharomyces Cerevisiae and Candida Albicans. Masters Theses Theses Years 2012.

116. Yesbolatova, A.; Saito, Y.; Kitamoto, N.; Makino-Itou, H.; Ajima, R.; Nakano, R.; Nakaoka, H.; Fukui, K.; Gamo, K.; Tominari, Y.; Takeuchi, H.; Saga, Y.; Hayashi, K.; Kanemaki, M. T. The Auxin-Inducible Degron 2 Technology Provides Sharp Degradation Control in Yeast, Mammalian Cells, and Mice. Nat. Commun. 2020, 11 (1), 5701. https://doi.org/10.1038/s41467-020-19532-z.

117. Liachko, I.; Dunham, M. J. An Autonomously Replicating Sequence for Use in a Wide Range of Budding Yeasts. FEMS Yeast Res. 2014, 14 (2), 364–367. https://doi.org/10.1111/1567-1364.12123.

118. Hayashi, K.; Neve, J.; Hirose, M.; Kuboki, A.; Shimada, Y.; Kepinski, S.; Nozaki, H. Rational Design of an Auxin Antagonist of the SCF(TIR1) Auxin Receptor Complex. ACS Chem. Biol. 2012, 7 (3), 590–598. https://doi.org/10.1021/cb200404c.

119. Prigge, M. J.; Platre, M.; Kadakia, N.; Zhang, Y.; Greenham, K.; Szutu, W.; Pandey, B. K.; Bhosale, R. A.; Bennett, M. J.; Busch, W.; Estelle, M. Genetic Analysis of the Arabidopsis TIR1/AFB Auxin Receptors Reveals Both Overlapping and Specialized Functions. eLife 2020, 9, e54740. https://doi.org/10.7554/eLife.54740.

120. Iglesias, M. J.; Terrile, M. C.; Correa-Aragunde, N.; Colman, S. L.; Izquierdo-Álvarez, A.; Fiol, D. F.; París, R.; Sánchez-López, N.; Marina, A.; Calderón Villalobos, L. I. A.; Estelle, M.; Lamattina, L.; Martínez-Ruiz, A.; Casalongué, C. A. Regulation of SCFTIR1/AFBs E3 Ligase Assembly by S-Nitrosylation of Arabidopsis SKP1-Like1 Impacts on Auxin Signaling. Redox Biol. 2018, 18, 200–210. https://doi.org/10.1016/j.redox.2018.07.003.

121. Laha, N. P.; Giehl, R. F. H.; Riemer, E.; Qiu, D.; Pullagurla, N. J.; Schneider, R.; Dhir, Y. W.; Yadav, R.; Mihiret, Y. E.; Gaugler, P.; Gaugler, V.; Mao, H.; Zheng, N.; von Wirén, N.; Saiardi, A.; Bhattacharjee, S.; Jessen, H. J.; Laha, D.; Schaaf, G. INOSITOL (1,3,4) TRIPHOSPHATE 5/6 KINASE1-Dependent Inositol Polyphosphates Regulate Auxin Responses in Arabidopsis. Plant Physiol. 2022, kiac425. https://doi.org/10.1093/plphys/kiac425.

122. Fendrych, M.; Leung, J.; Friml, J. TIR1/AFB-Aux/IAA Auxin Perception Mediates Rapid Cell Wall Acidification and Growth of Arabidopsis Hypocotyls. eLife 2016, 5, e19048. https://doi.org/10.7554/eLife.19048.

123. Chen, H.; Li, L.; Qi, L.; Friml, J. Distinct Functions of TIR1 and AFB1 Receptors in Auxin Signalling. bioRxiv January 6, 2023, p 2023.01.05.522749. https://doi.org/10.1101/2023.01.05.522749.

124. Dubey, S. M.; Han, S.; Stutzman, N.; Prigge, M. J.; Medvecká, E.; Platre, M. P.; Busch, W.; Fendrych, M.; Estelle, M. The AFB1 Auxin Receptor Controls the Cytoplasmic Auxin Response Pathway in Arabidopsis Thaliana. bioRxiv January 4, 2023, p 2023.01.04.522696. https://doi.org/10.1101/2023.01.04.522696.

125. Dezfulian, M. H.; Jalili, E.; Roberto, D. K. A.; Moss, B. L.; Khoo, K.; Nemhauser, J. L.; Crosby, W. L. Oligomerization of SCF TIR1 Is Essential for Aux/IAA Degradation and Auxin Signaling in *Arabidopsis*. PLOS Genet 2016, 12 (9), e1006301. https://doi.org/10.1371/journal.pgen.1006301.

126. Uchida, N.; Takahashi, K.; Iwasaki, R.; Yamada, R.; Yoshimura, M.; Endo, T. A.; Kimura, S.; Zhang, H.; Nomoto, M.; Tada, Y.; Kinoshita, T.; Itami, K.; Hagihara, S.; Torii, K. U. Chemical Hijacking of Auxin Signaling with an Engineered Auxin–TIR1 Pair. Nat. Chem. Biol. 2018, 14 (3), 299–305. https://doi.org/10.1038/nchembio.2555.

127. Jing, H.; Yang, X.; Feng, J.; Zhang, J.; Strader, L. C.; Zuo, J. S-Nitrosylation of Aux/IAA Protein Represses Auxin Signaling. bioRxiv October 7, 2022, p 2022.10.07.511298. https://doi.org/10.1101/2022.10.07.511298.

128. Jing, H.; Yang, X.; Zhang, J.; Liu, X.; Zheng, H.; Dong, G.; Nian, J.; Feng, J.; Xia, B.; Qian, Q.; Li, J.; Zuo, J. Peptidyl-Prolyl Isomerization Targets Rice Aux/IAAs for Proteasomal Degradation during Auxin Signalling. Nat. Commun. 2015, 6, 7395. https://doi.org/10.1038/ncomms8395.

129. Enders Tara A.; Frick Elizabeth M.; Strader Lucia C. An Arabidopsis Kinase Cascade Influences Auxin-responsive Cell Expansion. Plant J. 2017, 92 (1), 68–81. https://doi.org/10.1111/tpj.13635.

130. Figueiredo, M. R. A. de; Strader, L. C. Intrinsic and Extrinsic Regulators of Aux/IAA Protein Degradation Dynamics. Trends Biochem. Sci. 2022, S0968000422001451. https://doi.org/10.1016/j.tibs.2022.06.004.

131. Tao, S.; Estelle, M. Mutational Studies of the Aux/IAA Proteins in Physcomitrella Reveal Novel Insights into Their Function. New Phytol. 218 (4), 1534–1542. https://doi.org/10.1111/nph.15039.

132. de Figueiredo, M. R. A.; Küpper, A.; Malone, J. M.; Petrovic, T.; de Figueiredo, A. B. T. B.; Campagnola, G.; Peersen, O. B.; Prasad, K. V. S. K.; Patterson, E. L.; Reddy, A. S. N.; Kubeš, M. F.; Napier, R.; Preston, C.; Gaines, T. A. An In-Frame Deletion Mutation in the Degron Tail of Auxin Co-Receptor IAA2 Confers Resistance to the Herbicide 2,4-D in Sisymbrium Orientale; preprint; Plant Biology, 2021. https://doi.org/10.1101/2021.03.04.433944.

133. Harborough, S. R.; Kalverda, A. P.; Thompson, G. S.; Kieffer, M. L.; Kubes, M.; Quareshy, M.; Uzanova, V.; Prusinska, J.; Hayashi, K.-I.; Napier, R. M.; Manfield, I. W.; Kepinski, S. A Fuzzy Encounter Complex Precedes Formation of the Fully-Engaged TIR1-Aux/IAA Auxin Co-Receptor System. bioRxiv 2019, 781922. https://doi.org/10.1101/781922.

134. Shabek, N.; Zheng, N. Plant Ubiquitin Ligases as Signaling Hubs. Nat. Struct. Mol. Biol. 2014, 21 (4), 293–296. https://doi.org/10.1038/nsmb.2804.

135. Thines, B.; Katsir, L.; Melotto, M.; Niu, Y.; Mandaokar, A.; Liu, G.; Nomura, K.; He, S. Y.; Howe, G. A.; Browse, J. JAZ Repressor Proteins Are Targets of the SCF(COI1) Complex during Jasmonate Signalling. Nature 2007, 448 (7154), 661–665. https://doi.org/10.1038/nature05960.

136. Sheard, L. B.; Tan, X.; Mao, H.; Withers, J.; Ben-Nissan, G.; Hinds, T. R.; Kobayashi, Y.; Hsu, F.-F.; Sharon, M.; Browse, J.; He, S. Y.; Rizo, J.; Howe, G. A.; Zheng, N. Jasmonate Perception by Inositol Phosphate-Potentiated COI1-JAZ Co-Receptor. Nature 2010, 468 (7322), 400–405. https://doi.org/10.1038/nature09430.

137. Dill, A.; Thomas, S. G.; Hu, J.; Steber, C. M.; Sun, T. The Arabidopsis F-Box Protein SLEEPY1 Targets Gibberellin Signaling Repressors for Gibberellin-Induced Degradation. Plant Cell 2004, 16 (6), 1392–1405. https://doi.org/10.1105/tpc.020958.

138. Hirano, K.; Asano, K.; Tsuji, H.; Kawamura, M.; Mori, H.; Kitano, H.; Ueguchi-Tanaka, M.; Matsuoka, M. Characterization of the Molecular Mechanism Underlying Gibberellin Perception Complex Formation in Rice. Plant Cell 2010, 22 (8), 2680–2696. https://doi.org/10.1105/tpc.110.075549.

139. Murase, K.; Hirano, Y.; Sun, T.; Hakoshima, T. Gibberellin-Induced DELLA Recognition by the Gibberellin Receptor GID1. Nature 2008, 456 (7221), 459–463. https://doi.org/10.1038/nature07519.

140. Shimada, A.; Ueguchi-Tanaka, M.; Nakatsu, T.; Nakajima, M.; Naoe, Y.; Ohmiya, H.; Kato, H.; Matsuoka, M. Structural Basis for Gibberellin Recognition by Its Receptor GID1. Nature 2008, 456 (7221), 520–523. https://doi.org/10.1038/nature07546.

141. Shabek, N.; Ticchiarelli, F.; Mao, H.; Hinds, T. R.; Leyser, O.; Zheng, N. Structural Plasticity of D3–D14 Ubiquitin Ligase in Strigolactone Signalling. Nature 2018, 563 (7733), 652–656. https://doi.org/10.1038/s41586-018-0743-5.

142. Stirnberg, P.; van De Sande, K.; Leyser, H. M. O. MAX1 and MAX2 Control Shoot Lateral Branching in Arabidopsis. Dev. Camb. Engl. 2002, 129 (5), 1131–1141. https://doi.org/10.1242/dev.129.5.1131.

143. Nelson, D. C.; Scaffidi, A.; Dun, E. A.; Waters, M. T.; Flematti, G. R.; Dixon, K. W.; Beveridge, C. A.; Ghisalberti, E. L.; Smith, S. M. F-Box Protein MAX2 Has Dual Roles in Karrikin and Strigolactone Signaling in Arabidopsis Thaliana. Proc. Natl. Acad. Sci. 2011, 108 (21), 8897–8902. https://doi.org/10.1073/pnas.1100987108.

144. Guo, Y.; Zheng, Z.; La Clair, J. J.; Chory, J.; Noel, J. P. Smoke-Derived Karrikin Perception by the α/β-Hydrolase KAI2 from Arabidopsis. Proc. Natl. Acad. Sci. 2013, 110 (20), 8284–8289. https://doi.org/10.1073/pnas.1306265110.

145. Spaepen, S. Plant Hormones Produced by Microbes. In Principles of Plant-Microbe Interactions: Microbes for Sustainable Agriculture; Lugtenberg, B., Ed.; Springer International Publishing: Cham, 2015; pp 247–256. https://doi.org/10.1007/978-3-319-08575-3_26.

146. Khosla, A.; Rodriguez-Furlan, C.; Kapoor, S.; Van Norman, J. M.; Nelson, D. C. A Series of Dual-Reporter Vectors for Ratiometric Analysis of Protein Abundance in Plants. Plant Direct 2020, 4 (6), e00231. https://doi.org/10.1002/pld3.231.

147. Hillson, N. J.; Rosengarten, R. D.; Keasling, J. D. J5 DNA Assembly Design Automation Software. ACS Synth. Biol. 2012, 1 (1), 14–21. https://doi.org/10.1021/sb2000116.

148. Gietz, R. D.; Schiestl, R. H. Frozen Competent Yeast Cells That Can Be Transformed with High Efficiency Using the LiAc/SS Carrier DNA/PEG Method. Nat. Protoc. 2007, 2 (1), 1–4. https://doi.org/10.1038/nprot.2007.17.

149. Wright, R. C. FlowTime: Annotation and Analysis of Biological Dynamical Systems Using Flow Cytometry, 2017. http://bioconductor.org/packages/flowTime/ (accessed 2017-07-27).

150. Zhong, Z.; Ravikumar, A.; Liu, C. C. Tunable Expression Systems for Orthogonal DNA Replication. ACS Synth. Biol. 2018. https://doi.org/10.1021/acssynbio.8b00400.

