## Supplementary File Data Analysis S2 for "Genetically encoded, noise-tolerant, auxin biosensors in yeast facilitate metabolic engineering and directed evolution"

Supplmental reproducible analysis for: Genetically encoded, noise-tolerant, auxin biosensors in yeast facilitate metabolic engineering and directed evolution

Patarasuda Chaisupa^[1]^

Mahbubur Rahman*^[2]^

Sherry B. Hildreth^[3]^

Saede Moseley^[4]^

Chauncey Gatling^[5]^

Richard F. Helm^[6]^

R. Clay Wright^[7]^

```
knitr::opts_chunk$set(echo = TRUE, collapse = TRUE,
                      tidy = TRUE, message = FALSE,
                      warning = FALSE)
library(flowCore)
library(flowTime)
library(ggplot2)
library(stringr)
library(ggridges)
library(dplyr)
library(tidyr)
library(tidyverse)
library(drc)
library(gridExtra)
library(openCyto)
library(ggcyto)
library(flowStats)
library(flowClust)
library(wesanderson)
library(patchwork)
library(ggthemes)
library(agricolae)
```

### Time-course response and ratiometric measurement of the single-fusion biosensors

#### Time-course degradation

```
# Read in flow sets from 20200611 and 20200614
flowSet <- read.plateSet(path = "~/Google Drive/Shared drives/PlantSynBioLab/Data/Mahbub/FlowSets/",
    pattern = "202006", phenoData = "annotation.txt")
flowSet <- flowSet[which(flowSet@phenoData@data$strain %in% c("T1T1", "A2A2"))]

write.flowSet(flowSet, "flowSets/single-time-course")
flowSet <- read.flowSet(path = "flowSets/single-time-course", phenoData = "annotation.txt")
# load gates for this strain/cytometer
load("PSB_Accuri_W303.RData")
data_sum <- summarizeFlow(flowset = flowSet, ploidy = "diploid", only = "singlets")
## [1] "Gating with diploid singlet gates..."
time0_14 <- data_sum %>%
    dplyr::filter(name == "21D10.fcs") %>%
    pull(btime)
time0_11 <- data_sum %>%
    dplyr::filter(name == "11G02.fcs") %>%
    pull(btime)
data_sum <- data_sum %>%
    mutate(time = case_when(folder == "20200611_AFB_epistasis" ~ .$btime - time0_11,
        folder == "20200614_AFB_epistasis" ~ .$btime - time0_14))
shapes <- c(DMSO = 1, IAA = 16)
lines <- c(DMSO = 2, IAA = 1)

data_sum$ratio <- data_sum$FL1.Amean/data_sum$FL4.Amean
data_sum$Venus <- data_sum$FL1.Amean/10000
data_sum_long <- data_sum %>%
    dplyr::select(time, treatment, yWL, folder, strain, ratio, Venus) %>%
    pivot_longer(cols = c(ratio, Venus), names_to = "parameter")
deg_plot <- ggplot(data = subset(data_sum_long, yWL == "166"), aes(x = time, y = value,
    shape = treatment, color = treatment, group = interaction(treatment, folder))) +
    geom_point() + labs(y = "intensity (AU)", x = "time post auxin addition (min)") +
    facet_wrap(~fct_rev(parameter), scales = "free") + scale_shape_manual(values = shapes) +
    geom_line(aes(linetype = treatment)) + scale_linetype_manual(values = lines) +
    scale_color_viridis_d(option = "D", end = 0.75, direction = -1) + theme_test()
deg_plot
```

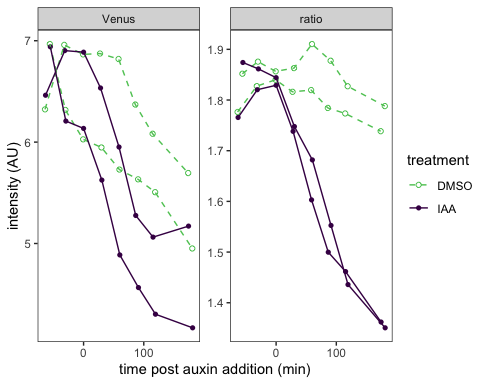

```
data <- flowTime::tidyFlow(flowSet, ploidy = "diploid", only = "singlets")
## [1] "No further gating applied."
## [1] "Converting events..."
# get late time points for one strain
last_reading <- tail(unique(data[which(data$yWL == 166), "name"]), 4) %>%
    head(2)
data <- dplyr::filter(data, yWL == "166" & name %in% last_reading)
# clean this up, cut off zeros
data <- subset(data, FL1.A > 1 & FL4.A > 1)
data$FLratio <- data$FL1.A/data$FL4.A
range(data$FL1.A)
## [1]     21 697262
# calculate cvs
sd(data$FLratio)/mean(data$FLratio)
## [1] 0.1842678
range(data$FLratio)
## [1] 0.002628614 3.836085188
```

#### Single-fusion CV plot

```
# calculate normalized values
data$Venus <- data$FL1.A/median(data$FL1.A)
data$mScarlet <- data$FL4.A/median(data$FL4.A)
data$ratio <- data$FLratio/median(data$FLratio)
# make a tidy, long dataset
data_long <- data %>%
    dplyr::select(treatment, Venus, mScarlet, ratio) %>%
    pivot_longer(cols = c(Venus, mScarlet, ratio), names_to = "parameter", values_to = "value")

# need to also format CVs approriately for annotating
cv <- function(x) return(round(sd(x)/mean(x), 2))
CVs <- data %>%
    group_by(treatment) %>%
    summarise(across(where(is_double), cv))
CVs <- CVs %>%
    dplyr::select(treatment, Venus, mScarlet, ratio) %>%
    pivot_longer(cols = c(Venus, mScarlet, ratio), names_to = "parameter", values_to = "value")

CV_plot <- ggplot(data = data_long, mapping = aes(x = value, color = treatment)) +
    geom_density() + coord_cartesian(x = c(0, 2)) + labs(x = "median normalized intensity",
    color = "treatment") + facet_grid(fct_relevel(parameter, "Venus") ~ .) + theme_test() +
    geom_text(data = subset(CVs, treatment == "IAA"), aes(label = paste0("CV = ",
        value)), x = 0.3, y = 2) + geom_text(data = subset(CVs, treatment == "DMSO"),
    aes(label = paste0("CV = ", value)), x = 1.8, y = 2) + scale_color_viridis_d(option = "D",
    end = 0.75, direction = -1)
CV_plot
```

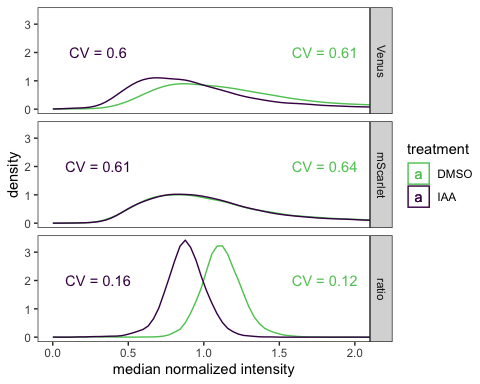

```
layout <- "
AAAAB
CCCCC
"
CV_plot + guide_area() + deg_plot + plot_annotation(tag_levels = "A") + plot_layout(guides = "collect",
    heights = c(5, 2), design = layout)
```

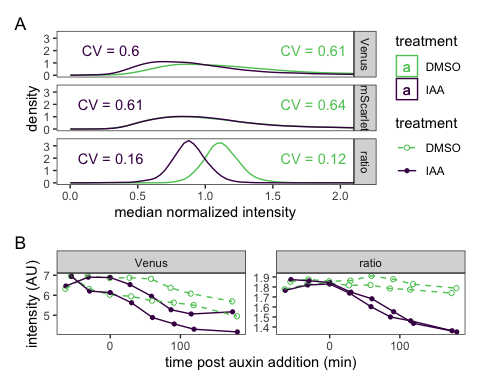

ggsave("ratio-deg.pdf", width = 5, height = 5)
ggsave("ratio-deg.png", width = 5, height = 5)

### Dose-response curves of dual-fusion AFB2 and TIR1 biosensors

#### AFB2-based biosensor (yWL210 AFB2 dual-fusion, single ratiometric construct)

```
plate_all_210 <- read.plateSet(path = "~/Google Drive/Shared drives/PlantSynBioLab/Pat/Experiments/Does-response assay/11212022_DRA_overlaydata/Combine Data_yWL210/All data/",
    pattern = "DRA-*")

annotation <- createAnnotation(yourFlowSet = plate_all_210)
write.csv(annotation, "/Users/patchaisupa/Google Drive/Shared drives/PlantSynBioLab/Pat/Experiments/Does-response assay/11212022_DRA_overlaydata/overlaydata_annotation_yWL210_datagated_exJan31.csv")
annotation <- read.csv("~/Google Drive/Shared drives/PlantSynBioLab/Pat/Experiments/Does-response assay/11212022_DRA_overlaydata/overlaydata_annotation_yWL210_datagated_exJan31.csv")

aplate_all_210 <- annotateFlowSet(yourFlowSet = plate_all_210, annotation_df = annotation,
    mergeBy = "name")
head(rownames(pData(aplate_all_210)))
head(pData(aplate_all_210))
write.flowSet(aplate_all_210, outdir = "flowSets/AFB2-dual-yWL210-dose-response")
aplate_all_210 <- read.flowSet(path = "flowSets/AFB2-dual-yWL210-dose-response",
    phenoData = "annotation.txt")

plate_all_210_sum <- summarizeFlow(aplate_all_210, channel = NA, gated = TRUE)
## [1] "Summarizing all events..."
```

###Dose-response curve

###### Comparing log-logistic and Weibull models

(Figure 2 in Ritz (2009))

```
fitdrc.m1 <- drm(YL1.Amean/BL1.Amean ~ dose, data = plate_all_210_sum, fct = LL.4())
fitdrc.m2 <- drm(YL1.Amean/BL1.Amean ~ dose, data = plate_all_210_sum, fct = W1.4())
# fitdrc.m3 <- drm(YL1.Amean/BL1.Amean~dose, data=plate_all_210_sum, fct =
# W2.4()) Error in drmOpt(opfct, opdfct1, startVecSc, optMethod, constrained,
# warnVal, : Convergence failed

summary(fitdrc.m1)
##
## Model fitted: Log-logistic (ED50 as parameter) (4 parms)
##
## Parameter estimates:
##
##                 Estimate Std. Error t-value p-value
## b:(Intercept) -1.5032417  0.6338695 -2.3715 0.01985 *
## c:(Intercept)  0.0526074  0.0020790 25.3043 < 2e-16 ***
## d:(Intercept)  0.0845079  0.0021128 39.9976 < 2e-16 ***
## e:(Intercept)  0.0403739  0.0159429  2.5324 0.01306 *
## ---
## Signif. codes:  0 '***' 0.001 '**' 0.01 '*' 0.05 '.' 0.1 ' ' 1
##
## Residual standard error:
##
##  0.01246335 (90 degrees of freedom)
summary(fitdrc.m2)
##
## Model fitted: Weibull (type 1) (4 parms)
##
## Parameter estimates:
##
##                 Estimate Std. Error t-value   p-value
## b:(Intercept) -0.7846961  0.2493021 -3.1476  0.002233 **
## c:(Intercept)  0.0522487  0.0021769 24.0009 < 2.2e-16 ***
## d:(Intercept)  0.0847793  0.0022414 37.8241 < 2.2e-16 ***
## e:(Intercept)  0.0189778  0.0111048  1.7090  0.090902 .
## ---
## Signif. codes:  0 '***' 0.001 '**' 0.01 '*' 0.05 '.' 0.1 ' ' 1
##
## Residual standard error:
##
##  0.01263587 (90 degrees of freedom)
# summary(fitdrc.m3)
```

The 4-parameter log-logistic model has a slightly lower residual standard error.

```
model.LL4_all_210 <- drm(YL1.Amean/BL1.Amean ~ dose, data = plate_all_210_sum, fct = LL.4(names = c("Slope",
    "Lower Limit", "Upper Limit", "ED50")))
plot(model.LL4_all_210, broken = TRUE, type = "none", lty = 1, lwd = 5, xlab = "Extracellular IAA concentration (µM)",
    ylab = "Response Signal")
# plot(model.LL4_all_210, broken = TRUE, col = 'black', add=TRUE)
plot(model.LL4_all_210, broken = TRUE, type = "confidence", col = "black", add = TRUE)
```

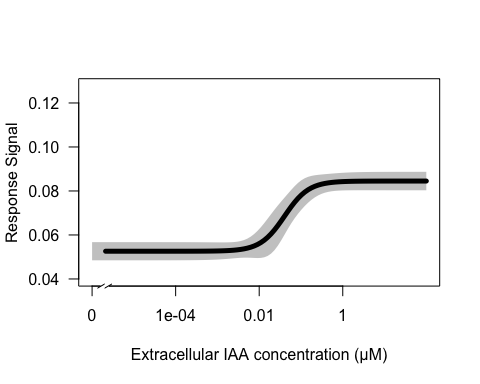

```
summary(model.LL4_all_210)
##
## Model fitted: Log-logistic (ED50 as parameter) (4 parms)
##
## Parameter estimates:
##
##                           Estimate Std. Error t-value p-value
## Slope:(Intercept)       -1.5032417  0.6338695 -2.3715 0.01985 *
## Lower Limit:(Intercept)  0.0526074  0.0020790 25.3043 < 2e-16 ***
## Upper Limit:(Intercept)  0.0845079  0.0021128 39.9976 < 2e-16 ***
## ED50:(Intercept)         0.0403739  0.0159429  2.5324 0.01306 *
## ---
## Signif. codes:  0 '***' 0.001 '**' 0.01 '*' 0.05 '.' 0.1 ' ' 1
##
## Residual standard error:
##
##  0.01246335 (90 degrees of freedom)
```

##### Individual replicate dose-response curves

```
replicate1_210 <- drm(YL1.Amean/BL1.Amean ~ dose, data = subset(plate_all_210_sum,
    replicate == "1"), fct = LL.4())
replicate2_210 <- drm(YL1.Amean/BL1.Amean ~ dose, data = subset(plate_all_210_sum,
    replicate == "2"), fct = LL.4())
replicate3_210 <- drm(YL1.Amean/BL1.Amean ~ dose, data = subset(plate_all_210_sum,
    replicate == "3"), fct = LL.4())
replicate4_210 <- drm(YL1.Amean/BL1.Amean ~ dose, data = subset(plate_all_210_sum,
    replicate == "4"), fct = LL.4())
replicate5_210 <- drm(YL1.Amean/BL1.Amean ~ dose, data = subset(plate_all_210_sum,
    replicate == "5"), fct = LL.4())
replicate6_210 <- drm(YL1.Amean/BL1.Amean ~ dose, data = subset(plate_all_210_sum,
    replicate == "6"), fct = LL.4())
replicate7_210 <- drm(YL1.Amean/BL1.Amean ~ dose, data = subset(plate_all_210_sum,
    replicate == "7"), fct = LL.4())

plot(replicate1_210, broken = TRUE, type = "all", col = "dark green", lty = 2)
plot(replicate2_210, broken = TRUE, add = TRUE, type = "all", col = "dark blue",
    lty = 2)
plot(replicate3_210, broken = TRUE, add = TRUE, type = "all", col = "yellow", lty = 2)
plot(replicate4_210, broken = TRUE, add = TRUE, type = "all", col = "dark grey",
    lty = 2)
plot(replicate5_210, broken = TRUE, add = TRUE, type = "all", col = "dark orange",
    lty = 2)
plot(replicate6_210, broken = TRUE, add = TRUE, type = "all", col = "brown", lty = 2)
plot(replicate7_210, broken = TRUE, add = TRUE, type = "all", col = "dark red", lty = 2)
plot(model.LL4_all_210, broken = TRUE, add = TRUE, type = "none", lty = 1, lwd = 5,
    xlab = "Extracellular IAA concentration (µM)", ylab = "Response Signal")
# plot(model.LL4_all_210, broken = TRUE, col = 'black', add=TRUE)
plot(model.LL4_all_210, broken = TRUE, type = "confidence", col = "black", add = TRUE)
```

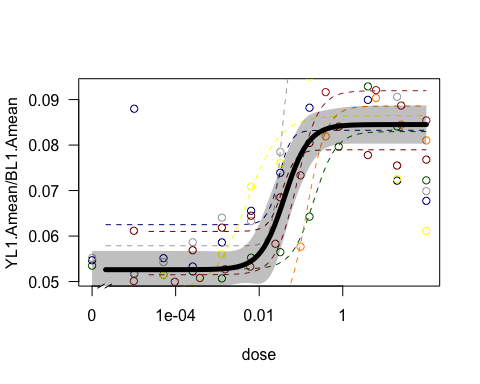

#### TIR1-based biosensor (yWL209 TIR dual-fusion, single ratiometric construct)

```
plate_all_209 <- read.plateSet(path = "~/Google Drive/Shared drives/PlantSynBioLab/Pat/Experiments/Does-response assay/11212022_DRA_overlaydata/Combine Data_yWL209/",
    pattern = "DRA-*")
annotation <- createAnnotation(yourFlowSet = plate_all_209)
write.csv(annotation, "/Users/patchaisupa/Google Drive/Shared drives/PlantSynBioLab/Pat/Experiments/Does-response assay/11212022_DRA_overlaydata/overlaydata_annotation_yWL209.csv")
annotation <- read.csv("~/Google Drive/Shared drives/PlantSynBioLab/Pat/Experiments/Does-response assay/11212022_DRA_overlaydata/overlaydata_annotation_yWL209.csv")

aplate_all_209 <- annotateFlowSet(yourFlowSet = plate_all_209, annotation_df = annotation,
    mergeBy = "name")
head(rownames(pData(aplate_all_209)))
head(pData(aplate_all_209))

write.flowSet(aplate_all_209, outdir = "flowSets/TIR1-dual-yWL209-dose-response")
aplate_all_209 <- read.flowSet(path = "flowSets/TIR1-dual-yWL209-dose-response",
    phenoData = "annotation.txt")

dat_sum_overlaydata_209 <- summarizeFlow(aplate_all_209, gated = TRUE)
## [1] "Summarizing all events..."
```

###Dose-response curve

```
model.LL4_rep1_209 <- drm(YL1.Amean/BL1.Amean ~ dose, data = subset(dat_sum_overlaydata_209,
    replicate == "1"), fct = LL.4(names = c("Slope", "Lower Limit", "Upper Limit",
    "ED50")))
plot(model.LL4_rep1_209, type = "all", col = "red", lty = 1, lwd = 2, xlab = "Extracellular IAA concentration (µM)",
    ylab = "Red/Green Fluorescent Ratio")
plot(model.LL4_rep1_209, broken = TRUE, type = "confidence", col = "red", add = TRUE)
```

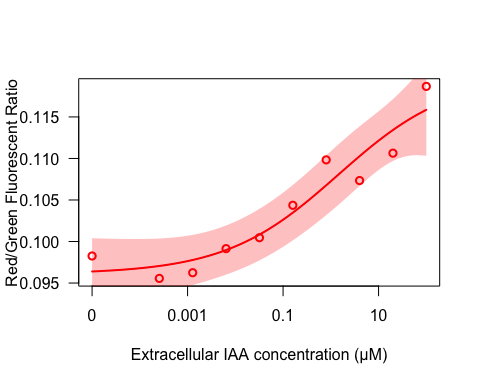

```
print(summary(model.LL4_rep1_209))
##
## Model fitted: Log-logistic (ED50 as parameter) (4 parms)
##
## Parameter estimates:
##
##                           Estimate Std. Error t-value   p-value
## Slope:(Intercept)       -0.3671010  0.1395354 -2.6309   0.03901 *
## Lower Limit:(Intercept)  0.0960915  0.0019149 50.1808 4.201e-09 ***
## Upper Limit:(Intercept)  0.1200258  0.0068403 17.5470 2.198e-06 ***
## ED50:(Intercept)         1.4438822  2.8035445  0.5150   0.62496
## ---
## Signif. codes:  0 '***' 0.001 '**' 0.01 '*' 0.05 '.' 0.1 ' ' 1
##
## Residual standard error:
##
##  0.002705295 (6 degrees of freedom)


model.LL4_rep2_209 <- drm(YL1.Amean/BL1.Amean ~ dose, data = subset(dat_sum_overlaydata_209,
    replicate == "2"), fct = LL.4(names = c("Slope", "Lower Limit", "Upper Limit",
    "ED50")))
plot(model.LL4_rep2_209, type = "all", col = "blue", lty = 1, lwd = 2, xlab = "Extracellular IAA concentration (µM)",
    ylab = "Red/Green Fluorescent Ratio")
plot(model.LL4_rep2_209, broken = TRUE, type = "confidence", col = "blue", add = TRUE)
```

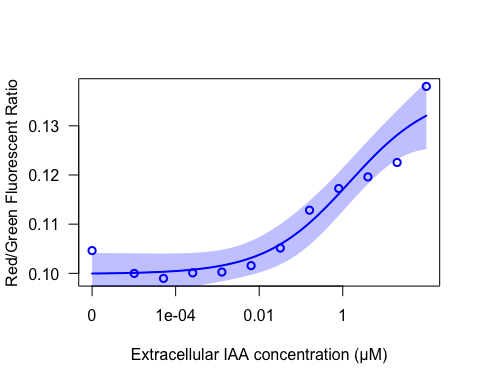

```
print(summary(model.LL4_rep2_209))
##
## Model fitted: Log-logistic (ED50 as parameter) (4 parms)
##
## Parameter estimates:
##
##                           Estimate Std. Error t-value   p-value
## Slope:(Intercept)       -0.4323121  0.1153679 -3.7472  0.005646 **
## Lower Limit:(Intercept)  0.0998826  0.0018914 52.8084 1.833e-11 ***
## Upper Limit:(Intercept)  0.1372100  0.0062312 22.0197 1.910e-08 ***
## ED50:(Intercept)         1.4546642  1.4213107  1.0235  0.336036
## ---
## Signif. codes:  0 '***' 0.001 '**' 0.01 '*' 0.05 '.' 0.1 ' ' 1
##
## Residual standard error:
##
##  0.003755989 (8 degrees of freedom)


model.LL4_rep3_209 <- drm(YL1.Amean/BL1.Amean ~ dose, data = subset(dat_sum_overlaydata_209,
    replicate == "3"), fct = LL.4(names = c("Slope", "Lower Limit", "Upper Limit",
    "ED50")))
plot(model.LL4_rep3_209, type = "all", col = "dark green", lty = 1, lwd = 2, xlab = "Extracellular IAA concentration (µM)",
    ylab = "Red/Green Fluorescent Ratio")
plot(model.LL4_rep3_209, broken = TRUE, type = "confidence", col = "dark green",
    add = TRUE)
```

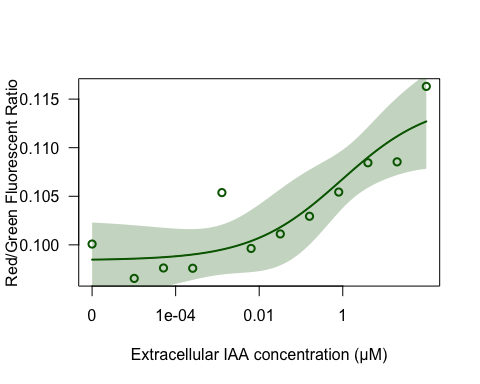

```

print(summary(model.LL4_rep3_209))
##
## Model fitted: Log-logistic (ED50 as parameter) (4 parms)
##
## Parameter estimates:
##
##                           Estimate Std. Error t-value   p-value
## Slope:(Intercept)       -0.4007190  0.2177393 -1.8404    0.1030
## Lower Limit:(Intercept)  0.0984103  0.0018186 54.1128 1.509e-11 ***
## Upper Limit:(Intercept)  0.1148849  0.0037802 30.3916 1.492e-09 ***
## ED50:(Intercept)         0.9199369  1.1883595  0.7741    0.4611
## ---
## Signif. codes:  0 '***' 0.001 '**' 0.01 '*' 0.05 '.' 0.1 ' ' 1
##
## Residual standard error:
##
##  0.002908608 (8 degrees of freedom)


model.LL4_all_209 <- drm(YL1.Amean/BL1.Amean ~ dose, data = dat_sum_overlaydata_209,
    fct = LL.4(names = c("Slope", "Lower Limit", "Upper Limit", "ED50")))
plot(model.LL4_all_209, type = "all", col = "black", lty = 1, lwd = 3)
plot(model.LL4_all_209, broken = TRUE, col = "black", add = TRUE)
plot(model.LL4_all_209, broken = TRUE, type = "confidence", col = "black", add = TRUE)
print(summary(model.LL4_all_209))
##
## Model fitted: Log-logistic (ED50 as parameter) (4 parms)
##
## Parameter estimates:
##
##                           Estimate Std. Error t-value   p-value
## Slope:(Intercept)       -0.3420811  0.0895742 -3.8190 0.0004353 ***
## Lower Limit:(Intercept)  0.0973731  0.0014017 69.4690 < 2.2e-16 ***
## Upper Limit:(Intercept)  0.1337956  0.0109331 12.2377 1.972e-15 ***
## ED50:(Intercept)        10.9140480 20.3972838  0.5351 0.5954208
## ---
## Signif. codes:  0 '***' 0.001 '**' 0.01 '*' 0.05 '.' 0.1 ' ' 1
##
## Residual standard error:
##
##  0.004904408 (42 degrees of freedom)

replicate1_209 <- drm(YL1.Amean/BL1.Amean ~ dose, data = subset(dat_sum_overlaydata_209,
    replicate == "1"), fct = LL.4())
replicate2_209 <- drm(YL1.Amean/BL1.Amean ~ dose, data = subset(dat_sum_overlaydata_209,
    replicate == "2"), fct = LL.4())
replicate3_209 <- drm(YL1.Amean/BL1.Amean ~ dose, data = subset(dat_sum_overlaydata_209,
    replicate == "3"), fct = LL.4())


plot(replicate1_209, broken = TRUE, add = TRUE, type = "none", col = "red", lty = 2)
plot(replicate2_209, broken = TRUE, add = TRUE, type = "none", col = "blue", lty = 2)
plot(replicate3_209, broken = TRUE, add = TRUE, type = "none", col = "dark green",
    lty = 2)
```

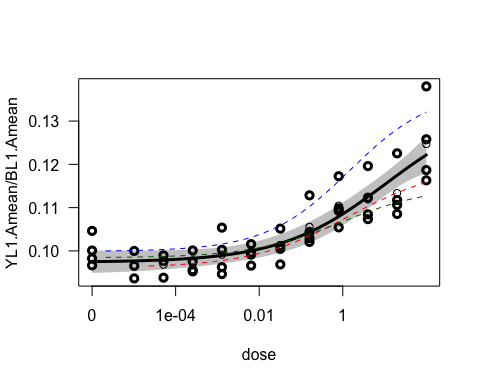

#### Combined Dose-response curveswith ggplot

```
pm210 <- expand.grid(treatment = exp(seq(log(1e-05), log(100), length = 1000)))
pm210 <- cbind(pm210, predict(model.LL4_all_210, newdata = pm210, interval = "confidence"))

plot210 <- ggplot(plate_all_210_sum, aes(x = dose, y = YL1.Amean/BL1.Amean)) + scale_x_log10() +
    geom_ribbon(data = pm210, aes(x = treatment, y = Prediction, ymin = Lower, ymax = Upper),
        alpha = 0.4) + geom_line(data = pm210, aes(x = treatment, y = Prediction),
    linewidth = 1.2) + ylab("mScarlet-I/Venus") + xlab("Auxin (IAA, µM)") + scale_x_log10(labels = scales::label_number(drop0trailing = TRUE)) +
    scale_color_viridis_c() + geom_point(aes(color = replicate), size = 1, alpha = 0.6) +
    theme_classic() + theme(legend.position = "none", plot.title = element_text(hjust = 0.5),
    plot.title.position = "plot") + ggtitle("AFB2")
plot210
```

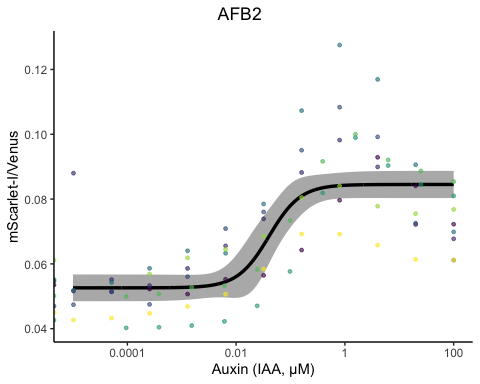

```


pm209 <- expand.grid(treatment = exp(seq(log(1e-05), log(100), length = 1000)))
pm209 <- cbind(pm209, predict(model.LL4_all_209, newdata = pm209, interval = "confidence"))

plot209 <- ggplot(dat_sum_overlaydata_209, aes(x = dose, y = YL1.Amean/BL1.Amean)) +
    scale_x_log10() + geom_ribbon(data = pm209, aes(x = treatment, y = Prediction,
    ymin = Lower, ymax = Upper), alpha = 0.4) + geom_line(data = pm209, aes(x = treatment,
    y = Prediction), linewidth = 1.2) + ylab("mScarlet-I/Venus") + xlab("Auxin (IAA, µM)") +
    scale_x_log10(labels = scales::label_number(drop0trailing = TRUE)) + scale_color_viridis_c() +
    geom_point(aes(color = replicate), size = 1, alpha = 0.6) + theme_classic() +
    theme(legend.position = "none", plot.title = element_text(hjust = 0.5), plot.title.position = "plot") +
    # ylim(0.095, 0.12) +
ggtitle("TIR1")
plot209
```

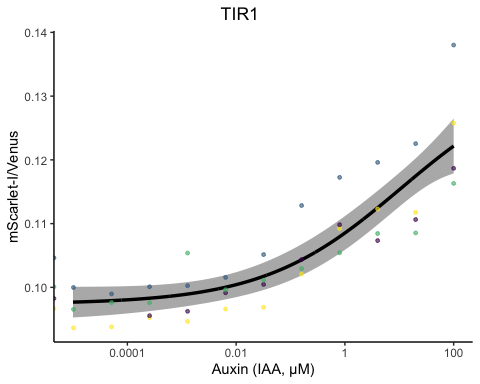

plot210 + plot209 + patchwork::plot_annotation(tag_levels = "A")

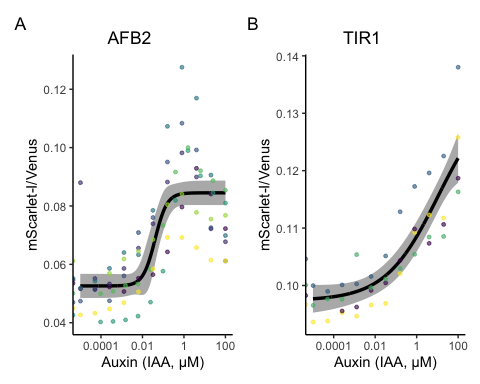

```

ggsave(plot = plot210 + plot209 + patchwork::plot_annotation(tag_levels = "A"), filename = "dose-response.pdf",
    width = 6, height = 3)
ggsave(plot = plot210 + plot209 + patchwork::plot_annotation(tag_levels = "A"), filename = "dose-response.png",
    width = 6, height = 3)
```

### LCMS measurements of intracellular auxin

```
plate_08052022 <- read.plateSet(path = "~/Google Drive/Shared drives/PlantSynBioLab/Pat/Experiments/Does-response assay/08052022_yWL210_DRA-LCMS-R2/Alldata/",
    pattern = "DRA*")
annotation <- read.csv("~/Google Drive/Shared drives/PlantSynBioLab/Pat/Experiments/Does-response assay/08052022_yWL210_DRA-LCMS-R2/08052022_alldata_annotation.csv")

aplate_08052022 <- annotateFlowSet(yourFlowSet = plate_08052022, annotation_df = annotation,
    mergeBy = "name")
head(rownames(pData(aplate_08052022)))
head(pData(aplate_08052022))
write.flowSet(aplate_08052022, "flowSets/AFB2-dual-DR-LCMS")
aplate_08052022 <- read.flowSet(path = "flowSets/AFB2-dual-DR-LCMS", phenoData = "annotation.txt")
plate_08052022_sum <- summarizeFlow(aplate_08052022, gated = TRUE)
## [1] "Summarizing all events..."
model.LL4_08052022 <- drm(YL1.Amean/BL1.Amean ~ dose, data = subset(plate_08052022_sum,
    reading == "10"), fct = LL.4(names = c("Slope", "Lower Limit", "Upper Limit",
    "ED50")))
summary(model.LL4_08052022)
##
## Model fitted: Log-logistic (ED50 as parameter) (4 parms)
##
## Parameter estimates:
##
##                           Estimate Std. Error t-value   p-value
## Slope:(Intercept)       -1.0145173  0.2536243 -4.0001  0.003949 **
## Lower Limit:(Intercept)  0.0485554  0.0019439 24.9779 7.061e-09 ***
## Upper Limit:(Intercept)  0.0735356  0.0010291 71.4543 1.639e-12 ***
## ED50:(Intercept)         0.0987332  0.0305612  3.2307  0.012045 *
## ---
## Signif. codes:  0 '***' 0.001 '**' 0.01 '*' 0.05 '.' 0.1 ' ' 1
##
## Residual standard error:
##
##  0.002267079 (8 degrees of freedom)
pm_08052022 <- expand.grid(treatment = exp(seq(log(0.001), log(100), length = 1000)))
pm_08052022 <- cbind(pm_08052022, predict(model.LL4_08052022, newdata = pm_08052022,
    interval = "confidence"))

plot_08052022 <- ggplot(data = subset(plate_08052022_sum, reading == "10"), aes(x = dose,
    y = YL1.Amean/BL1.Amean)) + geom_ribbon(data = pm_08052022, aes(x = treatment,
    y = Prediction, ymin = Lower, ymax = Upper), alpha = 0.4) + geom_line(data = pm_08052022,
    aes(x = treatment, y = Prediction)) + xlab("Extracellular IAA (µM)") + ylab("AFB2-mScarlet-/Venus-IAA17") +
    scale_x_log10(labels = scales::label_number(drop0trailing = TRUE)) + geom_point(color = "black") +
    theme_classic(base_size = 12) + theme(legend.position = "none")
plot_08052022
```

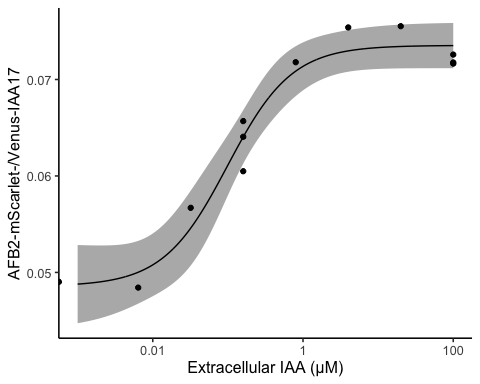

```

LCMSdata <- read.csv("08052022_yWL210_DRA-LCMS_data.csv")

model.LL4_08052022_LCMS <- drm(Observed_IAA_in_uM ~ Dose_IAA, data = LCMSdata, fct = LL.4(names = c("Slope",
    "Lower Limit", "Upper Limit", "ED50")))
summary(model.LL4_08052022_LCMS)
##
## Model fitted: Log-logistic (ED50 as parameter) (4 parms)
##
## Parameter estimates:
##
##                            Estimate  Std. Error t-value   p-value
## Slope:(Intercept)       -5.6514e-01  8.9868e-02 -6.2885 4.016e-05 ***
## Lower Limit:(Intercept)  3.7324e-02  3.6766e-01  0.1015    0.9208
## Upper Limit:(Intercept)  2.6037e+02  2.7474e+02  0.9477    0.3620
## ED50:(Intercept)         1.1866e+05  2.1885e+05  0.5422    0.5976
## ---
## Signif. codes:  0 '***' 0.001 '**' 0.01 '*' 0.05 '.' 0.1 ' ' 1
##
## Residual standard error:
##
##  1.019652 (12 degrees of freedom)

plot(model.LL4_08052022_LCMS, type = "all", col = "black", lty = 1, lwd = 3, xlab = "Extracellular concentrations of IAA (uM)",
    ylab = "Observed IAA (uM)")
```

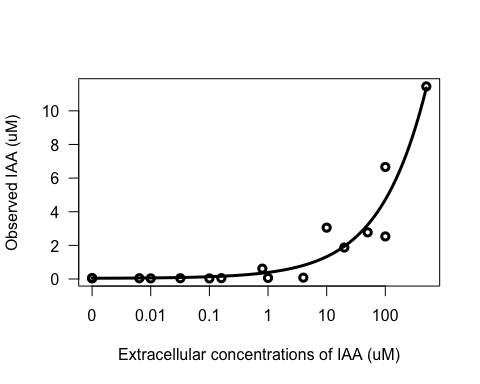

```

pm_08052022_LCMS <- expand.grid(treatment = exp(seq(log(0.001), log(500), length = 1000)))
pm_08052022_LCMS <- cbind(pm_08052022_LCMS, predict(model.LL4_08052022_LCMS, newdata = pm_08052022_LCMS,
    interval = "confidence"))
LCMS_intracellular <- ggplot(data = LCMSdata, mapping = aes(x = Dose_IAA, y = Observed_IAA_in_uM)) +
    geom_point() + geom_ribbon(data = pm_08052022_LCMS, aes(x = treatment, y = Prediction,
    ymin = Lower, ymax = Upper), alpha = 0.4) + geom_line(data = pm_08052022_LCMS,
    aes(x = treatment, y = Prediction)) + scale_x_log10(labels = scales::label_number(drop0trailing = TRUE)) +
    theme_classic(base_size = 13) + theme(legend.position = "none") + ylab("Intracellular IAA (µM)") +
    xlab("Extracellular IAA (µM)")

plot_08052022 + LCMS_intracellular + plot_annotation(tag_levels = "A")
```

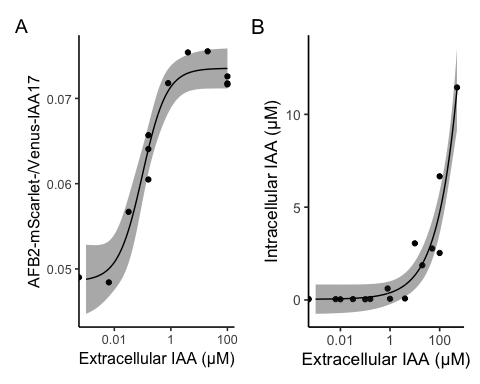

ggsave("LCMS-comparison.pdf", width = 6, height = 3)
ggsave("LCMS-comparison.png", width = 6, height = 3)

### Auxin-induced degradation time-course assay for the dual-fusion biosensors in different yeast strains

Two isolated colonies from each strain were selected at random and tested in auxin-induced protein degradation assay. IAA working solution was added to obtain the IAA concentration at 50 µM in each culture. The assay was carried out using ThermoFisher Attune NxT B/Y flow cytometer.

Strains:

- yWL185 (TIR1 dual-fusion in W303)
- yWL186 (AFB2 dual-fusion in W303)
- yWL209 (TIR1 dual-fusion in YPH499)
- yWL210 (AFB2 dual-fusion in YPH499)

```
plate_03112022 <- read.plateSet(path = "~/Google Drive/Shared drives/PlantSynBioLab/Pat/Experiments/Time course assays/03112022_Time-course assay/10readings/onlyPatstrains/",
    pattern = "TCA*")
annotation <- createAnnotation(yourFlowSet = plate_03112022)
write.csv(annotation, "~/Google Drive/Shared drives/PlantSynBioLab/Pat/Experiments/Time course assays/03112022_Time-course assay/03112022_Time-course assay_10platesPatStrains2.csv")
annotation <- read.csv("~/Google Drive/Shared drives/PlantSynBioLab/Pat/Experiments/Time course assays/03112022_Time-course assay/03112022_Time-course assay_10platesPatStrains2.csv")
aplate_03112022 <- annotateFlowSet(yourFlowSet = plate_03112022, annotation_df = annotation,
    mergeBy = "name")
head(rownames(pData(aplate_03112022)))
head(pData(aplate_03112022))

write.flowSet(aplate_03112022, outdir = "flowSets/dual-time-course")
aplate_03112022 <- read.flowSet(path = "flowSets/dual-time-course/", phenoData = "annotation.txt")
plate_03112022_sum <- summarizeFlow(aplate_03112022, gated = TRUE)
## [1] "Summarizing all events..."


plate_03112022_sum <- plate_03112022_sum %>%
    mutate(background_p = case_when(strain %in% c("yWL185", "yWL186") ~ "W303", strain %in%
        c("yWL209", "yWL210") ~ "YPH499"), receptor_p = case_when(strain %in% c("yWL185",
        "yWL209") ~ "TIR1", strain %in% c("yWL186", "yWL210") ~ "AFB2"))
# The time auxin addition is equal to time zero
time0 <- "303112022-Pat-TCA03_Time-course assay_Auxin_yWL185-C1.fcs"
# or whatever well was being read when auxin was added

plate_03112022_sum$time <- plate_03112022_sum$btime - plate_03112022_sum[[which(plate_03112022_sum$name ==
    time0), "btime"]]
# single bracket -->extracting the all name, 2 brackets extract just single
# 'value' or 'values'
# To normalize data

plate_03112022_sum <- plate_03112022_sum %>%
    mutate(ratio = BL1.Amean/YL1.Amean) %>%
    group_by(background_p, receptor_p) %>%
    mutate(normalizedratio = ratio/mean(ratio))
ratio <- ggplot(data = subset(plate_03112022_sum), aes(x = time, y = normalizedratio,
    group = interaction(factor(colony), factor(treatment)), linetype = factor(treatment),
    shape = factor(treatment), color = factor(treatment))) + geom_point(aes(color = treatment),
    size = 2) + geom_line(aes(color = treatment), linewidth = 1) + xlab("Time post auxin addition (min)") +
    scale_shape_manual(values = c(19, 1)) + ylab("Normalized Venus/mScarlet-I") +
    facet_grid(background_p ~ receptor_p) + scale_color_manual(values = c("#5499C7",
    "#999999")) + theme_base() + theme(legend.position = "none", panel.grid.minor = element_line(linewidth = 0.3,
    linetype = "solid", colour = "white"), axis.line = element_line(colour = "black",
    linewidth = 1, linetype = "solid"))
ratio
```

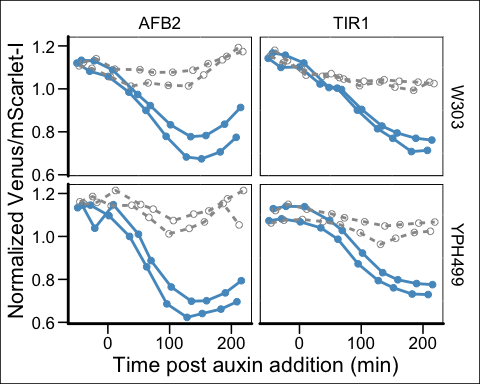

Without normalization

```
ratio_raw <- ggplot(data = subset(plate_03112022_sum), aes(x = time, y = ratio, group = interaction(factor(colony),
    factor(treatment)), linetype = factor(treatment), shape = factor(treatment),
    color = factor(treatment))) + geom_point(aes(color = treatment), size = 2) +
    geom_line(aes(color = treatment), linewidth = 1) + xlab("Time post auxin addition (min)") +
    scale_shape_manual(values = c(19, 1)) + ylab("Venus/mScarlet-I") + facet_grid(background_p ~
    receptor_p, scales = "free") + scale_color_manual(values = c("#5499C7", "#999999")) +
    theme_base() + theme(legend.position = "none", panel.grid.minor = element_line(linewidth = 0.3,
    linetype = "solid", colour = "white"), axis.line = element_line(colour = "black",
    linewidth = 1, linetype = "solid"))
ratio_raw
```

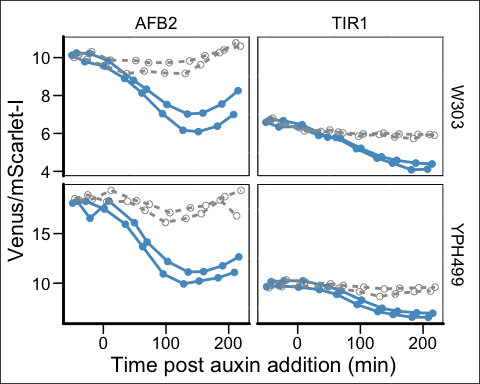

ratio_raw/ratio + plot_annotation(tag_levels = "A") & theme(plot.background = element_blank())

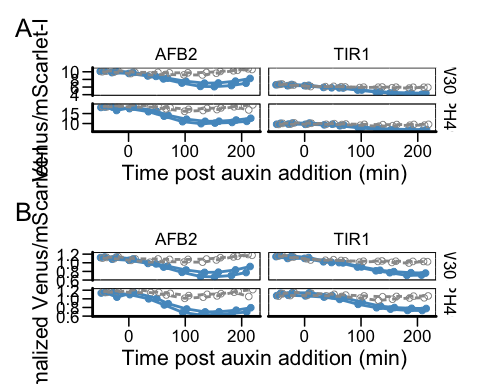

```
ggsave("tc-strains.pdf", width = 5, height = 7)
ggsave("tc-strains.png", width = 5, height = 7)
# normalize green and red fluorescent expression
plate_03112022_sum <- plate_03112022_sum %>%
    mutate(green = BL1.Amean) %>%
    group_by(background_p, receptor_p) %>%
    mutate(normalized_Greenexpression = BL1.Amean/mean(BL1.Amean)) %>%
    mutate(red = YL1.Amean) %>%
    mutate(normalized_Redexpression = YL1.Amean/mean(YL1.Amean))

normgreen <- qplot(x = time, y = normalized_Greenexpression, data = plate_03112022_sum,
    group = interaction(factor(colony), factor(treatment)), linetype = factor(treatment),
    shape = factor(treatment), color = factor(treatment)) + geom_point(aes(color = treatment),
    size = 2) + geom_line(aes(color = treatment), linewidth = 1) + xlab("Time post first reading (min)") +
    scale_shape_manual(values = c(19, 1)) + ylab("Normalized Venus expression") +
    facet_grid(receptor_p ~ background_p) + scale_color_manual(values = c("#F1C40F",
    "#85929E")) + theme_minimal() + theme(legend.position = "none", panel.grid.major = element_line(linewidth = 0.3,
    linetype = "solid", colour = "white"), axis.line = element_line(colour = "black",
    linewidth = 1, linetype = "solid"))

normgreen
```

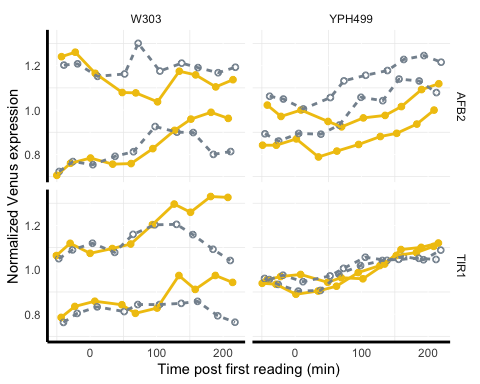

```

normred <- qplot(x = time, y = normalized_Redexpression, data = plate_03112022_sum,
    group = interaction(factor(colony), factor(treatment)), linetype = factor(treatment),
    shape = factor(treatment), color = factor(treatment)) + geom_point(aes(color = treatment),
    size = 2) + geom_line(aes(color = treatment), linewidth = 1) + xlab("Time post first reading (min)") +
    scale_shape_manual(values = c(19, 1)) + ylab("Normalized mScarlet expression") +
    facet_grid(receptor_p ~ background_p) + scale_color_manual(values = c("#EC7063",
    "#616A6B")) + theme_minimal() + theme(legend.position = "none", panel.grid.major = element_line(linewidth = 0.3,
    linetype = "solid", colour = "white"), axis.line = element_line(colour = "black",
    linewidth = 1, linetype = "solid"))
normred
```

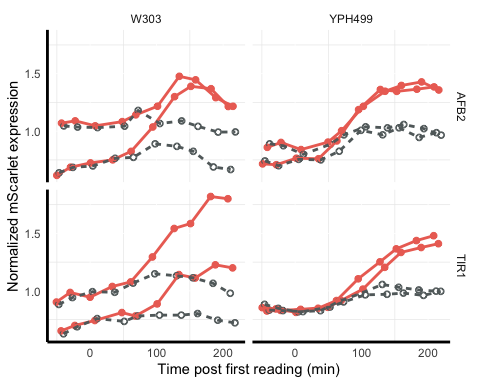

```

green <- qplot(x = time, y = BL1.Amean, data = plate_03112022_sum, group = interaction(factor(colony),
    factor(treatment)), linetype = factor(treatment), shape = factor(treatment),
    color = factor(treatment)) + geom_point(aes(color = treatment), size = 2) + geom_line(aes(color = treatment),
    linewidth = 1) + xlab("Time post first reading (min)") + scale_shape_manual(values = c(19,
    1)) + ylab("Normalized Venus expression") + facet_grid(receptor_p ~ background_p) +
    scale_color_manual(values = c("#F1C40F", "#85929E")) + theme_minimal() + theme(legend.position = "none",
    panel.grid.major = element_line(linewidth = 0.3, linetype = "solid", colour = "white"),
    axis.line = element_line(colour = "black", linewidth = 1, linetype = "solid"))
green
```

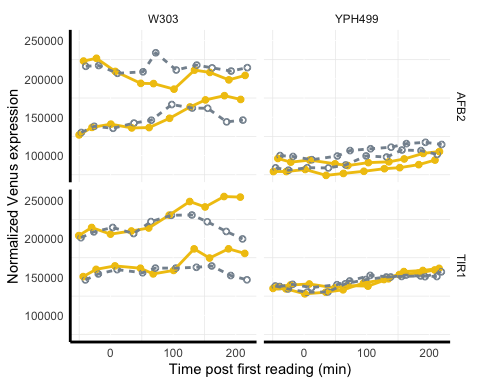

```

red <- qplot(x = time, y = YL1.Amean, data = plate_03112022_sum, group = interaction(factor(colony),
    factor(treatment)), linetype = factor(treatment), shape = factor(treatment),
    color = factor(treatment)) + geom_point(aes(color = treatment), size = 2) + geom_line(aes(color = treatment),
    linewidth = 1) + xlab("Time post first reading (min)") + scale_shape_manual(values = c(19,
    1)) + ylab("Normalized mScarlet expression") + facet_grid(receptor_p ~ background_p) +
    scale_color_manual(values = c("#EC7063", "#616A6B")) + theme_minimal() + theme(legend.position = "none",
    panel.grid.major = element_line(linewidth = 0.3, linetype = "solid", colour = "white"),
    axis.line = element_line(colour = "black", linewidth = 1, linetype = "solid"))
red
```

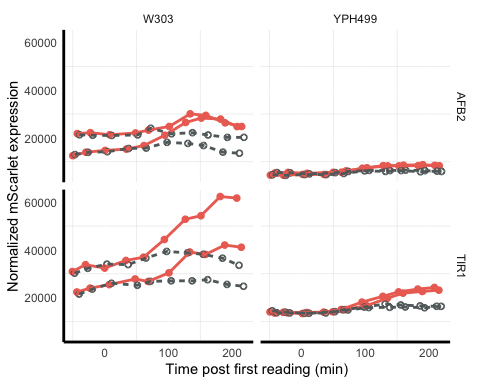

```

conc <- qplot(x = time, y = conc, data = plate_03112022_sum, group = interaction(factor(colony),
    factor(treatment)), linetype = factor(treatment), shape = factor(treatment),
    color = factor(treatment)) + geom_point(aes(color = treatment), size = 2) + geom_line(aes(color = treatment),
    linewidth = 1) + xlab("Time post first reading (min)") + scale_shape_manual(values = c(19,
    1)) + ylab("Normalized mScarlet expression") + facet_grid(receptor_p ~ background_p) +
    scale_color_manual(values = c("black", "#616A6B")) + theme_minimal() + theme(legend.position = "none",
    panel.grid.major = element_line(linewidth = 0.3, linetype = "solid", colour = "white"),
    axis.line = element_line(colour = "black", linewidth = 1, linetype = "solid"))
conc
```

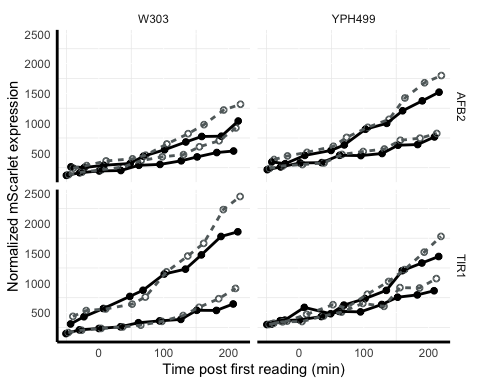

### CV plot

Using the above time course dataset we can compare coefficients of variation in the individual parameters and the ratio for each of the strains and biosensors at steady state.

#### yWL185 (TIR1 dual-fusion, W303 yeast)

```

data185 <- steadyState(aplate_03112022, gated = TRUE)
## [1] "No further gating applied."
## [1] "Converting events..."
data185 <- subset(data185, strain == "yWL185" & name %in% c("803112022-Pat-TCA08_Time-course assay_Auxin_yWL185-C1.fcs",
    "803112022-Pat-TCA08_Time-course assay_Control_yWL185-C1.fcs"))

cv <- function(x) return(round(sd(x)/mean(x), 2))


data185 <- subset(data185, BL1.A > 1 & YL1.A > 1)
data185$FLratio <- data185$BL1.A/data185$YL1.A
range(data185$BL1.A)
## [1]      45 1048575
cv <- function(x) return(round(sd(x)/mean(x), 2))

data185$Venus <- data185$BL1.A/median(data185$BL1.A)
data185$mScarlet <- data185$YL1.A/median(data185$YL1.A)
data185$FLratio <- data185$BL1.A/data185$YL1.A
data185$ratio <- data185$FLratio/median(data185$FLratio)


data_long185 <- data185 %>%
    dplyr::select(treatment, Venus, mScarlet, ratio, strain) %>%
    pivot_longer(cols = c(Venus, mScarlet, ratio), names_to = "parameter", values_to = "value") %>%
    dplyr::mutate(parameter = fct_relevel(parameter, "Venus"))

# need to also format CVs approriately for annotating

CV185 <- data185 %>%
    group_by(treatment) %>%
    dplyr::summarise(across(dplyr::where(is_double), cv)) %>%
    dplyr::select(treatment, Venus, mScarlet, ratio) %>%
    pivot_longer(cols = c(Venus, mScarlet, ratio), names_to = "parameter", values_to = "value")

# data185

plot185 <- ggplot(data = data_long185, mapping = aes(x = value, color = treatment)) +
    geom_density() + xlim(c(-1, 4)) + labs(x = "median normalized intensity", color = "treatment") +
    theme_test() + geom_text(data = subset(CV185, treatment == "50 uM Auxin"), aes(label = paste0("CV = ",
    value)), x = 2, y = 1) + geom_text(data = subset(CV185, treatment == "Control"),
    aes(label = paste0("CV = ", value)), x = 0, y = 1) + scale_color_viridis_d(option = "D",
    end = 0.75, direction = -1)


venus185 <- ggplot(data185, aes(x = data185$Venus, group = treatment, fill = treatment,
    color = treatment)) + geom_density(adjust = 1.5, alpha = 0.5) + labs(x = "Venus",
    y = "Density") + xlim(-0.1, 2) + ylim(0, 2) + theme_classic() + theme(legend.position = "none") +
    geom_text(data = subset(CV185, parameter == "Venus" & treatment == "50 uM Auxin"),
        aes(label = paste0("CV = ", value)), x = 1.2, y = 1.1) + geom_text(data = subset(CV185,
    parameter == "Venus" & treatment == "Control"), aes(label = paste0("CV = ", value)),
    x = 0.55, y = 1.5) + scale_color_manual(values = c("#F1C40F", "#626567")) + scale_fill_manual(values = c("#F1C40F",
    "#626567"))
# venus185

red185 <- ggplot(data185, aes(x = data185$mScarlet, group = treatment, fill = treatment,
    color = treatment)) + geom_density(adjust = 1.5, alpha = 0.5) + labs(x = "mScarlet",
    y = "Density") + xlim(-0.1, 2) + ylim(0, 2) + theme_classic() + theme(legend.position = "none") +
    geom_text(data = subset(CV185, parameter == "mScarlet" & treatment == "50 uM Auxin"),
        aes(label = paste0("CV = ", value)), x = 1.2, y = 1.1) + geom_text(data = subset(CV185,
    parameter == "mScarlet" & treatment == "Control"), aes(label = paste0("CV = ",
    value)), x = 0.55, y = 1.5) + scale_color_manual(values = c("#CB4335", "#626567")) +
    scale_fill_manual(values = c("#CB4335", "#626567"))
# red185


ratio185 <- ggplot(data185, aes(x = data185$ratio, group = treatment, fill = treatment,
    color = treatment)) + geom_density(adjust = 1.5, alpha = 0.5) + labs(x = "Venus/mScarlet ratio",
    y = "Density") + xlim(-0.1, 2) + ylim(0, 3.5) + theme_classic() + theme(legend.position = "none") +
    geom_text(data = subset(CV185, parameter == "ratio" & treatment == "50 uM Auxin"),
        aes(label = paste0("CV = ", value)), x = 0.4, y = 2) + geom_text(data = subset(CV185,
    parameter == "ratio" & treatment == "Control"), aes(label = paste0("CV = ", value)),
    x = 1.6, y = 2) + scale_color_manual(values = c("#5499C7", "#626567")) + scale_fill_manual(values = c("#5499C7",
    "#626567"))

# ratio185

plot185 <- grid.arrange(venus185, red185, ratio185, nrow = 3, ncol = 1)
```

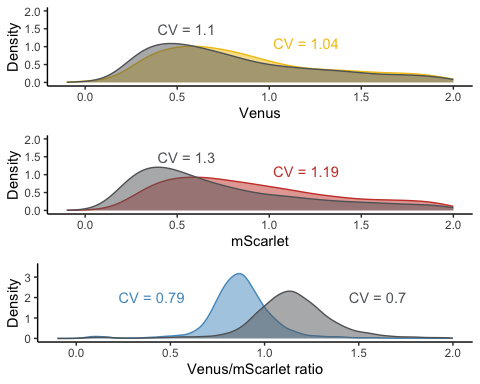

```
plot185
## TableGrob (3 x 1) "arrange": 3 grobs
##   z     cells    name           grob
## 1 1 (1-1,1-1) arrange gtable[layout]
## 2 2 (2-2,1-1) arrange gtable[layout]
## 3 3 (3-3,1-1) arrange gtable[layout]
```

#### yWL186 (AFB2 dual-fusion, W303 yeast)

```
data186 <- steadyState(aplate_03112022, gated = TRUE)
## [1] "No further gating applied."
## [1] "Converting events..."
data186 <- subset(data186, strain == "yWL186" & name %in% c("803112022-Pat-TCA08_Time-course assay_Auxin_yWL186-C1.fcs",
    "803112022-Pat-TCA08_Time-course assay_Control_yWL186-C1.fcs"))

cv <- function(x) return(round(sd(x)/mean(x), 2))


data186 <- subset(data186, BL1.A > 1 & YL1.A > 1)
data186$FLratio <- data186$BL1.A/data186$YL1.A
range(data186$BL1.A)
## [1]      25 1048575
cv <- function(x) return(round(sd(x)/mean(x), 2))

data186$Venus <- data186$BL1.A/median(data186$BL1.A)
data186$mScarlet <- data186$YL1.A/median(data186$YL1.A)
data186$FLratio <- data186$BL1.A/data186$YL1.A
data186$ratio <- data186$FLratio/median(data186$FLratio)


data_long186 <- data186 %>%
    dplyr::select(treatment, Venus, mScarlet, ratio, strain) %>%
    pivot_longer(cols = c(Venus, mScarlet, ratio), names_to = "parameter", values_to = "value") %>%
    dplyr::mutate(parameter = fct_relevel(parameter, "Venus"))

# need to also format CVs approriately for annotating

CV186 <- data186 %>%
    group_by(treatment) %>%
    dplyr::summarise(across(dplyr::where(is_double), cv)) %>%
    dplyr::select(treatment, Venus, mScarlet, ratio) %>%
    pivot_longer(cols = c(Venus, mScarlet, ratio), names_to = "parameter", values_to = "value")

# data186

plot186 <- ggplot(data = data_long186, mapping = aes(x = value, color = treatment)) +
    geom_density() + xlim(c(-1, 4)) + labs(x = "median normalized intensity", color = "treatment") +
    theme_test() + geom_text(data = subset(CV186, treatment == "50 uM Auxin"), aes(label = paste0("CV = ",
    value)), x = 2, y = 1) + geom_text(data = subset(CV186, treatment == "Control"),
    aes(label = paste0("CV = ", value)), x = 0, y = 1) + scale_color_viridis_d(option = "D",
    end = 0.75, direction = -1)


venus186 <- ggplot(data186, aes(x = data186$Venus, group = treatment, fill = treatment,
    color = treatment)) + geom_density(adjust = 1.5, alpha = 0.4) + labs(x = "Venus (normalized median)",
    y = "Density") + xlim(0, 2) + ylim(0, 2) + theme_classic() + theme(legend.position = "none") +
    geom_text(data = subset(CV186, parameter == "Venus" & treatment == "50 uM Auxin"),
        aes(label = paste0("CV = ", value)), x = 1.2, y = 1.1) + geom_text(data = subset(CV186,
    parameter == "Venus" & treatment == "Control"), aes(label = paste0("CV = ", value)),
    x = 0.55, y = 1.5) + scale_color_manual(values = c("#F1C40F", "#626567")) + scale_fill_manual(values = c("#F1C40F",
    "#626567"))


red186 <- ggplot(data186, aes(x = data186$mScarlet, group = treatment, fill = treatment,
    color = treatment)) + geom_density(adjust = 1.5, alpha = 0.4) + labs(x = "mScarlet (normalized median)",
    y = "Density") + xlim(0, 2) + ylim(0, 2) + theme_classic() + theme(legend.position = "none") +
    scale_fill_manual(values = c("#EC7063", "#999999")) + geom_text(data = subset(CV186,
    parameter == "mScarlet" & treatment == "50 uM Auxin"), aes(label = paste0("CV = ",
    value)), x = 1.2, y = 1.1) + geom_text(data = subset(CV186, parameter == "mScarlet" &
    treatment == "Control"), aes(label = paste0("CV = ", value)), x = 0.55, y = 1.5) +
    scale_color_manual(values = c("#CB4335", "#626567")) + scale_fill_manual(values = c("#CB4335",
    "#626567"))


ratio186 <- ggplot(data186, aes(x = data186$ratio, group = treatment, fill = treatment,
    color = treatment)) + geom_density(adjust = 1.5, alpha = 0.4) + labs(x = "Venus/mScarlet ratio",
    y = "Density") + xlim(0, 2) + ylim(0, 3.5) + theme_classic() + theme(legend.position = "none") +
    geom_text(data = subset(CV186, parameter == "ratio" & treatment == "50 uM Auxin"),
        aes(label = paste0("CV = ", value)), x = 0.2, y = 2) + geom_text(data = subset(CV186,
    parameter == "ratio" & treatment == "Control"), aes(label = paste0("CV = ", value)),
    x = 1.3, y = 2) + scale_color_manual(values = c("#5499C7", "#626567")) + scale_fill_manual(values = c("#5499C7",
    "#626567"))

plot186 <- grid.arrange(venus186, red186, ratio186, nrow = 3, ncol = 1)
```

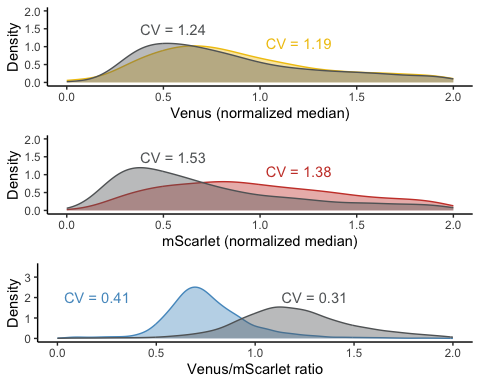

```
plot186
## TableGrob (3 x 1) "arrange": 3 grobs
##   z     cells    name           grob
## 1 1 (1-1,1-1) arrange gtable[layout]
## 2 2 (2-2,1-1) arrange gtable[layout]
## 3 3 (3-3,1-1) arrange gtable[layout]
```

#### yWL209 (TIR1 dual-fusion, YPH499 yeast)

```

data209 <- steadyState(aplate_03112022, gated = TRUE)
## [1] "No further gating applied."
## [1] "Converting events..."
data209 <- subset(data209, strain == "yWL209" & name %in% c("803112022-Pat-TCA08_Time-course assay_Auxin_yWL209-C1.fcs",
    "803112022-Pat-TCA08_Time-course assay_Control_yWL209-C1.fcs"))

cv <- function(x) return(round(sd(x)/mean(x), 2))


data209 <- subset(data209, BL1.A > 1 & YL1.A > 1)
data209$FLratio <- data209$BL1.A/data209$YL1.A
range(data209$BL1.A)
## [1]       9 1048575
cv <- function(x) return(round(sd(x)/mean(x), 2))

data209$Venus <- data209$BL1.A/median(data209$BL1.A)
data209$mScarlet <- data209$YL1.A/median(data209$YL1.A)
data209$FLratio <- data209$BL1.A/data209$YL1.A
data209$ratio <- data209$FLratio/median(data209$FLratio)


data_long209 <- data209 %>%
    dplyr::select(treatment, Venus, mScarlet, ratio, strain) %>%
    pivot_longer(cols = c(Venus, mScarlet, ratio), names_to = "parameter", values_to = "value") %>%
    dplyr::mutate(parameter = fct_relevel(parameter, "Venus"))

# need to also format CVs approriately for annotating

CV209 <- data209 %>%
    group_by(treatment) %>%
    dplyr::summarise(across(dplyr::where(is_double), cv)) %>%
    dplyr::select(treatment, Venus, mScarlet, ratio) %>%
    pivot_longer(cols = c(Venus, mScarlet, ratio), names_to = "parameter", values_to = "value")

# data209

plot209 <- ggplot(data = data_long209, mapping = aes(x = value, color = treatment)) +
    geom_density() + xlim(c(-1, 4)) + labs(x = "median normalized intensity", color = "treatment") +
    theme_test() + geom_text(data = subset(CV209, treatment == "50 uM Auxin"), aes(label = paste0("CV = ",
    value)), x = 1.2, y = 1) + geom_text(data = subset(CV209, treatment == "Control"),
    aes(label = paste0("CV = ", value)), x = 0, y = 1) + scale_color_viridis_d(option = "D",
    end = 0.75, direction = -1)


venus209 <- ggplot(data209, aes(x = data209$Venus, group = treatment, fill = treatment,
    color = treatment)) + geom_density(adjust = 1.5, alpha = 0.4) + labs(x = "Venus",
    y = "Density") + xlim(0, 2) + ylim(0, 2) + theme_classic() + theme(legend.position = "none") +
    geom_text(data = subset(CV209, parameter == "Venus" & treatment == "50 uM Auxin"),
        aes(label = paste0("CV = ", value)), x = 1.2, y = 1.1) + geom_text(data = subset(CV209,
    parameter == "Venus" & treatment == "Control"), aes(label = paste0("CV = ", value)),
    x = 0.55, y = 1.5) + scale_color_manual(values = c("#F1C40F", "#626567")) + scale_fill_manual(values = c("#F1C40F",
    "#626567"))


red209 <- ggplot(data209, aes(x = data209$mScarlet, group = treatment, fill = treatment,
    color = treatment)) + geom_density(adjust = 1.5, alpha = 0.4) + labs(x = "mScarlet",
    y = "Density") + xlim(0, 2) + ylim(0, 2) + theme_classic() + theme(legend.position = "none") +
    geom_text(data = subset(CV209, parameter == "mScarlet" & treatment == "50 uM Auxin"),
        aes(label = paste0("CV = ", value)), x = 1.2, y = 1.1) + geom_text(data = subset(CV209,
    parameter == "mScarlet" & treatment == "Control"), aes(label = paste0("CV = ",
    value)), x = 0.55, y = 1.5) + scale_color_manual(values = c("#CB4335", "#626567")) +
    scale_fill_manual(values = c("#CB4335", "#626567"))


ratio209 <- ggplot(data209, aes(x = data209$ratio, group = treatment, fill = treatment,
    color = treatment)) + geom_density(adjust = 1.5, alpha = 0.4) + labs(x = "Venus/mScarlet ratio",
    y = "Density") + xlim(0, 2) + ylim(0, 3.5) + theme_classic() + theme(legend.position = "none") +
    geom_text(data = subset(CV209, parameter == "ratio" & treatment == "50 uM Auxin"),
        aes(label = paste0("CV = ", value)), x = 0.3, y = 2) + geom_text(data = subset(CV209,
    parameter == "ratio" & treatment == "Control"), aes(label = paste0("CV = ", value)),
    x = 1.6, y = 2) + scale_color_manual(values = c("#5499C7", "#626567")) + scale_fill_manual(values = c("#5499C7",
    "#626567"))


plot209 <- grid.arrange(venus209, red209, ratio209, nrow = 3, ncol = 1)
```

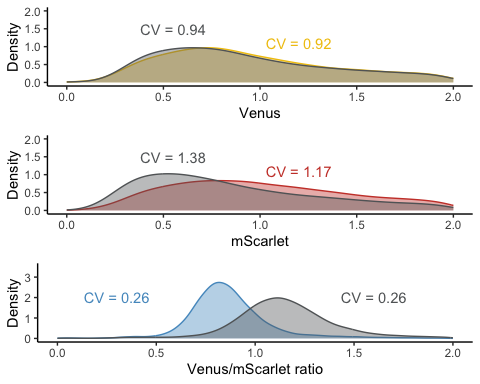

```
plot209
## TableGrob (3 x 1) "arrange": 3 grobs
##   z     cells    name           grob
## 1 1 (1-1,1-1) arrange gtable[layout]
## 2 2 (2-2,1-1) arrange gtable[layout]
## 3 3 (3-3,1-1) arrange gtable[layout]
```

#### yWL210 (AFB2 dual-fusion, YPH499 yeast)

```

data210 <- steadyState(aplate_03112022, gated = TRUE)
## [1] "No further gating applied."
## [1] "Converting events..."
# data <- tidyFlow(aplate_20210619_W303)

data210 <- subset(data210, strain == "yWL210" & name %in% c("803112022-Pat-TCA08_Time-course assay_Auxin_yWL210-C1.fcs",
    "803112022-Pat-TCA08_Time-course assay_Control_yWL210-C1.fcs"))

cv <- function(x) return(round(sd(x)/mean(x), 2))


data210 <- subset(data210, BL1.A > 1 & YL1.A > 1)
data210$FLratio <- data210$BL1.A/data210$YL1.A
range(data210$BL1.A)
## [1]      25 1048575
cv <- function(x) return(round(sd(x)/mean(x), 2))

data210$Venus <- data210$BL1.A/median(data210$BL1.A)
data210$mScarlet <- data210$YL1.A/median(data210$YL1.A)
data210$FLratio <- data210$BL1.A/data210$YL1.A
data210$ratio <- data210$FLratio/median(data210$FLratio)


data_long210 <- data210 %>%
    dplyr::select(treatment, Venus, mScarlet, ratio, strain) %>%
    pivot_longer(cols = c(Venus, mScarlet, ratio), names_to = "parameter", values_to = "value") %>%
    dplyr::mutate(parameter = fct_relevel(parameter, "Venus"))

# need to also format CVs approriately for annotating

CV210 <- data210 %>%
    group_by(treatment) %>%
    dplyr::summarise(across(dplyr::where(is_double), cv)) %>%
    dplyr::select(treatment, Venus, mScarlet, ratio) %>%
    pivot_longer(cols = c(Venus, mScarlet, ratio), names_to = "parameter", values_to = "value")

# data210

plot210 <- ggplot(data = data_long210, mapping = aes(x = value, color = treatment)) +
    geom_density() + xlim(c(-1, 4)) + labs(x = "median normalized intensity", color = "treatment") +
    theme_test() + geom_text(data = subset(CV210, treatment == "50 uM Auxin"), aes(label = paste0("CV = ",
    value)), x = 2, y = 1) + geom_text(data = subset(CV210, treatment == "Control"),
    aes(label = paste0("CV = ", value)), x = 0, y = 1) + scale_color_viridis_d(option = "D",
    end = 0.75, direction = -1) + facet_grid(parameter ~ .)


venus210 <- ggplot(data210, aes(x = data210$Venus, group = treatment, color = treatment,
    fill = treatment)) + geom_density(adjust = 1.5, alpha = 0.4) + labs(x = "Venus",
    y = "Density") + xlim(0, 2) + ylim(0, 2) + theme_classic() + theme(legend.position = "none") +
    geom_text(data = subset(CV210, parameter == "Venus" & treatment == "50 uM Auxin"),
        aes(label = paste0("CV = ", value)), x = 1.5, y = 1) + geom_text(data = subset(CV210,
    parameter == "Venus" & treatment == "Control"), aes(label = paste0("CV = ", value)),
    x = 0.7, y = 1.5) + scale_color_manual(values = c("#F1C40F", "#626567")) + scale_fill_manual(values = c("#F1C40F",
    "#626567"))


red210 <- ggplot(data210, aes(x = data210$mScarlet, group = treatment, color = treatment,
    fill = treatment)) + geom_density(adjust = 1.5, alpha = 0.4) + labs(x = "mScarlet",
    y = "Density") + xlim(0, 2) + ylim(0, 2) + theme_classic() + theme(legend.position = "none") +
    geom_text(data = subset(CV210, parameter == "mScarlet" & treatment == "50 uM Auxin"),
        aes(label = paste0("CV = ", value)), x = 1.2, y = 1.2) + geom_text(data = subset(CV210,
    parameter == "mScarlet" & treatment == "Control"), aes(label = paste0("CV = ",
    value)), x = 0.4, y = 1.5) + scale_color_manual(values = c("#CB4335", "#626567")) +
    scale_fill_manual(values = c("#CB4335", "#626567"))


ratio210 <- ggplot(data210, aes(x = data210$ratio, group = treatment, color = treatment,
    fill = treatment)) + geom_density(adjust = 1.5, alpha = 0.4) + labs(x = "Venus/mScarlet ratio",
    y = "Density") + xlim(0, 2) + ylim(0, 3.5) + theme_classic() + theme(legend.position = "none") +
    geom_text(data = subset(CV210, parameter == "ratio" & treatment == "50 uM Auxin"),
        aes(label = paste0("CV = ", value)), x = 0.2, y = 1.8) + geom_text(data = subset(CV210,
    parameter == "ratio" & treatment == "Control"), aes(label = paste0("CV = ", value)),
    x = 1.6, y = 1.8) + scale_color_manual(values = c("#5499C7", "#626567")) + scale_fill_manual(values = c("#5499C7",
    "#626567"))
# ratio210

plot210 <- grid.arrange(venus210, red210, ratio210, nrow = 3, ncol = 1)
```

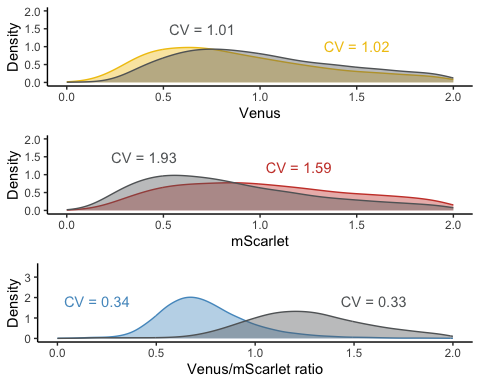

```
plot210
## TableGrob (3 x 1) "arrange": 3 grobs
##   z     cells    name           grob
## 1 1 (1-1,1-1) arrange gtable[layout]
## 2 2 (2-2,1-1) arrange gtable[layout]
## 3 3 (3-3,1-1) arrange gtable[layout]
dual_fusion_cv <- (venus210 + ggtitle("AFB2") + theme(plot.title = element_text(hjust = 0.5)) |
    venus209 + ggtitle("TIR1") + theme(plot.title = element_text(hjust = 0.5)))/(red210 |
    red209)/(ratio210 | ratio209) + theme(title = element_text(hjust = 0))
dual_fusion_cv
```

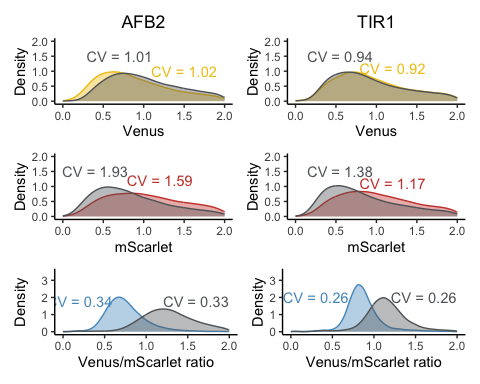

ggsave("dual_fusion_cv.pdf", width = 6.3, height = 6)
ggsave("dual_fusion_cv.png", width = 6.3, height = 6)

### Single-fusion vs dual-fusion comparison in W303 yeast strain

- yWL161 (TIR1 single-fusion, W303 yeast)
- yWL162 (AFB2 single-fusion, W303 yeast)
- yWL185 (TIR1 dual-fusion, W303 yeast)
- yWL186 (AFB2 dual-fusion, W303 yeast)

```
plate_20210619_W303 <- read.plateSet(path = "~/Google Drive/Shared drives/PlantSynBioLab/Pat/Experiments/Time course assays/20210619/W303only/",
    pattern = "r*")
annotation <- createAnnotation(yourFlowSet = plate_20210619_W303)
write.csv(annotation, "/Users/patchaisupa/Google Drive/Shared drives/PlantSynBioLab/Pat/Experiments/Time course assays/20210619/20210619_W303only_annotation.csv")
annotation <- read.csv("~/Google Drive/Shared drives/PlantSynBioLab/Pat/Experiments/Time course assays/20210619/20210619_W303only_annotation.csv")
aplate_20210619_W303 <- annotateFlowSet(yourFlowSet = plate_20210619_W303, annotation_df = annotation,
    mergeBy = "name")
head(rownames(pData(aplate_20210619_W303)))
head(pData(aplate_20210619_W303))
write.flowSet(aplate_20210619_W303, outdir = "flowSets/design-relative-expression")
aplate_20210619_W303 <- read.flowSet(path = "flowSets/design-relative-expression/",
    phenoData = "annotation.txt")
plate_20210619_W303_sum <- summarizeFlow(aplate_20210619_W303, gated = TRUE)
## [1] "Summarizing all events..."


# The time auxin addition is equal to time zero
time0 <- "4E01.fcs"
# or whatever well was being read when auxin was added

plate_20210619_W303_sum$time <- plate_20210619_W303_sum$btime - plate_20210619_W303_sum[[which(plate_20210619_W303_sum$name ==
    time0), "btime"]]
# single bracket -->extracting the all name, 2 brackets extract just single
# 'value' or 'values'
plate_20210619_W303_sum <- plate_20210619_W303_sum %>%
    mutate(design = case_when(strain %in% c("yWL185", "yWL186") ~ "dual-fusion",
        strain %in% c("yWL161", "yWL162") ~ "single-fusion"), fbox = case_when(strain %in%
        c("yWL161", "yWL185") ~ "TIR1", strain %in% c("yWL162", "yWL186") ~ "AFB2")) %>%
    mutate(design_construct = paste(fbox, design))

plate_20210619_W303_sum <- plate_20210619_W303_sum %>%
    mutate(ratio = FL1.Amean/FL4.Amean) %>%
    group_by(design, fbox) %>%
    mutate(normalizedratio = ratio/mean(ratio))

plate_20210619_W303_sum <- plate_20210619_W303_sum %>%
    mutate(green = FL1.Amean) %>%
    group_by(design, fbox) %>%
    mutate(normalized_Greenexpression = FL1.Amean/mean(FL1.Amean))

plate_20210619_W303_sum <- plate_20210619_W303_sum %>%
    mutate(red = FL4.Amean) %>%
    group_by(design, fbox) %>%
    mutate(normalized_Redexpression = FL4.Amean/mean(FL4.Amean))
TIR1_single <- qplot(x = time, y = FL1.Amean/FL4.Amean, data = subset(plate_20210619_W303_sum,
    strain == "yWL161"), group = factor(treatment), linetype = factor(treatment),
    shape = factor(treatment), color = factor(treatment)) + xlab("Time (min)") +
    geom_point(aes(color = treatment, shape = treatment), size = 1) + geom_line(aes(color = treatment,
    shape = treatment), linewidth = 0.5) + scale_shape_manual(values = c(19, 1)) +
    ylab("Venus/Scarlet ratio") + scale_color_manual(values = c("#5499C7", "#626567")) +
    theme_minimal() + theme(legend.position = "none", panel.grid.major = element_line(linewidth = 0.3,
    linetype = "solid", colour = "white"), panel.grid.minor = element_line(linewidth = 0.3,
    linetype = "solid", colour = "white")) + facet_wrap(~design_construct, scale = "free_y")


AFB2_single <- qplot(x = time, y = FL1.Amean/FL4.Amean, data = subset(plate_20210619_W303_sum,
    strain == "yWL162"), group = factor(treatment), linetype = factor(treatment),
    shape = factor(treatment), color = factor(treatment)) + xlab("Time (min)") +
    geom_point(aes(color = treatment, shape = treatment), size = 1) + geom_line(aes(color = treatment,
    shape = treatment), linewidth = 0.5) + scale_shape_manual(values = c(19, 1)) +
    ylab("Venus/Scarlet ratio") + scale_color_manual(values = c("#5499C7", "#626567")) +
    theme_minimal() + theme(legend.position = "none", panel.grid.major = element_line(linewidth = 0.3,
    linetype = "solid", colour = "white"), panel.grid.minor = element_line(linewidth = 0.3,
    linetype = "solid", colour = "white")) + facet_wrap(~design_construct, scale = "free_y")

TIR1_dual <- qplot(x = time, y = FL1.Amean/FL4.Amean, data = subset(plate_20210619_W303_sum,
    strain == "yWL185"), group = factor(treatment), linetype = factor(treatment),
    shape = factor(treatment), color = factor(treatment)) + geom_point(aes(color = treatment,
    shape = treatment), size = 1) + geom_line(aes(color = treatment, shape = treatment),
    linewidth = 0.5) + xlab("Time (min)") + scale_shape_manual(values = c(19, 1)) +
    ylab("Venus/Scarlet ratio") + scale_color_manual(values = c("#5499C7", "#626567")) +
    theme_minimal() + theme(legend.position = "none", panel.grid.major = element_line(linewidth = 0.3,
    linetype = "solid", colour = "white"), panel.grid.minor = element_line(linewidth = 0.3,
    linetype = "solid", colour = "white")) + facet_wrap(~design_construct, scale = "free_y")

AFB2_dual <- qplot(x = time, y = FL1.Amean/FL4.Amean, data = subset(plate_20210619_W303_sum,
    strain == "yWL186"), group = factor(treatment), linetype = factor(treatment),
    shape = factor(treatment), color = factor(treatment)) + xlab("Time (min)") +
    geom_point(aes(color = treatment, shape = treatment), size = 1) + geom_line(aes(color = treatment,
    shape = treatment), linewidth = 0.5) + scale_shape_manual(values = c(19, 1)) +
    ylab("Venus/Scarlet ratio") + scale_color_manual(values = c("#5499C7", "#626567")) +
    theme_minimal() + theme(legend.position = "none", panel.grid.major = element_line(linewidth = 0.3,
    linetype = "solid", colour = "white"), panel.grid.minor = element_line(linewidth = 0.3,
    linetype = "solid", colour = "white")) + facet_wrap(~design_construct, scale = "free_y")

grid.arrange(AFB2_single, TIR1_single, TIR1_dual, AFB2_dual, nrow = 2, ncol = 4)
```

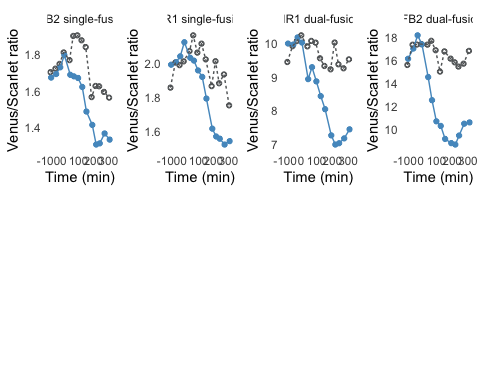

```
W303ratio <- qplot(x = time, y = normalizedratio, data = subset(plate_20210619_W303_sum),
    group = factor(treatment), linetype = factor(treatment), shape = factor(treatment),
    color = factor(treatment)) + xlab("Time (min)") + geom_point(aes(color = treatment),
    size = 0.5) + geom_line(aes(color = treatment), linewidth = 1) + scale_shape_manual(values = c(19,
    1)) + ylab("Venus") + scale_color_manual(values = c("#2E86C1", "#626567")) +
    theme_minimal() + theme(legend.position = "bottom", panel.grid.major = element_line(linewidth = 0.3,
    linetype = "solid", colour = "white"), panel.grid.minor = element_line(linewidth = 0.3,
    linetype = "solid", colour = "white"), axis.line = element_line(colour = "black",
    linewidth = 1, linetype = "solid")) + facet_wrap(~design_construct, scale = "free_y")

W303venus <- qplot(x = time, y = normalized_Greenexpression, data = subset(plate_20210619_W303_sum),
    group = factor(treatment), linetype = factor(treatment), shape = factor(treatment),
    color = factor(treatment)) + xlab("Time (min)") + geom_point(aes(color = treatment),
    size = 0.5) + geom_line(aes(color = treatment), linewidth = 1) + scale_shape_manual(values = c(19,
    1)) + ylab("Venus/Scarlet") + scale_color_manual(values = c("#F1C40F", "#626567")) +
    theme_minimal() + theme(legend.position = "bottom", panel.grid.major = element_line(linewidth = 0.3,
    linetype = "solid", colour = "white"), panel.grid.minor = element_line(linewidth = 0.3,
    linetype = "solid", colour = "white"), axis.line = element_line(colour = "black",
    linewidth = 1, linetype = "solid")) + facet_wrap(~design_construct, scale = "free_y")

grid.arrange(W303venus, W303ratio, nrow = 2, ncol = 2)
```

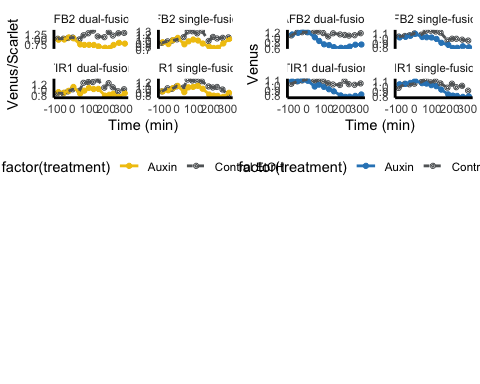

```
Venus_aov <- aov(FL1.Amean ~ design_construct * treatment * before_after, data = plate_20210619_W303_sum %>%
    dplyr::filter(time <= 0 | time >= 200))
summary(Venus_aov)
##                                         Df    Sum Sq   Mean Sq F value   Pr(>F)
## design_construct                         3 5.902e+09 1.967e+09 108.760  < 2e-16
## treatment                                1 9.695e+08 9.695e+08  53.601 5.82e-09
## before_after                             1 4.570e+06 4.570e+06   0.253    0.618
## design_construct:treatment               3 5.293e+07 1.764e+07   0.976    0.414
## design_construct:before_after            3 9.180e+07 3.060e+07   1.692    0.184
## treatment:before_after                   1 6.641e+08 6.641e+08  36.715 3.56e-07
## design_construct:treatment:before_after  3 3.063e+07 1.021e+07   0.564    0.642
## Residuals                               41 7.416e+08 1.809e+07
##
## design_construct                        ***
## treatment                               ***
## before_after
## design_construct:treatment
## design_construct:before_after
## treatment:before_after                  ***
## design_construct:treatment:before_after
## Residuals
## ---
## Signif. codes:  0 '***' 0.001 '**' 0.01 '*' 0.05 '.' 0.1 ' ' 1
(Venus_HSD.test <- agricolae::HSD.test(Venus_aov, trt = c("design_construct", "treatment",
    "before_after"), console = TRUE))
##
## Study: Venus_aov ~ c("design_construct", "treatment", "before_after")
##
## HSD Test for FL1.Amean
##
## Mean Square Error:  18087298
##
## design_construct:treatment:before_after,  means
##
##                                        FL1.Amean      std r      Min      Max
## AFB2 dual-fusion:Auxin:after            24425.97 3887.943 4 19046.09 27855.18
## AFB2 dual-fusion:Auxin:before           34631.51 2164.181 3 33214.07 37122.60
## AFB2 dual-fusion:Control EtOH:after     43943.78 2563.443 4 40314.31 46305.36
## AFB2 dual-fusion:Control EtOH:before    35281.28  747.563 3 34785.81 36141.17
## AFB2 single-fusion:Auxin:after          48152.32 8233.437 4 41566.06 58870.26
## AFB2 single-fusion:Auxin:before         52801.44 3789.368 3 49120.11 56690.32
## AFB2 single-fusion:Control EtOH:after   61495.06 1241.776 4 59958.17 62859.15
## AFB2 single-fusion:Control EtOH:before  53911.90 2286.620 3 51418.04 55909.91
## TIR1 dual-fusion:Auxin:after            41225.93 2818.571 4 37362.85 43733.63
## TIR1 dual-fusion:Auxin:before           46003.40 1595.801 3 44645.69 47761.17
## TIR1 dual-fusion:Control EtOH:after     53108.52 5964.120 4 46969.08 60966.52
## TIR1 dual-fusion:Control EtOH:before    44671.84 1768.004 3 43508.04 46706.33
## TIR1 single-fusion:Auxin:after          53550.99 3388.730 4 50475.84 58381.61
## TIR1 single-fusion:Auxin:before         63962.75 3201.204 3 61073.09 67403.85
## TIR1 single-fusion:Control EtOH:after   65619.10 7089.748 5 53646.79 71126.31
## TIR1 single-fusion:Control EtOH:before  64865.95 3237.883 3 61178.65 67245.10
##
## Alpha: 0.05 ; DF Error: 41
## Critical Value of Studentized Range: 5.154955
##
## Groups according to probability of means differences and alpha level( 0.05 )
##
## Treatments with the same letter are not significantly different.
##
##                                        FL1.Amean groups
## TIR1 single-fusion:Control EtOH:after   65619.10      a
## TIR1 single-fusion:Control EtOH:before  64865.95     ab
## TIR1 single-fusion:Auxin:before         63962.75     ab
## AFB2 single-fusion:Control EtOH:after   61495.06     ab
## AFB2 single-fusion:Control EtOH:before  53911.90     bc
## TIR1 single-fusion:Auxin:after          53550.99     bc
## TIR1 dual-fusion:Control EtOH:after     53108.52     bc
## AFB2 single-fusion:Auxin:before         52801.44    bcd
## AFB2 single-fusion:Auxin:after          48152.32     cd
## TIR1 dual-fusion:Auxin:before           46003.40    cde
## TIR1 dual-fusion:Control EtOH:before    44671.84    cde
## AFB2 dual-fusion:Control EtOH:after     43943.78    cde
## TIR1 dual-fusion:Auxin:after            41225.93     de
## AFB2 dual-fusion:Control EtOH:before    35281.28     ef
## AFB2 dual-fusion:Auxin:before           34631.51     ef
## AFB2 dual-fusion:Auxin:after            24425.97      f
## $statistics
##    MSerror Df     Mean       CV
##   18087298 41 49475.37 8.596028
##
## $parameters
##    test                                  name.t ntr StudentizedRange alpha
##   Tukey design_construct:treatment:before_after  16         5.154955  0.05
##
## $means
##                                        FL1.Amean      std r      Min      Max
## AFB2 dual-fusion:Auxin:after            24425.97 3887.943 4 19046.09 27855.18
## AFB2 dual-fusion:Auxin:before           34631.51 2164.181 3 33214.07 37122.60
## AFB2 dual-fusion:Control EtOH:after     43943.78 2563.443 4 40314.31 46305.36
## AFB2 dual-fusion:Control EtOH:before    35281.28  747.563 3 34785.81 36141.17
## AFB2 single-fusion:Auxin:after          48152.32 8233.437 4 41566.06 58870.26
## AFB2 single-fusion:Auxin:before         52801.44 3789.368 3 49120.11 56690.32
## AFB2 single-fusion:Control EtOH:after   61495.06 1241.776 4 59958.17 62859.15
## AFB2 single-fusion:Control EtOH:before  53911.90 2286.620 3 51418.04 55909.91
## TIR1 dual-fusion:Auxin:after            41225.93 2818.571 4 37362.85 43733.63
## TIR1 dual-fusion:Auxin:before           46003.40 1595.801 3 44645.69 47761.17
## TIR1 dual-fusion:Control EtOH:after     53108.52 5964.120 4 46969.08 60966.52
## TIR1 dual-fusion:Control EtOH:before    44671.84 1768.004 3 43508.04 46706.33
## TIR1 single-fusion:Auxin:after          53550.99 3388.730 4 50475.84 58381.61
## TIR1 single-fusion:Auxin:before         63962.75 3201.204 3 61073.09 67403.85
## TIR1 single-fusion:Control EtOH:after   65619.10 7089.748 5 53646.79 71126.31
## TIR1 single-fusion:Control EtOH:before  64865.95 3237.883 3 61178.65 67245.10
##                                             Q25      Q50      Q75
## AFB2 dual-fusion:Auxin:after           22934.16 25401.31 26893.13
## AFB2 dual-fusion:Auxin:before          33385.97 33557.87 35340.23
## AFB2 dual-fusion:Control EtOH:after    43299.85 44577.73 45221.67
## AFB2 dual-fusion:Control EtOH:before   34851.33 34916.85 35529.01
## AFB2 single-fusion:Auxin:after         41748.87 46086.48 52489.94
## AFB2 single-fusion:Auxin:before        50856.99 52593.87 54642.10
## AFB2 single-fusion:Control EtOH:after  60845.16 61581.46 62231.36
## AFB2 single-fusion:Control EtOH:before 52912.90 54407.76 55158.83
## TIR1 dual-fusion:Auxin:after           40076.43 41903.61 43053.11
## TIR1 dual-fusion:Auxin:before          45124.52 45603.34 46682.25
## TIR1 dual-fusion:Control EtOH:after    49652.60 52249.25 55705.17
## TIR1 dual-fusion:Control EtOH:before   43654.60 43801.16 45253.75
## TIR1 single-fusion:Auxin:after         51941.54 52673.26 54282.71
## TIR1 single-fusion:Auxin:before        62242.20 63411.32 65407.59
## TIR1 single-fusion:Control EtOH:after  64830.36 68765.73 69726.29
## TIR1 single-fusion:Control EtOH:before 63676.38 66174.10 66709.60
##
## $comparison
## NULL
##
## $groups
##                                        FL1.Amean groups
## TIR1 single-fusion:Control EtOH:after   65619.10      a
## TIR1 single-fusion:Control EtOH:before  64865.95     ab
## TIR1 single-fusion:Auxin:before         63962.75     ab
## AFB2 single-fusion:Control EtOH:after   61495.06     ab
## AFB2 single-fusion:Control EtOH:before  53911.90     bc
## TIR1 single-fusion:Auxin:after          53550.99     bc
## TIR1 dual-fusion:Control EtOH:after     53108.52     bc
## AFB2 single-fusion:Auxin:before         52801.44    bcd
## AFB2 single-fusion:Auxin:after          48152.32     cd
## TIR1 dual-fusion:Auxin:before           46003.40    cde
## TIR1 dual-fusion:Control EtOH:before    44671.84    cde
## AFB2 dual-fusion:Control EtOH:after     43943.78    cde
## TIR1 dual-fusion:Auxin:after            41225.93     de
## AFB2 dual-fusion:Control EtOH:before    35281.28     ef
## AFB2 dual-fusion:Auxin:before           34631.51     ef
## AFB2 dual-fusion:Auxin:after            24425.97      f
##
## attr(,"class")
## [1] "group"
Venus_groups <- Venus_HSD.test$groups %>%
    as_tibble(rownames = "names") %>%
    separate(col = names, into = c("design_construct", "treatment", "before_after"),
        sep = "\\:", remove = FALSE) %>%
    left_join(Venus_HSD.test$means %>%
        as_tibble(rownames = "names") %>%
        dplyr::select(c(names, Max)), by = "names") %>%
    mutate(treatment = fct_rev(treatment), before_after = fct_rev(before_after))
mScarlet_aov <- aov(FL4.Amean ~ design_construct * treatment * before_after, data = plate_20210619_W303_sum %>%
    dplyr::filter(time <= 0 | time >= 200))
summary(mScarlet_aov)
##                                         Df    Sum Sq   Mean Sq  F value
## design_construct                         3 1.318e+10 4.393e+09 1152.935
## treatment                                1 2.843e+06 2.843e+06    0.746
## before_after                             1 7.962e+07 7.962e+07   20.894
## design_construct:treatment               3 4.175e+06 1.392e+06    0.365
## design_construct:before_after            3 6.634e+07 2.211e+07    5.803
## treatment:before_after                   1 7.820e+02 7.820e+02    0.000
## design_construct:treatment:before_after  3 7.087e+06 2.362e+06    0.620
## Residuals                               41 1.562e+08 3.811e+06
##                                           Pr(>F)
## design_construct                         < 2e-16 ***
## treatment                                0.39279
## before_after                            4.41e-05 ***
## design_construct:treatment               0.77844
## design_construct:before_after            0.00211 **
## treatment:before_after                   0.98864
## design_construct:treatment:before_after  0.60610
## Residuals
## ---
## Signif. codes:  0 '***' 0.001 '**' 0.01 '*' 0.05 '.' 0.1 ' ' 1
(mScarlet_HSD.test <- agricolae::HSD.test(mScarlet_aov, trt = c("design_construct",
    "treatment", "before_after"), console = TRUE))
##
## Study: mScarlet_aov ~ c("design_construct", "treatment", "before_after")
##
## HSD Test for FL4.Amean
##
## Mean Square Error:  3810672
##
## design_construct:treatment:before_after,  means
##
##                                        FL4.Amean        std r       Min
## AFB2 dual-fusion:Auxin:after            2490.844  198.85812 4  2207.416
## AFB2 dual-fusion:Auxin:before           2026.397   46.66275 3  1973.433
## AFB2 dual-fusion:Control EtOH:after     2759.117  144.66745 4  2612.374
## AFB2 dual-fusion:Control EtOH:before    2110.115  112.55555 3  2013.715
## AFB2 single-fusion:Auxin:after         36197.490 5828.74906 4 31851.266
## AFB2 single-fusion:Auxin:before        31074.988 1904.65789 3 29016.085
## AFB2 single-fusion:Control EtOH:after  38385.588 1153.01803 4 36886.238
## AFB2 single-fusion:Control EtOH:before 31251.821  936.56398 3 30193.587
## TIR1 dual-fusion:Auxin:after            5753.411  381.70386 4  5202.843
## TIR1 dual-fusion:Auxin:before           4582.236  107.55234 3  4483.615
## TIR1 dual-fusion:Control EtOH:after     5556.851  452.28202 4  5075.024
## TIR1 dual-fusion:Control EtOH:before    4557.292  144.44763 3  4390.506
## TIR1 single-fusion:Auxin:after         34660.947 2185.28924 4 33220.282
## TIR1 single-fusion:Auxin:before        31775.292 1187.70866 3 30675.467
## TIR1 single-fusion:Control EtOH:after  34156.499 2380.95848 5 30640.164
## TIR1 single-fusion:Control EtOH:before 33266.440  288.20961 3 32971.733
##                                              Max
## AFB2 dual-fusion:Auxin:after            2666.530
## AFB2 dual-fusion:Auxin:before           2061.453
## AFB2 dual-fusion:Control EtOH:after     2929.881
## AFB2 dual-fusion:Control EtOH:before    2233.807
## AFB2 single-fusion:Auxin:after         44192.014
## AFB2 single-fusion:Auxin:before        32774.074
## AFB2 single-fusion:Control EtOH:after  39681.451
## AFB2 single-fusion:Control EtOH:before 31973.960
## TIR1 dual-fusion:Auxin:after            6084.835
## TIR1 dual-fusion:Auxin:before           4696.914
## TIR1 dual-fusion:Control EtOH:after     6086.477
## TIR1 dual-fusion:Control EtOH:before    4642.111
## TIR1 single-fusion:Auxin:after         37917.098
## TIR1 single-fusion:Auxin:before        33034.740
## TIR1 single-fusion:Control EtOH:after  37060.388
## TIR1 single-fusion:Control EtOH:before 33547.680
##
## Alpha: 0.05 ; DF Error: 41
## Critical Value of Studentized Range: 5.154955
##
## Groups according to probability of means differences and alpha level( 0.05 )
##
## Treatments with the same letter are not significantly different.
##
##                                        FL4.Amean groups
## AFB2 single-fusion:Control EtOH:after  38385.588      a
## AFB2 single-fusion:Auxin:after         36197.490     ab
## TIR1 single-fusion:Auxin:after         34660.947     ab
## TIR1 single-fusion:Control EtOH:after  34156.499     ab
## TIR1 single-fusion:Control EtOH:before 33266.440     ab
## TIR1 single-fusion:Auxin:before        31775.292      b
## AFB2 single-fusion:Control EtOH:before 31251.821      b
## AFB2 single-fusion:Auxin:before        31074.988      b
## TIR1 dual-fusion:Auxin:after            5753.411      c
## TIR1 dual-fusion:Control EtOH:after     5556.851      c
## TIR1 dual-fusion:Auxin:before           4582.236      c
## TIR1 dual-fusion:Control EtOH:before    4557.292      c
## AFB2 dual-fusion:Control EtOH:after     2759.117      c
## AFB2 dual-fusion:Auxin:after            2490.844      c
## AFB2 dual-fusion:Control EtOH:before    2110.115      c
## AFB2 dual-fusion:Auxin:before           2026.397      c
## $statistics
##   MSerror Df    Mean       CV
##   3810672 41 19226.9 10.15293
##
## $parameters
##    test                                  name.t ntr StudentizedRange alpha
##   Tukey design_construct:treatment:before_after  16         5.154955  0.05
##
## $means
##                                        FL4.Amean        std r       Min
## AFB2 dual-fusion:Auxin:after            2490.844  198.85812 4  2207.416
## AFB2 dual-fusion:Auxin:before           2026.397   46.66275 3  1973.433
## AFB2 dual-fusion:Control EtOH:after     2759.117  144.66745 4  2612.374
## AFB2 dual-fusion:Control EtOH:before    2110.115  112.55555 3  2013.715
## AFB2 single-fusion:Auxin:after         36197.490 5828.74906 4 31851.266
## AFB2 single-fusion:Auxin:before        31074.988 1904.65789 3 29016.085
## AFB2 single-fusion:Control EtOH:after  38385.588 1153.01803 4 36886.238
## AFB2 single-fusion:Control EtOH:before 31251.821  936.56398 3 30193.587
## TIR1 dual-fusion:Auxin:after            5753.411  381.70386 4  5202.843
## TIR1 dual-fusion:Auxin:before           4582.236  107.55234 3  4483.615
## TIR1 dual-fusion:Control EtOH:after     5556.851  452.28202 4  5075.024
## TIR1 dual-fusion:Control EtOH:before    4557.292  144.44763 3  4390.506
## TIR1 single-fusion:Auxin:after         34660.947 2185.28924 4 33220.282
## TIR1 single-fusion:Auxin:before        31775.292 1187.70866 3 30675.467
## TIR1 single-fusion:Control EtOH:after  34156.499 2380.95848 5 30640.164
## TIR1 single-fusion:Control EtOH:before 33266.440  288.20961 3 32971.733
##                                              Max       Q25       Q50       Q75
## AFB2 dual-fusion:Auxin:after            2666.530  2438.961  2544.714  2596.597
## AFB2 dual-fusion:Auxin:before           2061.453  2008.869  2044.305  2052.879
## AFB2 dual-fusion:Control EtOH:after     2929.881  2655.805  2747.106  2850.417
## AFB2 dual-fusion:Control EtOH:before    2233.807  2048.269  2082.823  2158.315
## AFB2 single-fusion:Auxin:after         44192.014 31870.387 34373.340 38700.443
## AFB2 single-fusion:Auxin:before        32774.074 30225.445 31434.806 32104.440
## AFB2 single-fusion:Control EtOH:after  39681.451 37980.543 38487.331 38892.376
## AFB2 single-fusion:Control EtOH:before 31973.960 30890.752 31587.918 31780.939
## TIR1 dual-fusion:Auxin:after            6084.835  5692.279  5862.984  5924.116
## TIR1 dual-fusion:Auxin:before           4696.914  4524.898  4566.180  4631.547
## TIR1 dual-fusion:Control EtOH:after     6086.477  5250.871  5532.952  5838.933
## TIR1 dual-fusion:Control EtOH:before    4642.111  4514.882  4639.258  4640.685
## TIR1 single-fusion:Auxin:after         37917.098 33608.882 33753.203 34805.267
## TIR1 single-fusion:Auxin:before        33034.740 31145.568 31615.670 32325.205
## TIR1 single-fusion:Control EtOH:after  37060.388 33524.915 34184.394 35372.637
## TIR1 single-fusion:Control EtOH:before 33547.680 33125.820 33279.906 33413.793
##
## $comparison
## NULL
##
## $groups
##                                        FL4.Amean groups
## AFB2 single-fusion:Control EtOH:after  38385.588      a
## AFB2 single-fusion:Auxin:after         36197.490     ab
## TIR1 single-fusion:Auxin:after         34660.947     ab
## TIR1 single-fusion:Control EtOH:after  34156.499     ab
## TIR1 single-fusion:Control EtOH:before 33266.440     ab
## TIR1 single-fusion:Auxin:before        31775.292      b
## AFB2 single-fusion:Control EtOH:before 31251.821      b
## AFB2 single-fusion:Auxin:before        31074.988      b
## TIR1 dual-fusion:Auxin:after            5753.411      c
## TIR1 dual-fusion:Control EtOH:after     5556.851      c
## TIR1 dual-fusion:Auxin:before           4582.236      c
## TIR1 dual-fusion:Control EtOH:before    4557.292      c
## AFB2 dual-fusion:Control EtOH:after     2759.117      c
## AFB2 dual-fusion:Auxin:after            2490.844      c
## AFB2 dual-fusion:Control EtOH:before    2110.115      c
## AFB2 dual-fusion:Auxin:before           2026.397      c
##
## attr(,"class")
## [1] "group"
mScarlet_groups <- mScarlet_HSD.test$groups %>%
    as_tibble(rownames = "names") %>%
    separate(col = names, into = c("design_construct", "treatment", "before_after"),
        sep = "\\:", remove = FALSE) %>%
    left_join(mScarlet_HSD.test$means %>%
        as_tibble(rownames = "names") %>%
        dplyr::select(c(names, Max)), by = "names") %>%
    mutate(treatment = fct_rev(treatment), before_after = fct_rev(before_after))
boxvenus <- plate_20210619_W303_sum %>%
    dplyr::filter(time <= 0 | time >= 200) %>%
    ggplot(aes(x = fct_rev(before_after), y = FL1.Amean/1000, )) + geom_boxplot(aes(fill = fct_rev(treatment)),
    alpha = 0.7, outlier.shape = NA, show.legend = FALSE) + geom_point(aes(color = fct_rev(treatment)),
    position = position_dodge2(width = 0.55), size = 1) + geom_text(data = Venus_groups,
    mapping = aes(y = 1.05 * Max/1000, label = groups), position = position_dodge2(width = 1),
    color = "black") + scale_fill_manual(values = c("#8f8f8f", "#edc215")) + scale_color_manual(values = c("#8f8f8f",
    "#edc215")) + ylab("Venus") + xlab("50 µM Auxin treatment") + facet_wrap(~fbox,
    scale = "free_y") + facet_grid(~design_construct) + theme_classic() + labs(color = "")


boxmScarlet <- plate_20210619_W303_sum %>%
    dplyr::filter(time <= 0 | time >= 200) %>%
    ggplot(aes(x = fct_rev(before_after), y = FL4.Amean/1000)) + geom_boxplot(aes(fill = fct_rev(treatment)),
    alpha = 0.7, outlier.shape = NA, show.legend = FALSE) + geom_point(aes(color = fct_rev(treatment)),
    position = position_dodge2(width = 0.55), size = 1) + geom_text(data = mScarlet_groups,
    mapping = aes(y = 1.2 * Max/1000, label = groups), position = position_dodge2(width = 1),
    color = "black") + scale_fill_manual(values = c("#8f8f8f", "#EC7063")) + scale_color_manual(values = c("#8f8f8f",
    "#EC7063")) + ylab("mScarlet-I") + xlab("50 µM Auxin treatment") + facet_wrap(~fbox,
    scale = "free_y") + facet_grid(~design_construct, scales = "free_y") + theme_classic() +
    labs(color = "") + scale_y_log10()


guide_area()/boxvenus/boxmScarlet + plot_annotation(tag_levels = "A") + plot_layout(guides = "collect",
    heights = c(0.3, 1, 1)) & theme(legend.box = "horizontal", legend.spacing = unit(0,
    "pt"), legend.justification = "top", legend.margin = margin(), legend.box.spacing = unit(0,
    "pt"), legend.background = element_blank(), legend.box.background = element_blank(),
    legend.title = element_blank(), legend.key = element_blank())
```

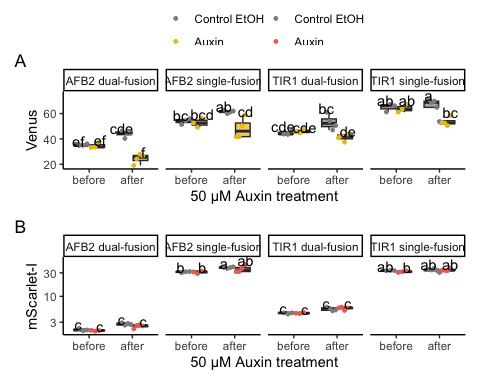

```

ggsave("relative-expression-box.pdf", width = 6, height = 5)
ggsave("relative-expression-box.png", width = 6, height = 5)
plate_20210619_read3and12 <- read.flowSet(path = "~/Google Drive/Shared drives/PlantSynBioLab/Pat/Experiments/Time course assays/20210619/W303only_read3and12/",
    alter.names = TRUE)
annotation <- createAnnotation(yourFlowSet = plate_20210619_read3and12)
write.csv(annotation, "/Users/patchaisupa/Google Drive/Shared drives/PlantSynBioLab/Pat/Experiments/Time course assays/20210619/20210619_W303only_read3and12_annotation.csv")
annotation <- read.csv("~/Google Drive/Shared drives/PlantSynBioLab/Pat/Experiments/Time course assays/20210619/20210619_W303only_read3and12_annotation.csv")
aplate_20210619_read3and12 <- annotateFlowSet(yourFlowSet = plate_20210619_read3and12,
    annotation_df = annotation, mergeBy = "name")
head(rownames(pData(aplate_20210619_read3and12)))
## [1] "D01.fcs" "D02.fcs" "D03.fcs" "D04.fcs" "D07.fcs" "D08.fcs"
head(pData(aplate_20210619_read3and12))
##            name X strain    treatment reading before_after design_construct
## D01.fcs D01.fcs 1 yWL161 Control EtOH       3       before       TIR1 trans
## D02.fcs D02.fcs 2 yWL162 Control EtOH       3       before       AFB2 trans
## D03.fcs D03.fcs 3 yWL185 Control EtOH       3       before         TIR1 cis
## D04.fcs D04.fcs 4 yWL186 Control EtOH       3       before         AFB2 cis
## D07.fcs D07.fcs 5 yWL161        Auxin       3       before       TIR1 trans
## D08.fcs D08.fcs 6 yWL162        Auxin       3       before       AFB2 trans
W303_read3and12 <- summarizeFlow(aplate_20210619_read3and12, gated = TRUE)
## [1] "Summarizing all events..."
W303_read3and12 <- W303_read3and12 %>%
    mutate(design = str_extract(design_construct, "(?<=\\s).*")) %>%
    mutate(design = str_extract(design_construct, ".*(?=\\s)"))

venus3and12_unnorm <- ggplot(W303_read3and12, aes(x = design_construct, y = FL1.Amean,
    alpha = fct_rev(before_after), group = treatment)) + geom_point(aes(colour = factor(treatment),
    shape = factor(treatment)), size = 2, position = position_jitterdodge(dodge.width = 0.7,
    jitter.width = 0.5)) + scale_color_manual(values = c("#F1C40F", "#5F6A6A")) +
    ylab("Venus") + theme_classic() + theme(axis.text.x = element_text(angle = 45,
    hjust = 1)) + scale_alpha_manual(values = c(0.3, 0.8))

mScarlet3and12_unnorm <- ggplot(W303_read3and12, aes(x = design_construct, y = FL4.Amean,
    alpha = fct_rev(before_after), group = treatment)) + geom_point(aes(colour = factor(treatment),
    shape = factor(treatment)), size = 2, position = position_jitterdodge(dodge.width = 0.7,
    jitter.width = 0.5)) + scale_color_manual(values = c("#E74C3C", "#5F6A6A")) +
    ylab("mScarlet-I") + theme_classic() + theme(axis.text.x = element_text(angle = 45,
    hjust = 1)) + scale_alpha_manual(values = c(0.3, 0.8))

guide_area()/(venus3and12_unnorm | mScarlet3and12_unnorm) + plot_layout(guides = "collect",
    heights = c(1, 3)) + plot_annotation(tag_levels = "A") & theme(legend.position = "top",
    legend.direction = "vertical", legend.title = element_blank(), axis.title.x = element_blank(),
    legend.justification = "left", legend.box.just = "left", legend.margin = margin())
```

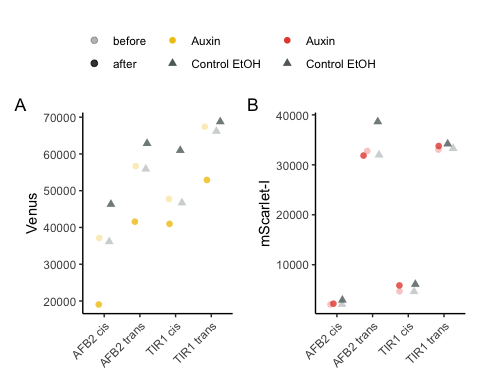

ggsave("relative-expression.png", width = 4, height = 3)
ggsave("relative-expression.pdf", width = 4, height = 3)

Coefficient of variation (CV) analysis

```

data <- steadyState(aplate_20210619_W303, gated = TRUE)
## [1] "No further gating applied."
## [1] "Converting events..."
# data <- tidyFlow(aplate_20210619_W303)

data <- subset(data, strain == "yWL161" & name %in% c("11L01.fcs", "11L07.fcs"))

# range(data$FL1.A) sd(data$FLratio)/mean(data$FLratio) range(data$FLratio)
data <- subset(data, FL1.A > 1 & FL4.A > 1)
data$FLratio <- data$FL1.A/data$FL4.A
range(data$FL1.A)
## [1]     722 2579491
# calculate cvs
sd(data$FLratio)/mean(data$FLratio)
## [1] 1.821223
range(data$FLratio)
## [1]   0.06052985 488.44186047
cv <- function(x) return(round(sd(x)/mean(x), 2))

# calculate normalized values
data$Venus <- data$FL1.A/median(data$FL1.A)
data$mScarlet <- data$FL4.A/median(data$FL4.A)
data$ratio <- data$FLratio/median(data$FLratio)
CVs <- data %>%
    group_by(treatment) %>%
    summarise(across(where(is_double), cv))
# make a tidy, long dataset

data_long <- data %>%
    dplyr::select(treatment, Venus, mScarlet, ratio) %>%
    pivot_longer(cols = c(Venus, mScarlet, ratio), names_to = "parameter", values_to = "value")
# need to also format CVs appropriately for annotating
CVs <- CVs %>%
    dplyr::select(treatment, Venus, mScarlet, ratio) %>%
    pivot_longer(cols = c(Venus, mScarlet, ratio), names_to = "parameter", values_to = "value")

# data
CV_plot <- ggplot(data = data_long, mapping = aes(x = value, color = treatment)) +
    geom_density() + xlim(c(-1, 4)) + labs(x = "median normalized intensity", color = "treatment") +
    facet_grid(parameter ~ .) + theme_test() + geom_text(data = subset(CVs, treatment ==
    "Auxin"), aes(label = paste0("CV = ", value)), x = 0.1, y = 0.6) + geom_text(data = subset(CVs,
    treatment == "Control EtOH"), aes(label = paste0("CV = ", value)), x = 1.8, y = 0.6) +
    scale_color_viridis_d(option = "D", end = 0.75, direction = -1)
CV_plot
```

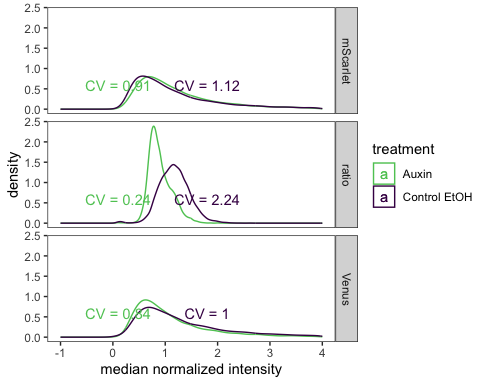

### Auxin prodcution in stationary phase

Read in annotated flowSet

```
flowSet <- read.flowSet(path = paste0("~/Google Drive/Shared drives/PlantSynBioLab/Auxin Biosensor Manuscript/Auxin Biosensor Data/FlowSets/",
    experiment_date, "_", experiment_name), phenoData = "annotation.txt")
write.flowSet(flowSet, "flowSets/auxin-biosynthesis")
flowSet <- read.flowSet(path = "flowSets/auxin-biosynthesis/", phenoData = "annotation.txt")
```

#### Summary Analysis

```
load("PSB_Accuri_W303.RData")
dat_sum <- summarizeFlow(flowSet, ploidy = "haploid", only = "yeast")  # These gates might not work well for YPH499, but there was a lot of debris
## [1] "Gating with haploid yeast gate..."
## [1] "Summarizing all yeast events..."
dat_sum <- mutate(.data = dat_sum, across(starts_with("FL"), as.numeric)) %>%
    mutate(strain = str_remove(strain, " ratiometric sensor") %>%
        str_replace(pattern = "in trans", replacement = "single-fusion") %>%
        str_replace(pattern = "in cis", replacement = "dual-fusion"))
dat_sum$pred_auxin <- dat_sum$FL4.Amean/dat_sum$FL1.Amean
dat_sum$pred_auxin_SE <- with(dat_sum, sqrt((FL4.Asd/100/FL4.Amean)^2 + (FL1.Asd/100/FL1.Amean)^2) *
    pred_auxin)
dat_sum$pred_auxin_sd <- with(dat_sum, sqrt((FL4.Asd/FL4.Amean)^2 + (FL1.Asd/FL1.Amean)^2) *
    pred_auxin)
ggplot(data = dat_sum, aes(x = fct_reorder(treat, pred_auxin), y = pred_auxin, color = factor(strain))) +
    geom_point() + theme(axis.text.x = element_text(angle = 45, hjust = 1, vjust = 1)) +
    facet_grid(. ~ strain, scales = "free_x") + coord_flip() + theme(legend.position = "bottom",
    legend.direction = "vertical") + labs(x = "", y = "Predicted auxin (mScarlet-I/Venus-IAA17)",
    color = "strain")
```

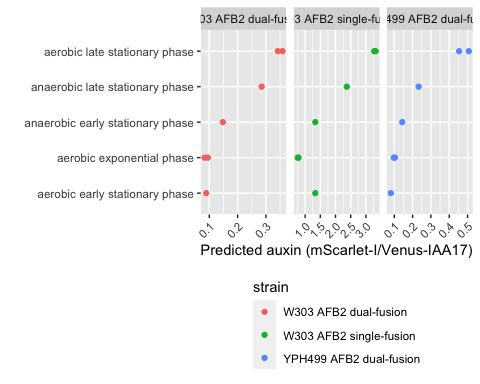

```
ggsave("strain_sensor_comparison_raw.pdf", width = 8, height = 3)
dat_sum <- dat_sum %>%
    group_by(strain) %>%
    mutate(norm_pred_auxin = pred_auxin/mean(pred_auxin[which(.data$treat == "aerobic exponential phase")]))

ggplot(data = dat_sum, aes(x = fct_reorder(treat, norm_pred_auxin), y = norm_pred_auxin,
    color = factor(strain))) + geom_point(position = position_jitter(width = 0.2)) +
    theme(axis.text.x = element_text(angle = 45, hjust = 1, vjust = 1)) + coord_flip() +
    theme(legend.position = "top", legend.direction = "vertical") + labs(x = "",
    y = "Normalized Predicted auxin (mScarlet-I/Venus-IAA17)", color = "strain")
```

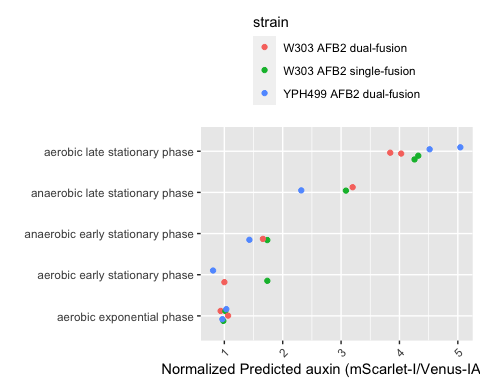

ggsave("strain_sensor_comparison_norm.pdf", width = 5, height = 5)

Full distributions

```
data <- steadyState(flowset = flowSet, ploidy = "haploid", only = "yeast")
## [1] "Gating with haploid yeast gate..."
## [1] "Converting events..."

hist(x = log(data$FL1.A, 10))
```

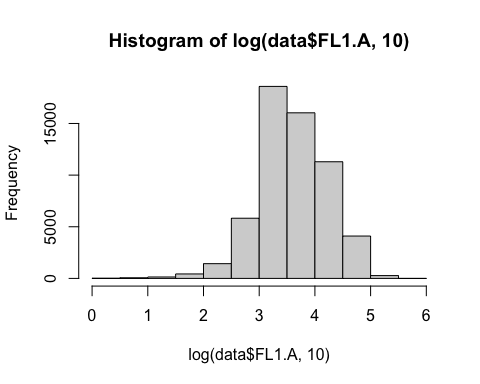

hist(x = log(data$FL4.A, 10))

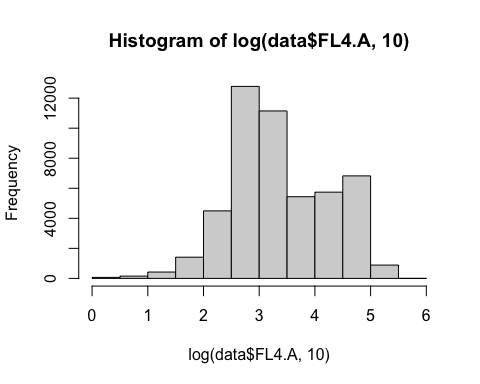

```
data$pred_auxin <- data$FL4.A/data$FL1.A
data <- data %>%
    dplyr::filter(FL1.A > 1 & FL4.A > 1)
data$treat <- data$treat %>%
    str_remove("phase")
endog_auxin <- ggplot(data = subset(data, strain == "W303 ratiometric sensor AFB2 in cis" &
    pred_auxin < 2), aes(x = pred_auxin, y = fct_reorder(treat, .x = pred_auxin,
    .fun = median, .desc = TRUE) %>%
    fct_relevel("aerobic exponential ", after = Inf), fill = factor(stat(quantile)))) +
    stat_density_ridges(geom = "density_ridges_gradient", calc_ecdf = TRUE, quantiles = 4,
        color = "grey", alpha = 0.5) + scale_fill_viridis_d(name = "Quartiles") +
    theme(legend.position = NULL) + #facet_wrap(.~treat) + theme(legend.position
    theme(legend.position = NULL) + #facet_wrap(.~treat) + = NULL) +
    theme(legend.position = NULL) + #facet_wrap(.~treat) + #facet_wrap(.~treat)
    theme(legend.position = NULL) + #facet_wrap(.~treat) + +
xlim(c(-0.1, 1)) + labs(y = NULL, x = "Predicted relative auxin production (AU) \n (AFB2-mScarlet-I/Venus-IAA17)") +
    theme_ridges() + theme(legend.position = "none")
ggsave("20210626_auxin_production_quartiles.pdf", width = 5, height = 4)
endog_auxin
```

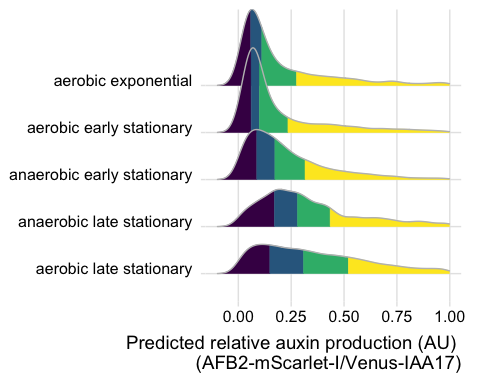

#### Biosensor sensitivity at stationary phase

```
plate_09282022 <- read.plateSet(path = "~/Google Drive/Shared drives/PlantSynBioLab/Pat/Experiments/Does-response assay/09282022_yWL210_DRA-stationary/Data/",
    pattern = "S-DRA*")
annotation_09282022 <- read.csv("~/Google Drive/Shared drives/PlantSynBioLab/Pat/Experiments/Does-response assay/09282022_yWL210_DRA-stationary/09282022_stationaryphase_annotation.csv")

aplate_09282022 <- annotateFlowSet(yourFlowSet = plate_09282022, annotation_df = annotation_09282022,
    mergeBy = "name")
head(rownames(pData(aplate_09282022)))
head(pData(aplate_09282022))
write.flowSet(aplate_09282022, "flowSets/AFB2-dual-stationary")
aplate_09282022 <- read.flowSet(path = "flowSets/AFB2-dual-stationary/", phenoData = "annotation.txt")

dat_sumGr_09282022 <- summarizeFlow(aplate_09282022, gated = TRUE)
## [1] "Summarizing all events..."

model.LL4_09282022 <- drm(YL1.Amean/BL1.Amean ~ dose, data = subset(dat_sumGr_09282022,
    set == "4"), fct = LL.4(names = c("Slope", "Lower Limit", "Upper Limit", "ED50")))

pm210stat <- expand.grid(treatment = exp(seq(log(1e-05), log(100), length = 1000)))
pm210stat <- cbind(pm210stat, predict(model.LL4_09282022, newdata = pm210stat, interval = "confidence"))  #m2all = model of 210 data

plot210_stat <- ggplot(subset(dat_sumGr_09282022, set == "4"), aes(x = dose, y = YL1.Amean/BL1.Amean)) +
    geom_ribbon(data = pm210stat, aes(x = treatment, y = Prediction, ymin = Lower,
        ymax = Upper), alpha = 0.2) + geom_line(data = pm210stat, aes(x = treatment,
    y = Prediction)) + ylab("AFB2-mScarlet-I/Venus-IAA17") + xlab("Exogenous Auxin (µM)") +
    scale_x_log10(labels = scales::label_number(drop0trailing = TRUE)) + scale_color_viridis_d() +
    geom_point() + theme_classic(base_size = 14) + theme(legend.position = "none")
plot210_stat
```

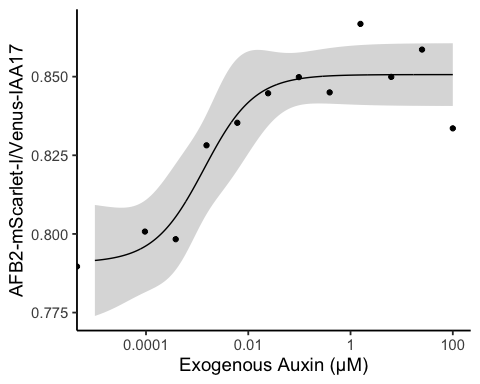

#### Combined figure

endog_auxin + plot210_stat + plot_annotation(tag_levels = "A")

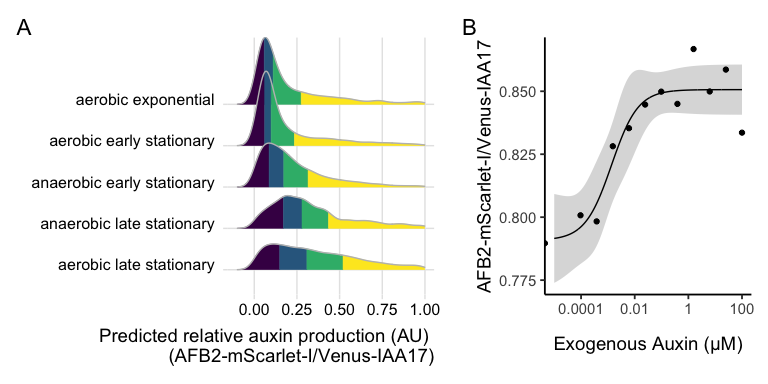

ggsave("auxin-accumulation.pdf", width = 8, height = 4)
ggsave("auxin-accumulation.png", width = 8, height = 4)

### Mutant library analysis

Here we aim to test auxin induced Venus-IAA17 degradation relative to the bicistronic mScarlet-I control with AFB2 expressed from the p1 plasmid in the OrthoRep continuous mutagenesis system.

#### Procedure

Temperature 30 degree celcius Shaking at 250rpm DO-URA-HIS, starting volume: 10 mL Overnight concentration: 30 events/uL 1PM started from 2 colonies

Initial reading ~10AM Auxin concentration: 100 uM (.1% DMSO) After 4th reading: B04.fcs

###Importing and annotating data

```
aplate1 <- read.flowSet(path = "flowSets/OrthoRep", phenoData = "annotation.txt")
dat_sum <- summarizeFlow(aplate1, gated = TRUE)
## [1] "Summarizing all events..."
data <- flowTime::tidyFlow(aplate1, gated = TRUE)
## [1] "No further gating applied."
## [1] "Converting events..."
data <- subset(data, FL4.A > 1)
range(data$FL1.A)
## [1]        0 15867629
range(data$FL4.A)
## [1]       6 7690137
data$ratio <- data$FL1.A/data$FL4.A
data <- subset(data, data$ratio < 10)
```

Make a kernel density plot overlapping mutant population with parent, DMSO and IAA that we can then stack a few timepoints with facets. For timepoints it looks like the 2nd, 7th and 12th would be a good demonstration of the timecourse.

```
wells <- dat_sum %>%
    # Get well/file names of the 2nd, 7th and 12th readings of the 4 strains.
dplyr::slice(1:4 + unlist(lapply((c(2, 7, 12) - 1) * 4, rep, 4))) %>%
    dplyr::pull("name")
data <- dplyr::filter(data, name %in% wells)

data$approxtime <- signif(data$etime, digits = 1)
signif(unique(data$approxtime), 1)
## [1]  30 200 600
data$approxtime <- as.factor(data$approxtime) %>%
    fct_recode(`0 hour` = "30", `1 hour` = "200", `6 hour` = "600")
data$strain <- fct_recode(data$strain, `high fidelity` = "wild", `error prone` = "mutant")
data <- data %>%
    mutate(Venus = FL1.A, mScarlet = FL4.A, ratio = Venus/mScarlet)

cv <- function(x) return(round(sd(x)/mean(x), 2))
CVs <- data %>%
    group_by(treatment, strain) %>%
    dplyr::summarise(across(where(is_double), cv))
CVs <- CVs %>%
    dplyr::select(treatment, Venus, mScarlet, ratio) %>%
    pivot_longer(cols = c(Venus, mScarlet, ratio), names_to = "parameter", values_to = "value")

kernel_plot <- ggplot(data = data, mapping = aes(x = ratio, color = treatment, linetype = strain)) +
    geom_density() + coord_cartesian(x = c(0.5, 2)) + facet_grid(approxtime ~ .) +
    theme_test() + scale_color_viridis_d(option = "D", end = 0.75, direction = -1) +
    labs(x = "Venus-IAA17/mScarlet ratio", linetype = "polymerase")
kernel_plot
```

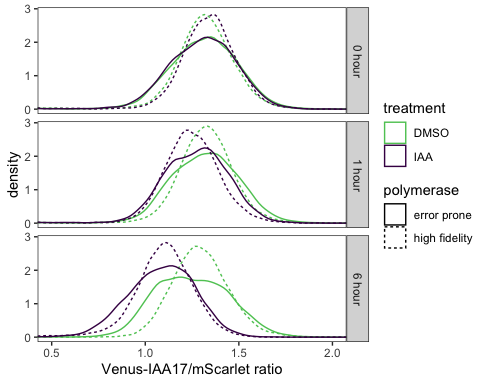

ggsave("orthorep_degradation.pdf", height = 4, width = 4)
ggsave("orthorep_degradation.png", height = 4, width = 4)

#### Gating Strategy

data <- aplate1[wells]

To gate out the high FSC-A debris we will use only the lower 99.5% of the data

g <- gate_quantile(fr = data[[1]], channel = "FSC.A", probs = 0.995)
autoplot(data[[1]], x = "FSC-A") + geom_gate(g)

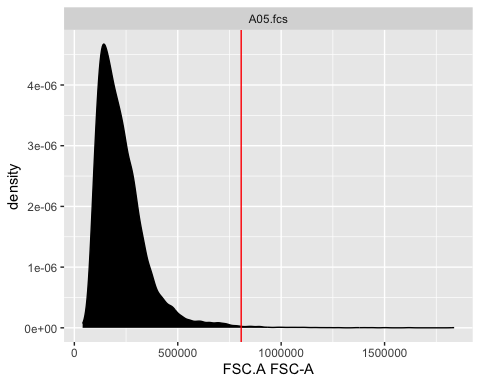

```
Subset(data[[1]], !g)
## flowFrame object 'A05.fcs'
## with 9950 cells and 14 observables:
##        name   desc     range  minRange  maxRange
## $P1   FSC.A  FSC-A  16777216         0  16777216
## $P2   SSC.A  SSC-A  16777216         0  16777216
## $P3   FL1.A  FL1-A  16777216         0  16777216
## $P4   FL2.A  FL2-A  16777216         0  16777216
## $P5   FL3.A  FL3-A  16777216         0  16777216
## ...     ...    ...       ...       ...       ...
## $P10  FL2.H  FL2-H  16777216         0  16777216
## $P11  FL3.H  FL3-H  16777216         0  16777216
## $P12  FL4.H  FL4-H  16777216         0  16777216
## $P13  Width  Width  16777216         0  16777216
## $P14   Time   Time  16777216         0  16777216
## 161 keywords are stored in the 'description' slot
autoplot(Subset(data[[3]], !g), x = "FSC-A", "SSC-A")
```

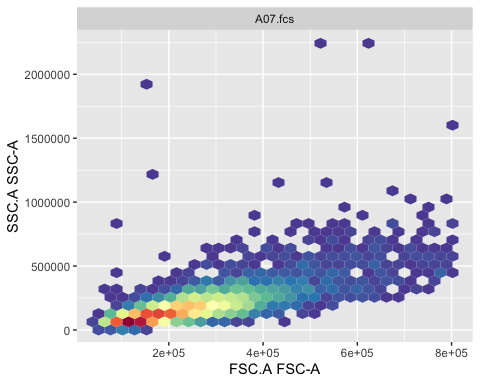

autoplot(data[[3]], "FSC-A", "SSC-A") + geom_gate(g)

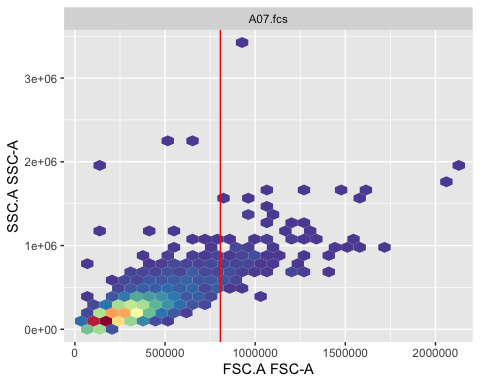
 Now we need to gate out only singlet cells

```
chnl <- c("FSC-A", "FSC-H")
singlets <- gate_singlet(x = Subset(data[[1]], !g), area = "FSC.A", height = "FSC.H",
    prediction_level = 0.999, maxit = 20)
autoplot(data[[1]], "FSC-A", "FSC-H") + geom_gate(singlets)
```

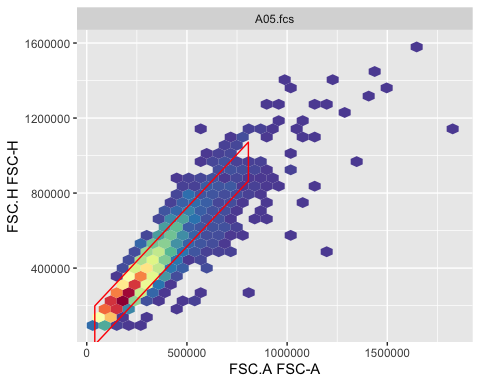
 Now we can assess this singlets gate over for several frames

```
length(data)
## [1] 12
autoplot(data, x = "FSC-A", y = "FSC-H") + geom_gate(singlets) + facet_wrap("name",
    ncol = 4)
```

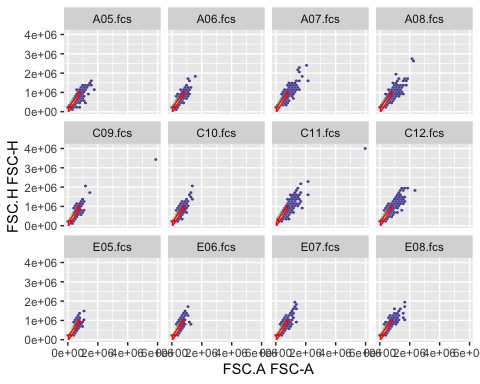

```

autoplot(Subset(data, !g) %>%
    Subset(singlets), x = "FL1-A") + facet_wrap("name", ncol = 4)
```

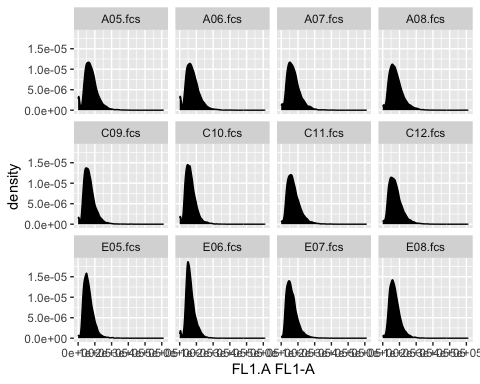

```
autoplot(Subset(data, !g | singlets), x = "FSC-A", y = "SSC-A") + facet_wrap("name",
    ncol = 4)
```

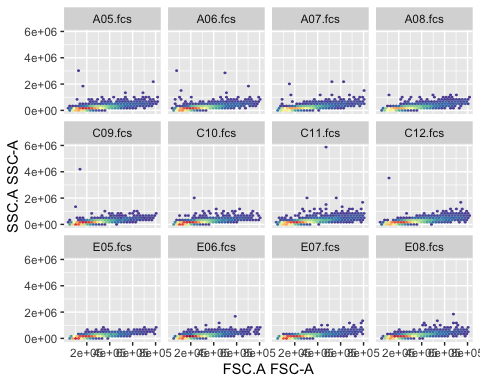

This looks very consistent across the course of this experiment. We can summarize this gating across the whole experiment, but will not show this here.

```
# summary(filter(data, !g & singlets))
data <- Subset(data, !g & singlets)
```

Plot Fluorescence vs. Time

```
dat_sum <- summarizeFlow(data, gated = TRUE)
## [1] "Summarizing all events..."

ggplot(data = dat_sum, aes(x = time, y = FL1.Amean/FL4.Amean, color = factor(strain),
    linetype = factor(treatment))) + geom_line() + xlab("Time post first reading (min)") +
    ylab("Reporter Fluorescence, FL1 (AU)") + geom_text(aes(label = name))
```

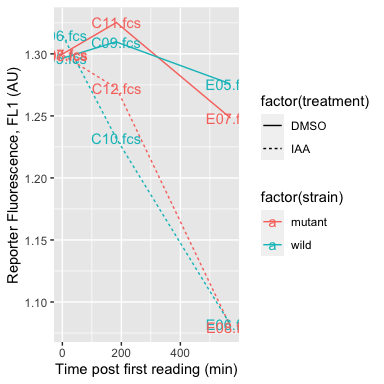

```

ggplot(data = dat_sum, aes(x = time, y = conc, color = factor(strain), linetype = factor(treatment))) +
    geom_line() + xlab("Time post first reading (min)") + ylab("Reporter Fluorescence, FL1 (AU)")
```

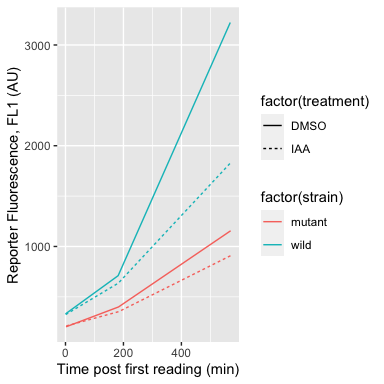
 Define a gate containing ~95% of the untreated cells expressing the wildtype polymerase.

```
logt <- estimateLogicle(data[["E05.fcs"]], channels = c("FL4.A", "FL1.A"))
data <- transform(data[c("E05.fcs", "E06.fcs", "E07.fcs", "E08.fcs")], logt)
untreated <- gate_flowclust_2d(data[["E05.fcs"]], xChannel = "FL4.A", yChannel = "FL1.A",
    K = 1, quantile = 0.9, filterId = "untreated")
treated <- gate_flowclust_2d(data[["E06.fcs"]], xChannel = "FL4.A", yChannel = "FL1.A",
    K = 1, quantile = 0.9, filterId = "treated")


ggcyto(data, aes(x = "FL4.A", y = "FL1.A")) + geom_point(color = "green4", alpha = 0.1,
    size = 0.5) + ggthemes::theme_base() + labs(x = "mScarlet-I (free FP)", y = "Venus-IAA17") +
    facet_grid(fct_rev(strain) %>%
        fct_recode(`wild-type` = "wild") ~ treatment) + coord_cartesian(xlim = c(1.4,
    2.6), ylim = c(1.4, 2.6)) + geom_gate(untreated, colour = "magenta", linetype = 2) +
    geom_gate(treated, colour = "magenta") + geom_stats(adjust = c(0.1, 0.05), size = 4.5,
    alpha = 0.5)
```

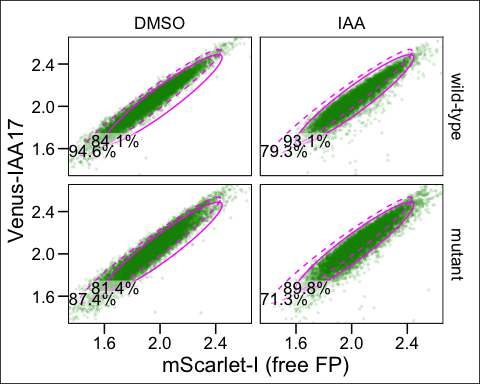

ggsave("sorting-strategy.pdf", height = 4.5, width = 6)
ggsave("sorting-strategy.png", height = 4.5, width = 6)

#### Mutation frequency calculation

As an overestimate, these cells were cultured for ~12 generations over 24 hours. So this results in 2^12^ cells for each initial.

AFB2 is 1725 bps, and the mutation rate of the error prone polymerase is calculated to be $1\times{10}^{-5}$ substitutions per base. We will assume there is only 1 copy of the p1 plasmid per cell. We will also assume withing this coding sequence, for every 2.1 substitutions (sub) there is 1 nonsynonymous substitution (nsub), per the average across the codon table, not factoring in codon usage across TIR1.

$$\begin{matrix} 1\times{10}^{-5}\text{ sub} & 1725\text{ bases}*2\text{ (bp)} & 1\text{ nsub} \\ \text{base} & \text{cell} & 2.1\text{ sub} \end{matrix}=0.02\frac{\text{nsub}}{\text{cell}}$$

So on the low end, ~0.0164286 percent of cells have a nonsynonymous substitution in *TIR1*. But because this substitution rate is really compounding over 12 generations, this estimate is quite low.

Based on this baseline rate per cell (or generation, cell duplication) $r$, we can then compound this over $t$ generations, to find the compounded rate $r_{c}$.

$$r_{c}=\left( 1+r \right)^{t}-1=\left( 1+0.0164286 \right)^{12}-1=0.2159686$$

And on the high end, which is a more accurate measure, ~22% of our population contains a nonsynonymous substitution in *TIR1*.

### Session Info

```
sessionInfo()
## R Under development (unstable) (2022-10-30 r83209)
## Platform: aarch64-apple-darwin20 (64-bit)
## Running under: macOS Ventura 13.2.1
##
## Matrix products: default
## BLAS:   /Library/Frameworks/R.framework/Versions/4.3-arm64/Resources/lib/libRblas.0.dylib
## LAPACK: /Library/Frameworks/R.framework/Versions/4.3-arm64/Resources/lib/libRlapack.dylib
##
## locale:
## [1] en_US.UTF-8/en_US.UTF-8/en_US.UTF-8/C/en_US.UTF-8/en_US.UTF-8
##
## attached base packages:
## [1] stats     graphics  grDevices utils     datasets  methods   base
##
## other attached packages:
##  [1] agricolae_1.3-5      ggthemes_4.2.4       patchwork_1.1.2
##  [4] wesanderson_0.3.6    flowClust_3.37.0     flowStats_4.11.0
##  [7] ggcyto_1.27.4        flowWorkspace_4.11.0 ncdfFlow_2.45.0
## [10] BH_1.78.0-0          openCyto_2.11.1      gridExtra_2.3
## [13] drc_3.0-1            MASS_7.3-58.1        lubridate_1.9.2
## [16] forcats_1.0.0        purrr_1.0.1          readr_2.1.4
## [19] tibble_3.1.8         tidyverse_2.0.0      tidyr_1.3.0
## [22] dplyr_1.1.0          ggridges_0.5.4       stringr_1.5.0
## [25] ggplot2_3.4.1        flowTime_1.23.1      flowCore_2.11.0
##
## loaded via a namespace (and not attached):
##   [1] RColorBrewer_1.1-3  rstudioapi_0.14     magrittr_2.0.3
##   [4] TH.data_1.1-1       rainbow_3.7         farver_2.1.1
##   [7] rmarkdown_2.19      ragg_1.2.5          zlibbioc_1.45.0
##  [10] vctrs_0.5.2         RCurl_1.98-1.9      htmltools_0.5.4
##  [13] haven_2.5.2         plotrix_3.8-2       deSolve_1.34
##  [16] hdrcde_3.4          pracma_2.4.2        KernSmooth_2.23-20
##  [19] plyr_1.8.8          sandwich_3.0-2      zoo_1.8-11
##  [22] mime_0.12           lifecycle_1.0.3     pkgconfig_2.0.3
##  [25] Matrix_1.5-3        R6_2.5.1            fastmap_1.1.0
##  [28] shiny_1.7.4         digest_0.6.31       klaR_1.7-1
##  [31] colorspace_2.0-3    S4Vectors_0.37.3    textshaping_0.3.6
##  [34] labeling_0.4.2      cytolib_2.11.0      fansi_1.0.3
##  [37] timechange_0.2.0    abind_1.4-5         compiler_4.3.0
##  [40] withr_2.5.0         carData_3.0-5       DBI_1.1.3
##  [43] highr_0.9           hexbin_1.28.2       corpcor_1.6.10
##  [46] gtools_3.9.4        tools_4.3.0         rrcov_1.7-2
##  [49] httpuv_1.6.9        glue_1.6.2          IDPmisc_1.1.20
##  [52] questionr_0.7.8     nlme_3.1-161        promises_1.2.0.1
##  [55] grid_4.3.0          cluster_2.1.4       generics_0.1.3
##  [58] gtable_0.3.1        fda_6.0.5           labelled_2.10.0
##  [61] tzdb_0.3.0          data.table_1.14.6   hms_1.1.2
##  [64] car_3.1-1           utf8_1.2.2          BiocGenerics_0.45.0
##  [67] pillar_1.8.1        later_1.3.0         robustbase_0.95-0
##  [70] splines_4.3.0       lattice_0.20-45     AlgDesign_1.2.1
##  [73] survival_3.4-0      deldir_1.0-6        ks_1.14.0
##  [76] RProtoBufLib_2.11.0 tidyselect_1.2.0    RBGL_1.75.0
##  [79] fds_1.8             miniUI_0.1.1.1      knitr_1.41
##  [82] stats4_4.3.0        xfun_0.35           Biobase_2.59.0
##  [85] matrixStats_0.63.0  DEoptimR_1.0-11     stringi_1.7.8
##  [88] yaml_2.3.6          evaluate_0.19       codetools_0.2-18
##  [91] interp_1.1-3        Rgraphviz_2.43.0    graph_1.77.1
##  [94] cli_3.6.0           flowViz_1.63.0      systemfonts_1.0.4
##  [97] xtable_1.8-4        munsell_0.5.0       Rcpp_1.0.9
## [100] png_0.1-8           XML_3.99-0.13       parallel_4.3.0
## [103] ellipsis_0.3.2      mclust_6.0.0        latticeExtra_0.6-30
## [106] jpeg_0.1-10         bitops_1.0-7        viridisLite_0.4.1
## [109] mvtnorm_1.1-3       scales_1.2.1        pcaPP_2.0-3
## [112] combinat_0.0-8      rlang_1.0.6         formatR_1.14
## [115] multcomp_1.4-22     mnormt_2.1.1
```

1. Biological Systems Engineering, Virginia Tech, Blacksburg, VA , USA, co-first author
2. Biological Systems Engineering, Virginia Tech, Blacksburg, VA, USA, co-first author
3. Fralin Life Sciences Institute, Virginia Tech, Blacksburg, VA, USA
4. Biological Systems Engineering, Virginia Tech, Blacksburg, VA, USA
5. Biological Systems Engineering, Virginia Tech, Blacksburg, VA, USA
6. Biochemistry and Fralin Life Sciences Institute, Virginia Tech, Blacksburg, VA, USA
7. Biological Systems Engineering, Fralin Life Sciences Institute, and Translational Plant Sciences Center, Virginia Tech, Blacksburg, VA, USA,
